# RNA-seq gene expression profiling of the bladder in a mouse model of urogenital schistosomiasis

**DOI:** 10.1101/2024.06.29.601185

**Authors:** Kenji Ishida, Derick N. M. Osakunor, Mario Rossi, Olivia K. Lamanna, Evaristus C. Mbanefo, James J. Cody, Loc Le, Michael H. Hsieh

**Affiliations:** Sheikh Zayed Institute for Pediatric Surgical Innovation, Children’s National Research Institute, Washington, District of Columbia, United States; Department of Biotechnological and Applied Clinical Sciences, University of L’Aquila, L’Aquila, Italy; National Eye Institute, National Institutes of Health, Bethesda, Maryland, United States; Charles River Laboratories, Rockville, Maryland, United States; National Cancer Institute, National Institutes of Health, Bethesda, Maryland, United States; Department of Urology, The George Washington University, Washington, District of Columbia, United States

## Abstract

**Background:** Parasitic flatworms of the *Schistosoma* genus cause schistosomiasis, which affects over 230 million people. *Schistosoma haematobium* causes the urogenital form of schistosomiasis (UGS), which can lead to hematuria, fibrosis, and increased risk of secondary infections by bacteria or viruses. UGS is also linked to bladder cancer. To understand the bladder pathology during *S. haematobium* infection, our group previously developed a mouse model that involves the injection of *S. haematobium* eggs into the bladder wall. Using this model, we studied changes in epigenetics profile, as well as changes in gene and protein expression in the host bladder tissues. In the current study, we expand upon this work by examining the expression level of both host and parasite genes using RNA sequencing (RNA-seq) in the mouse bladder wall injection model of *S. haematobium* infection.

**Methods:** We used a mouse model of *S. haematobium* infection in which parasite eggs or vehicle control were injected into the bladder walls of female BALB/c mice. RNA-seq was performed on the RNA isolated from the bladders four days after bladder wall injection.

**Results/Conclusions:** RNA-seq analysis of egg- and vehicle control-injected bladders revealed the differential expression of 1025 mouse genes in the egg-injected bladders, including genes associated with cellular infiltration, immune cell chemotaxis, cytokine signaling, and inflammation We also observed the upregulation of immune checkpoint-related genes, which suggests that while the infection causes an inflammatory response, it also dampens the response to avoid excessive inflammation-related damage to the host. Identifying these changes in host signaling and immune responses improves our understanding of the infection and how it may contribute to the development of bladder cancer. Analysis of the differential gene expression of the parasite eggs between bladder-injected versus uninjected eggs revealed 119 *S. haematobium* genes associated with transcription, intracellular signaling, and metabolism. The analysis of the parasite genes also revealed fewer transcript reads compared to that found in the analysis of mouse genes, highlighting the challenges of studying parasite egg biology in the mouse model of *S. haematobium* infection.

**Author summary:** More than 230 million people worldwide are estimated to carry infection with parasites belonging to the *Schistosoma* genus, which cause morbidity associated with parasite egg deposition. Praziquantel, the drug of choice to treat the infection, does not prevent reinfection, and its decades-long history as the main treatment raises concerns for drug resistance. Of the schistosome species, *Schistosoma haematobium* causes urogenital disease and has a strong association with bladder cancer. The possibility for drug resistance and the gap in knowledge with respect to the mechanisms driving *S. haematobium*-related bladder cancer highlight the need to better understand the biology of the infection to aid in the development of new therapeutic strategies. In this study, we used a mouse model of *S. haematobium* infection that delivers parasite eggs directly to the host mouse bladder wall, and we examined the changes in the gene expression profile of the host and the parasite by RNA-sequencing. The results corroborated previous findings with respect to the host’s inflammatory responses against the parasite eggs, as well as revealed alterations in other immune response genes that deepen our understanding of the mechanisms involved in urogenital schistosomiasis pathogenesis.

## Introduction

Schistosomes, the causative agents of schistosomiasis, affect more than 230 million people worldwide [1], with approximately 779 million people at risk of infection [2] and an estimated disease burden of 3.3 million disability-adjusted life years [3,4]. The life cycle of these parasites includes six distinct stages, featuring asexual and sexual reproduction in the intermediate snail host and definitive mammalian host, respectively [5]. In humans, infection with the freshwater larval stage of *Schistosoma* parasites results in the development of male and female worms that mate in the portal vein and migrate to the mesenteric veins of the intestines (*S. mansoni* and *S. japonicum*) or the venous plexus of the pelvis (*S. haematobium*) [1,6–8]. About 5-7 weeks (10- 12 weeks for *S. haematobium*) after exposure to the larvae, the adult worms begin to release eggs, causing a shift toward a type 2 host immune phenotype [9]. The eggs become encapsulated by immune cells, forming granulomas, and either continue the life cycle by traversing endothelial and epithelial barriers to escape via the excretory system, or remain trapped in the tissue, thereby serving as foci for chronic inflammation and eventual fibrosis [9].

Treatment with praziquantel (PZQ) can eliminate adult worms but has limited effect on immature worms [10,11], and it does not prevent reinfection [12]. Decades-long use of PZQ [13], together with the demonstration of inducible resistance to PZQ in the laboratory [14,15], highlight the need for a better understanding of parasite biology and host-parasite interactions to develop alternative therapeutics to combat schistosomiasis. While *S. mansoni* and *S. japonicum* infections cause hepatointestinal disease, *S. haematobium* infections lead to urogenital disease associated with hematuria, dysuria, fibrosis, and obstruction-induced pathology of the urinary tract [16–18]. Studies have also found a positive correlation between *S. haematobium* infection and bladder cancer, particularly in the form of squamous cell carcinoma [19], rather than the transitional cell carcinoma that is more commonly found in non-endemic regions [17], and the International Agency for Research on Cancer of the World Health Organization classifies *S. haematobium* as a Group 1 (i.e. definitive) carcinogen [20].

Recent studies to better understand and diagnose *S. haematobium* infection have included peripheral blood gene expression profiling by RNA-sequencing (RNA-seq) [21] and proteomic analysis of urine [22,23] that compared infected and non-infected patients, revealing the perturbation of cell differentiation and cell cycle pathways and potential urine protein biomarkers. Longitudinal studies comparing pre-infection, post-infection, and post-treatment patient samples could enhance the discovery of infection biomarkers but are difficult to coordinate, given uncertainties with respect to timing of infection and re-infection. The study of experimental schistosome infection offers greater control of sampling, but such studies largely rely on rodent animal models of infection, which, in the case of urogenital schistosomiasis, face the problem that natural *S. haematobium* infection in rodents leads to hepatointestinal egg deposition and pathology but no bladder or genital pathology [24]. Currently, the only rodent models that recapitulate key features of the urogenital disease observed in human infections involve directly injecting parasite eggs into the bladder wall [25,26] or vaginal wall [27] of mice. Using the bladder wall injection model, studies have explored host responses to the parasite involving: changes in the gene expression profile of the bladder [28]; natural killer T cells and their altered function [29]; macrophages [30]; changes in genome-wide methylation of the bladder [31]; urothelium-specific p53 [32,33]; concurrent natural *S. haematobium* infection [34]; interleukin-4 signaling [35]; angiogenesis, vascular permeability, and cell proliferation [36]; carcinogenesis during concurrent administration of sub-carcinogenic doses of nitrosamine compounds [37]; and changes in the protein expression profile of the bladder [38].

In this study, we sought to further characterize the biology of *S. haematobium* infection, including both host and parasite responses, by RNA-seq analysis of bladder tissue derived from the mouse bladder wall injection model. Our findings expand upon previous microarray and proteomic studies of the bladder wall injection model, revealing a broader set of differentially expressed genes in the mouse host. Consistent with previous findings, egg-injected samples showed an upregulation of host genes related to inflammation, immune cell chemotaxis, and cytokine signaling. We also observed the upregulation of key immune checkpoint-related genes, which may represent a mechanism directed at dampening the inflammatory response in this chronic disease. In addition to profiling changes in gene expression of the mouse host, we attempted to profile changes in gene expression of parasite genes in the bladder wall injection model. We found that the parasite genes showed an overall downregulation and that the bladder wall-injected samples provided drastically fewer sequencing reads compared to the number of reads obtained in the analysis of mouse genes. These findings improve our understanding of *S. haematobium*-related bladder pathology and disease modelling, providing clues for future investigations to identify mechanisms driving the disease.

## Methods

### Ethics statement

Animal work was conducted according to relevant U.S. and international animal use guidelines. Animal experimental protocols were reviewed and approved by the Institutional Animal Care and Use Committee (IACUC) of the Biomedical Research Institute (Rockville, MD, United States) and Children’s National Research Institute (Washington, DC, United States, protocol #00030764). These guidelines comply with the U.S. Public Health Service Policy on Humane Care and Use of Laboratory Animals.

### Mice and hamsters

Female 8-12 week old BALB/c mice and 8-12 week old Golden Syrian hamsters were obtained from Envigo and Charles River Laboratories, respectively, maintained under 12-hour light/dark cycle, and were provided unrestricted access to food and water.

### Parasite eggs

The *S. haematobium* parasite eggs were graciously prepared by the Schistosome Resource Center at Biomedical Research Institute [39,40]. In summary, Golden Syrian hamsters infected with *S. haematobium* were perfused and their egg-containing livers and intestines were placed in 1.2% saline at 4 °C overnight, followed by mechanical disruption of the tissues with a blender and flushing with cold 1.2% saline through a succession of sieves of decreasing mesh size. The resulting eggs were swirled for further separation from remaining tissues and washed with cold PBS containing 100 U/mL penicillin-streptomycin solution (Gibco #15140122) prior to bladder wall injection. Hamster liver and intestine homogenate was prepared from uninfected hamsters in a similar manner as above and used for the vehicle control samples.

### Bladder wall injection

Mouse bladder wall injection was performed as previously described [26,41]. Mice were anesthetized and maintained under 1.5-2% isoflurane and placed on a warming pad, followed by depilation of the abdomen with depilatory cream, disinfection of the surgical site with iodine solution, and subcutaneous injection with bupivacaine (8 mg/kg) and carprofen (5 mg/kg). A midline incision was made to expose the bladder. A 100 µL capacity 1700-series glass syringe (Hamilton #7656-01) fitted with a 29-gauge, 10 mm length, point style 4, 12° bevel needle (Hamilton #7803-06) was used to deliver two injections of a 50 µL volume of PBS (with 100 U/mL penicillin-streptomycin) containing either uninfected hamster liver and intestine homogenate (vehicle control) or 3000 *S. haematobium* eggs to the bladder wall. The syringe and needle were flushed with PBS between each injection. Following bladder wall injection, the peritoneal and skin incisions were respectively closed by continuous and interrupted suturing, using 4-0 size PGA and silk suture fitted with a 19 mm 3/8 circle reverse cutting edge NFS-2 needle (Oasis #MV-J397 and #MV-683). Antibiotic ointment was applied to the surgical area, and mice were withdrawn from isoflurane. Bladder wall injection for hamsters was performed in a similar manner as described above, with five injections instead of two injections.

### RNA isolation

Mice and hamsters were euthanized by CO_2_ asphyxiation followed by cervical dislocation, and their bladders were excised and placed into 2 mL conical-bottom screw-cap tubes (USA Scientific #1420-8700) on ice containing 1 mL TRIzol (Invitrogen #15596018) and six 2.8 mm diameter ceramic beads (Omni International #19-646). For “egg-alone” samples, *S. haematobium* eggs were used in the absence of bladder tissues. For “egg/bladder mix” samples, eggs mixed with bladder tissue from mice that did not undergo bladder wall injection were used.

The samples were homogenized by bead beating (BeadBug 3 Microtube Homogenizer, Benchmark Scientific #D1030) for 3 sets of 30 seconds alternating with 30 seconds of incubation on ice. After homogenization, the samples were frozen at -80 °C until subsequent processing.

RNA was extracted from the samples following the manufacturer’s instructions for quick-spin column purification of RNA samples suspended in TRIzol (PureLink RNA Mini Kit, Invitrogen #12183018A) with on-column DNase treatment (PureLink DNase, Invitrogen #12185010). The resulting RNA was quantitated by spectrophotometry (NanoDrop 8000, Thermo Scientific) and stored at -80 °C.

### RNA sequencing

RNA samples were sent to BGI for capillary electrophoresis (Bioanalyzer, Agilent Technologies)-based RNA quality analysis, followed by library preparation and sequencing on the DNBseq platform for 20 million paired-end 100 bp reads. The hamster bladder wall injection samples were subjected to a sequencing depth of 200 million paired-end 100 bp reads.

### Bioinformatics

Files containing RNA-sequencing reads were obtained from BGI and processed through bioinformatics software, including FastQC, STAR, featureCounts, and DESeq2. Genes related to immunoglobulin variable regions (Supplementary 1) were removed from the list of differentially expressed genes. Further analysis to associate differentially expressed genes to biological phenotypes and signaling pathways was conducted with gene set enrichment analysis (GSEA, Broad Institute), PANTHER, and Ingenuity Pathway Analysis (IPA, Qiagen). Unless otherwise specified, the following cutoffs were applied: absolute log_2_ fold change (|log_2_FC|) > 1 and adjusted *p*-value (*p*-adj) < 0.05. Scripts (Supplementary 2) and detailed methods (Supplementary 3) for the analysis are available.

### Estimation of parasite gene transcript expression level

To estimate the level of parasite gene transcript expression following bladder wall injection, RNA was isolated as described above, followed by cDNA synthesis using the iScript cDNA synthesis kit (Bio-Rad #1708891), and 20 ng of each cDNA sample, together with 500 nM forward and reverse primers, were used in triplicate 20 µL reactions of quantitative PCR using the iTaq Universal SYBR Green Supermix (Bio-Rad #1725121) in a CFX Connect Real-Time PCR Detection System (Bio-Rad #1855201) with the following thermal cycling conditions: (95 °C for 30 sec) × 1 cycle, (95 °C for 5 sec; 60 °C for 30 sec) × 40 cycles. The primers were chosen using the Primer3 software [42] and obtained from Eurofins Genomics. The primers targeting *S. haematobium* IPSE were derived from GenBank accession MG012894.1 (MS3_0018942), and the primers targeting *S. haematobium* COX1 were derived from GenBank accession DQ677664.1 (Supplementary 4).

## Results

### Mouse bladder wall injection with S. haematobium eggs induces changes in gene expression in the bladder wall

To better understand the effects of *S. haematobium* eggs on the host bladder, we performed bladder wall injection on mice, and followed this with RNA-seq of the bladder, 4 days post-injection. In this report, we present data pooled from two independent sets of bladder wall injection experiments, which included mice that underwent bladder wall injection with vehicle control (“hinj”) and mice that underwent bladder wall injection with eggs (“einj”), with initially 5 mice per group (Table 1). Other sequenced samples included mice that did not undergo surgery (“naive”), egg-alone (“egg”), and egg mixed with bladder tissue from mice that did not undergo bladder wall injection (“eb”).

**Table 1.**
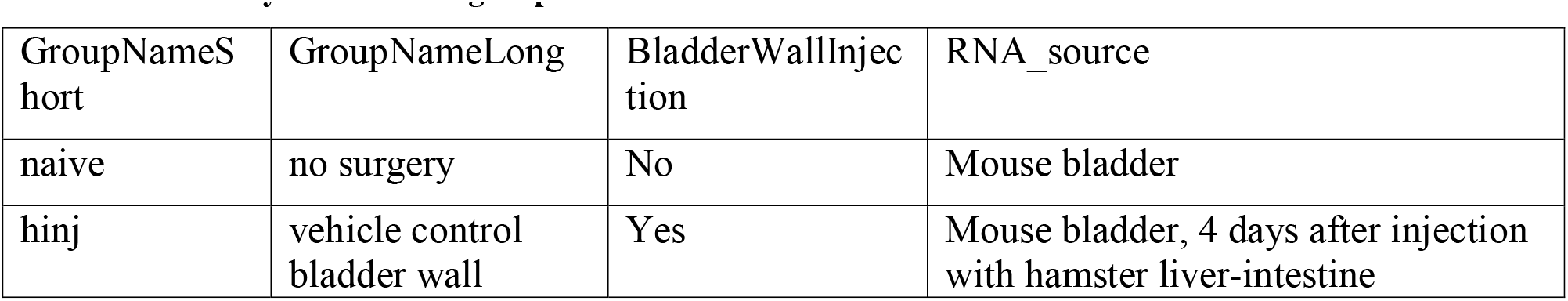
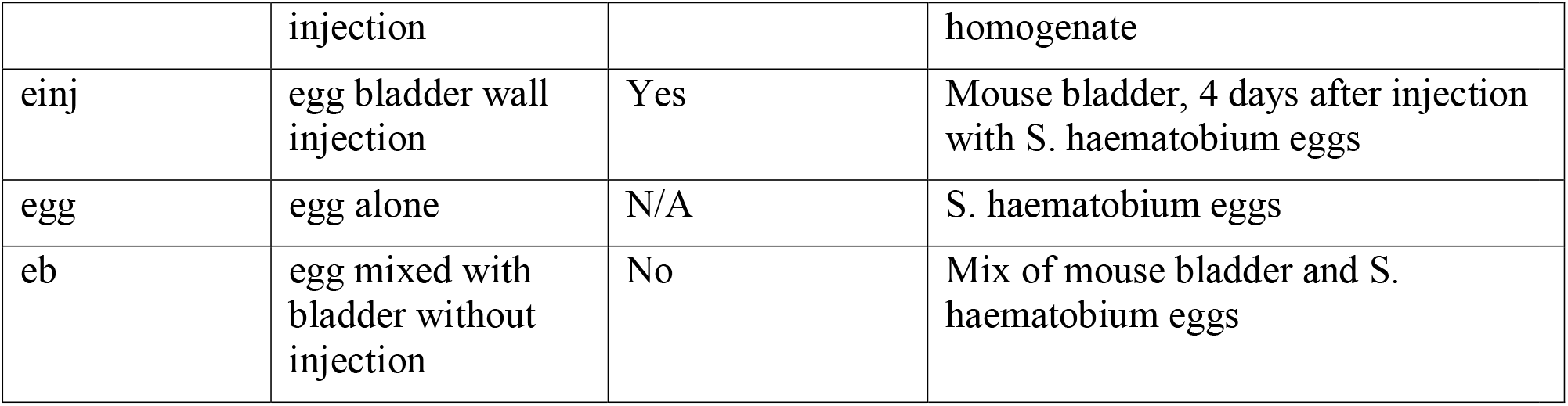
Summary of treatment groups.

| GroupNameShort | GroupNameLong | BladderWallInjection | RNA_source |
| --- | --- | --- | --- |
| naive | no surgery | No | Mouse bladder |
| hinj | vehicle control bladder wall | Yes | Mouse bladder, 4 days after injection with hamster liver-intestine |
|  | injection |  | homogenate |
| einj | egg bladder wall injection | Yes | Mouse bladder, 4 days after injection with *S. haematobium* eggs |
| egg | egg alone | N/A | *S. haematobium* eggs |
| eb | egg mixed with bladder without injection | No | Mix of mouse bladder and *S. haematobium* eggs |

Principal component analysis (PCA) of the sequencing reads aligned to the mouse genome qualitatively showed distinct clustering of samples among treatment groups (Figure 1A). In the “hinj” and “einj” groups, 21957 genes showed a non-zero numerical value for both Benjamini & Hochberg adjusted *p* (*p*-adj) and log_2_ fold change (log_2_FC), which, after applying a *p*-adj value cutoff (*p*-adj < 0.05) and log_2_FC absolute value cutoff (|log_2_FC| > 1), yielded 1025 genes (Figure 1B, Table 2; Supplementary 5). During the bladder wall injection procedure, several mice in the egg-injection group did not receive eggs because of technical error, and PCA of the sequencing reads from these samples showed that they cluster closely with the vehicle-injected samples (Supplementary 6). Subsequent analyses in this report omit these samples; however, the sequencing data are available for all experiments (see Data availability).

**Figure 1.**
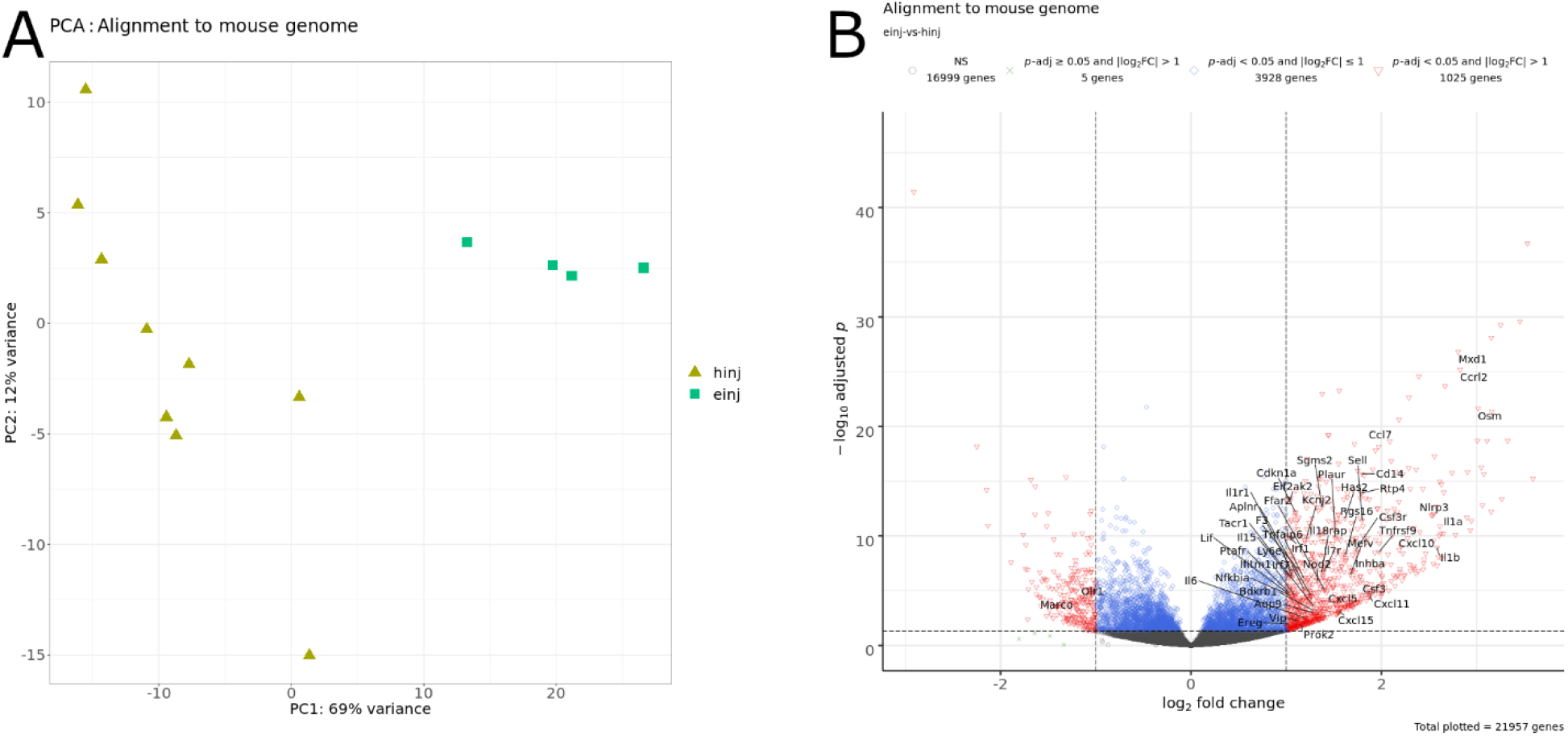
Comparison of samples aligned to the mouse genome. A) Principal component analysis. Each point represents a sample, color-coded by treatment group. Key: hinj, vehicle-injected (olive-yellow filled triangles); einj, *S. haematobium* egg-injected (cyan-green filled squares). B) Volcano plot labeled with significantly differentially expressed genes (*p*-adj < 0.05 and |log2FC| > 1) associated with the inflammatory response gene set from the molecular signatures database (MSigDB) in the comparison of egg-injected and vehicle-injected samples aligned to the mouse genome. Key: NS, not significant.

**Table 2.**
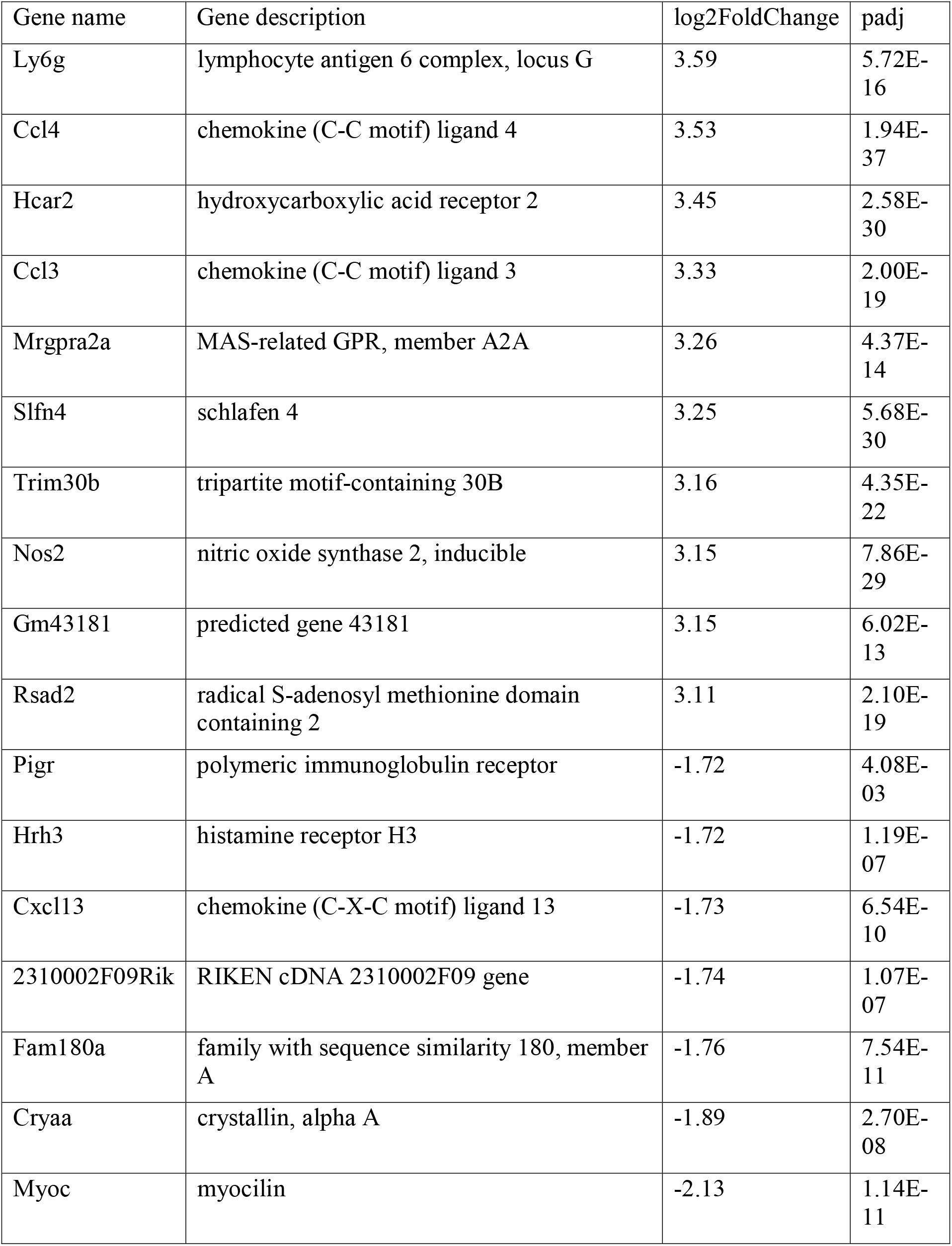
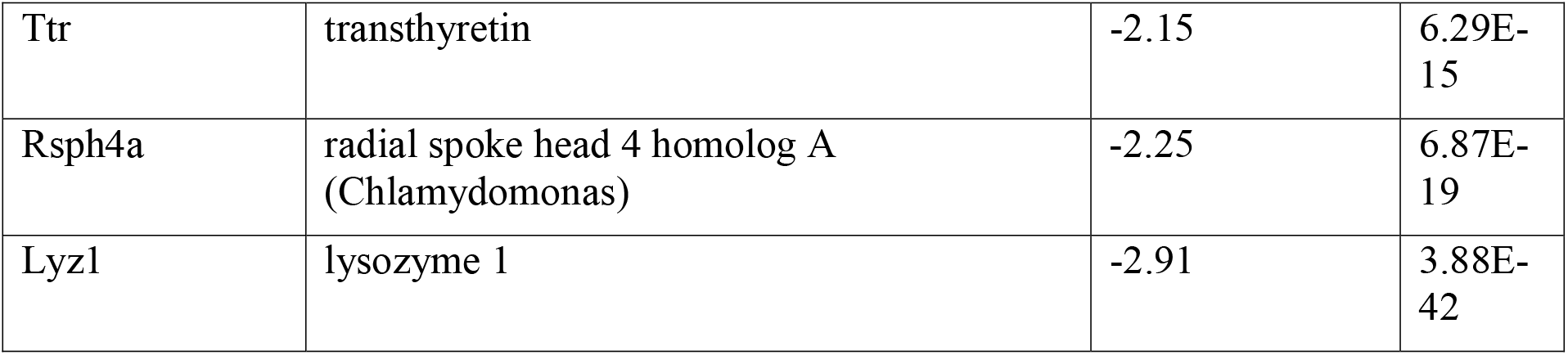
Top 10 most up- and down-regulated mouse genes (*p*-adj < 0.05; |log_2_FC| > 1) in the comparison of egg-injected and vehicle-injected samples.

Gene set enrichment analysis (GSEA) revealed 18 positively enriched and 4 negatively enriched mouse hallmark gene sets (“MH”, FDR q-val < 0.05) in the comparison of egg-injected samples and vehicle control-injected samples (Supplementary 7). Analysis of the differentially expressed genes by the PANTHER gene ontology overrepresentation test revealed that upregulated genes belonged to biological processes related to chemotaxis of immune cells, cytokine signaling, and inflammatory response (Figure 2A, Supplementary 8). The overrepresentation test for downregulated genes did not show an enrichment for any specific gene ontology or pathway.

**Figure 2.**
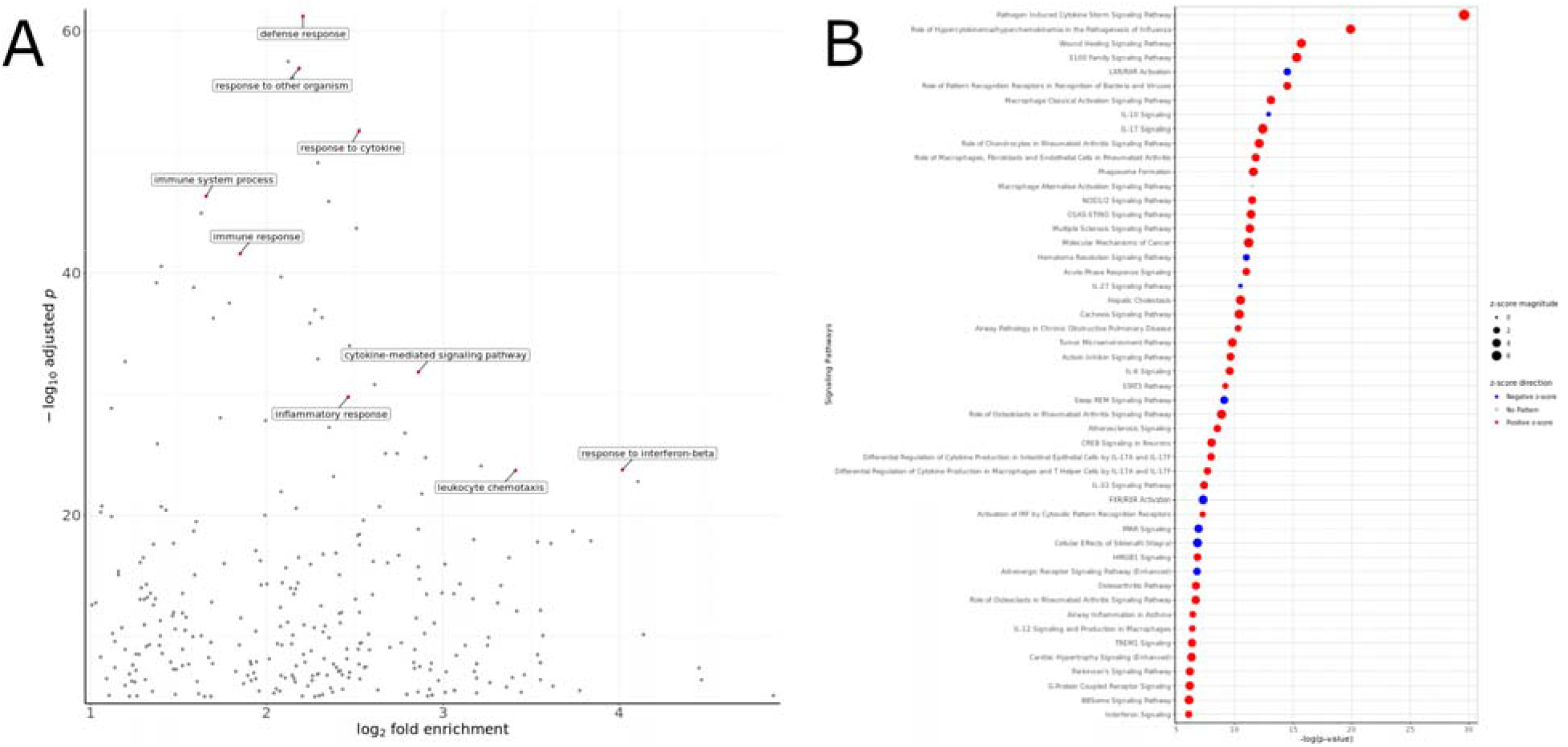
Gene ontology and pathways analysis of differentially expressed genes in the comparison of egg-injected and vehicle-injected samples aligned to the mouse genome. A) PANTHER gene ontology analysis of upregulated genes, with selected biological process gene ontologies labeled. B) IPA analysis for signaling pathways. Positive z-scores (red color) represent positive enrichment, while negative z-scores (blue color) represent negative enrichment.

Ingenuity Pathway Analysis (IPA) for signaling pathways showed a positive enrichment for pathways related to cytokine storm, wound healing, and macrophage activation; and negative enrichment for pathways related to liver X receptor (LXR), interleukin (IL)-10, and hematoma resolution (Figure 2B, Supplementary 9).

### S. haematobium eggs upregulate inflammatory response genes in the bladder wall

The most strongly upregulated genes (*p*-adj < 0.05 and log_2_FC > 1) included inflammatory response-related genes, such as the chemokines *Ccl3* and *Ccl4* (encodes MIP-1a and MIP-1b, respectively) (Table 2). Other upregulated genes related to inflammatory response included the chemokines *Cxcl2* (encodes MIP-2a), *Cxcl3* (encodes MIP-2b), *Ccl11* (encodes eotaxin-1), *Cxcl1* (encodes KC), and *Ccl7* (encodes MCP-3); and the cytokines *Il1a*, *Il1b*, *Il6*, *Tnf*, and related receptors (Supplementary 5). GSEA analysis showed positive enrichment of gene sets related to inflammatory response and IL-6/JAK-STAT3 signaling (Figure 3).

**Figure 3.**
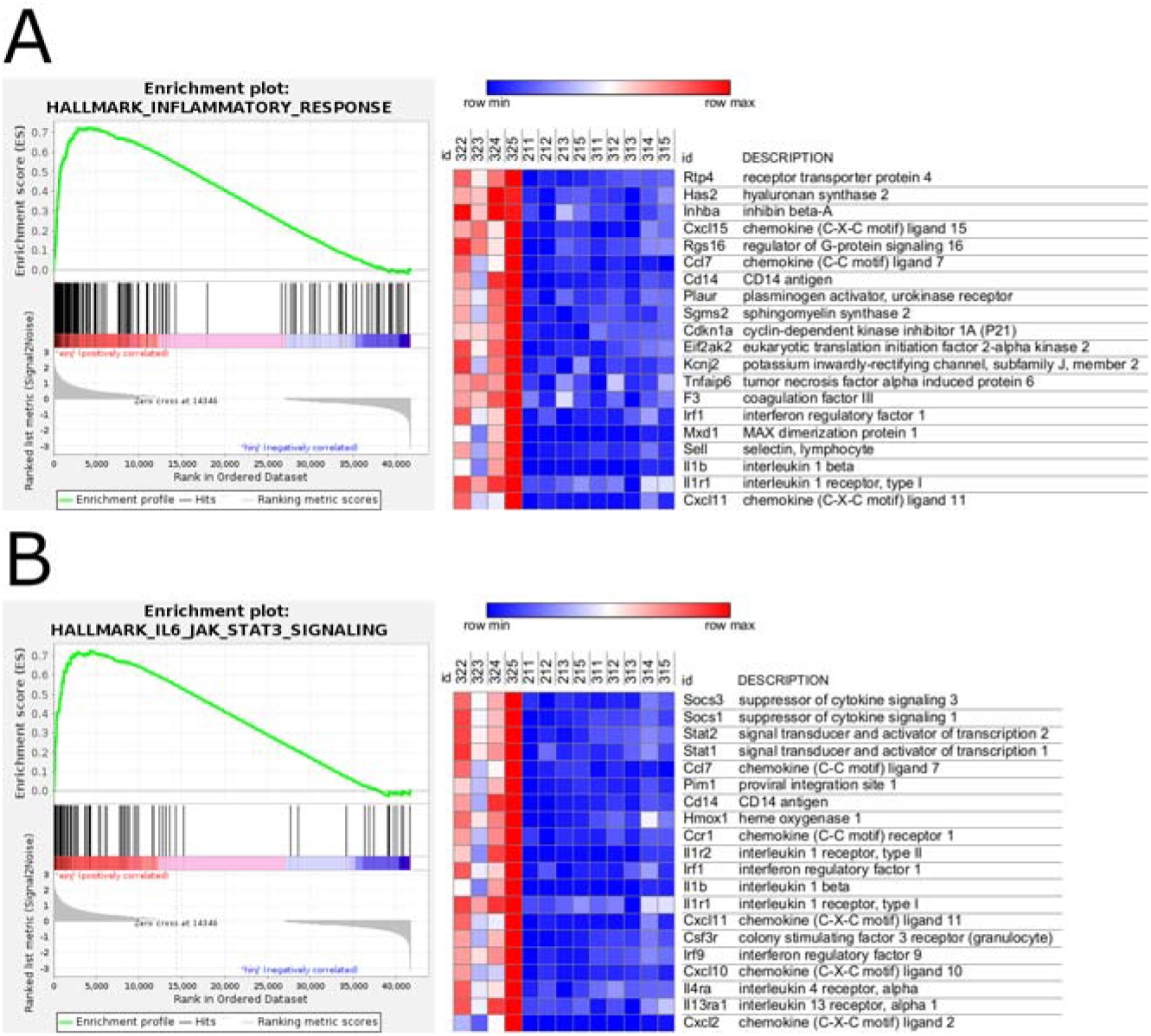
GSEA enrichment plot (left) and relative gene expression (right) of those belonging to the mouse hallmark gene sets related to A) inflammatory response and B) IL6/JAK-STAT3 signaling, in the comparison of egg-injected and vehicle-injected samples aligned to the mouse genome. Top 20 genes associated with each gene set, arranged by descending rank metric score, shown across 4 egg-injected (322-325) and 9 vehicle-injected (211-213, 215, 311-315) samples, with relative gene expression level represented in a blue (lowest expression)-red (highest expression) color gradient.

### S. haematobium eggs upregulate genes associated with immune phenotype and immune checkpoint in the bladder wall

The RNA-seq analysis showed the upregulation of the type 2 immunity cytokines *Il4* (log_2_FC = 0.42) and *Il13*; the immune regulatory gene *Foxp3*; the stimulatory immune checkpoint genes *Tnfrsf4* (encodes OX40), *Tnfrsf9* (encodes 4-1BB), and *Icos*; and the inhibitory immune checkpoint genes *Ctla4*, *Havcr2* (encodes TIM-3), *Cd274* (encodes PD-L1), and *Pdcd1lg2* (encodes PD-L2) (Supplementary 5, Supplementary 10).

### S. haematobium eggs alter the expression of genes involved in bladder hemostasis, integrity, and homeostasis

The RNA-seq analysis showed the downregulation (log_2_FC < -0.585) of the uroplakins *Upk1a*, *Upk1b*, *Upk2*, *Upk3a*, and *Upk3b*; and the claudins *Cldn8* and *Cldn10*, which play a role in urothelial barrier function (Supplementary 10). The analysis also showed the upregulation (log2FC > 0.585) of the angiogenesis-related genes *Vegfa* and *Vegfd* and matrix metalloproteinases (MMPs), such as *Mmp3*, *Mmp8*, and *Mmp9*, which play a role in remodeling of the extracellular matrix (Supplementary 10).

In the GSEA analysis, the hallmark gene sets related to KRAS signaling upregulation, epithelial-mesenchymal transition, hypoxia, and angiogenesis, which can play a role in cancer, appeared among the positively enriched gene sets (Supplementary 7). Consistent with hypoxia, gene sets related to oxidative phosphorylation and fatty acid metabolism appeared among the negatively enriched gene sets (Supplementary 7). In contrast, gene sets related to G2-M cell cycle checkpoint, apoptosis, and p53 signaling, which can play a role in impeding cancer progression, also appeared among the positively enriched gene sets (Supplementary 7).

### Comparison of the RNA-seq analysis with previous studies highlights key mouse genes involved in the bladder wall injection model of S. haematobium infection

Our previous studies of the mouse bladder wall egg injection model of *S. haematobium* infection included microarray and proteomic analyses. We compared the differentially expressed genes and proteins found in the previous studies with the differentially expressed genes found in the current RNA-seq study, reasoning that similarities among the results can help identify key host genes and pathways involved during parasite egg deposition. The comparison of the differentially expressed genes found in the RNA-seq analysis in this study (4 days post-egg injection, *p*-adj < 0.05, |log_2_FC| > 1) and data from the previous mouse bladder gene expression microarray analysis (week 1 post-egg injection) [28] revealed 70 genes common to both datasets sharing the same direction of differential expression that have associations with gene ontology biological process terms such as response to cytokine, neutrophil chemotaxis, and inflammatory response (Figure 4, Supplementary 10, Supplementary 11). The RNA-seq and proteomic (week 1 post-egg injection) [38] analyses showed the upregulation of *Steap4* (six-membrane epithelial antigen of the prostate, member 4); the microarray and proteomic analyses showed the upregulation of *Stab1* (stabilin 1); and all three studies showed the upregulation of *Serpina3n* (serine protease inhibitor, clade A, member 3N) (Figure 4, Supplementary 10).

**Figure 4.**
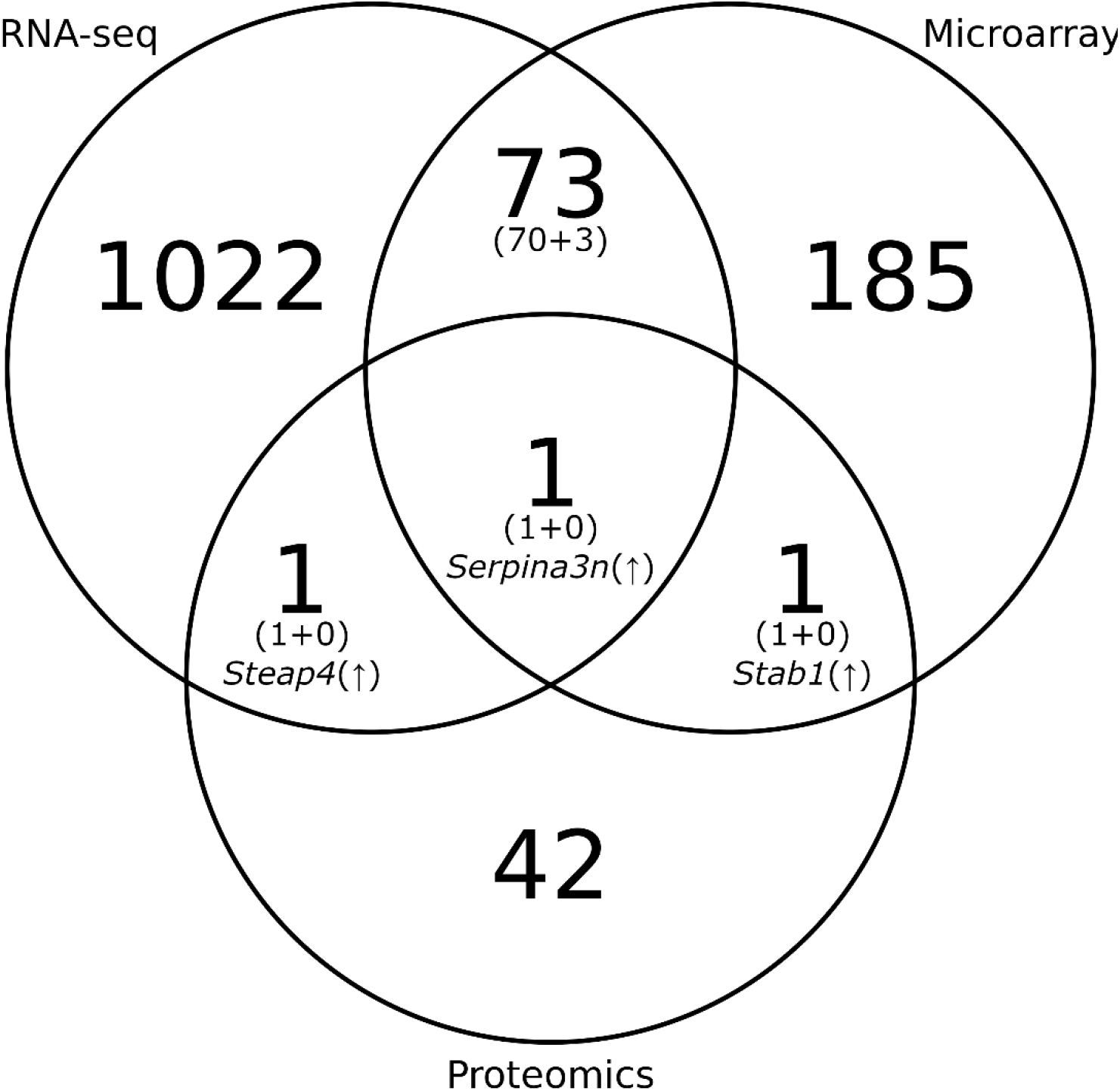
Venn diagram of differentially expressed genes in the egg-injected group and vehicle-injected group in the *S. haematobium* mouse bladder wall injection model by RNA-seq (*p*-adj < 0.05, |log_2_FC| > 1, day 4), microarray (week 1), and proteomic (week 1) analyses. Numbers in parentheses indicate the number of genes that share the same direction of differential expression (concordance) followed by the number of those that do not (discordance).

### Mouse bladder wall injection with S. haematobium eggs induces changes in parasite gene expression

To determine the effect of the bladder environment on the parasite eggs and their gene expression, we aligned the RNA-seq reads to the *S. haematobium* genome, focusing on comparisons among egg-injected (“einj”), egg mixed with bladder tissue from mice that did not undergo bladder wall injection (“eb”), and egg-alone samples (“egg”). Alignment to the *S. haematobium* genome resulted in the highest alignment rate (as a proportion of total number of sequencing reads), as well as the highest raw number of aligned sequencing reads in the “egg” samples (sample number 234), followed by a marked decrease in these numbers in the “eb” (sample numbers 241-245) and “einj” samples (sample numbers 322-325) (Supplementary 12). PCA showed clustering of the “eb” and egg-alone “egg” samples, as well as an egg-alone sample obtained from *S. haematobium* genome reference data (R31) [43–46], distinct from the egg-injected (“einj”), no-surgery (“naive”), and vehicle-injected (“hinj”) samples (Supplementary 13A). However, the PCA for pooled “eb” and “egg” samples compared against “einj” samples showed a separation of the R31 reference sample from the pooled “eb” and ”egg” samples (Supplementary 13B), prompting us to remove the reference sample from the differential gene expression analysis (Figure 5A). Differential gene expression analysis in the comparison of egg-injected “einj” samples against the egg-alone-clustered (pooled “eb” and “egg”) samples showed 3306 genes with a non-zero numerical value for both *p*-adj and log_2_FC, which, after applying a *p*-adj value cutoff (*p*-adj < 0.05) and log_2_FC absolute value cutoff (|log_2_FC| > 1), yielded 119 genes (Figure 5B). Upregulated parasite genes included those likely related to transcription, such as KRAB-A domain-containing protein; intracellular signaling via kinase activity; metabolism, such as glycine dehydrogenase and diacylglycerol O-acyltransferase 1; and protein (re)folding, such as heat shock protein 70 (Table 3, Supplementary 14). Downregulated parasite genes included those likely related to ribosomal activity and cell death (Table 3, Supplementary 14).

**Figure 5.**
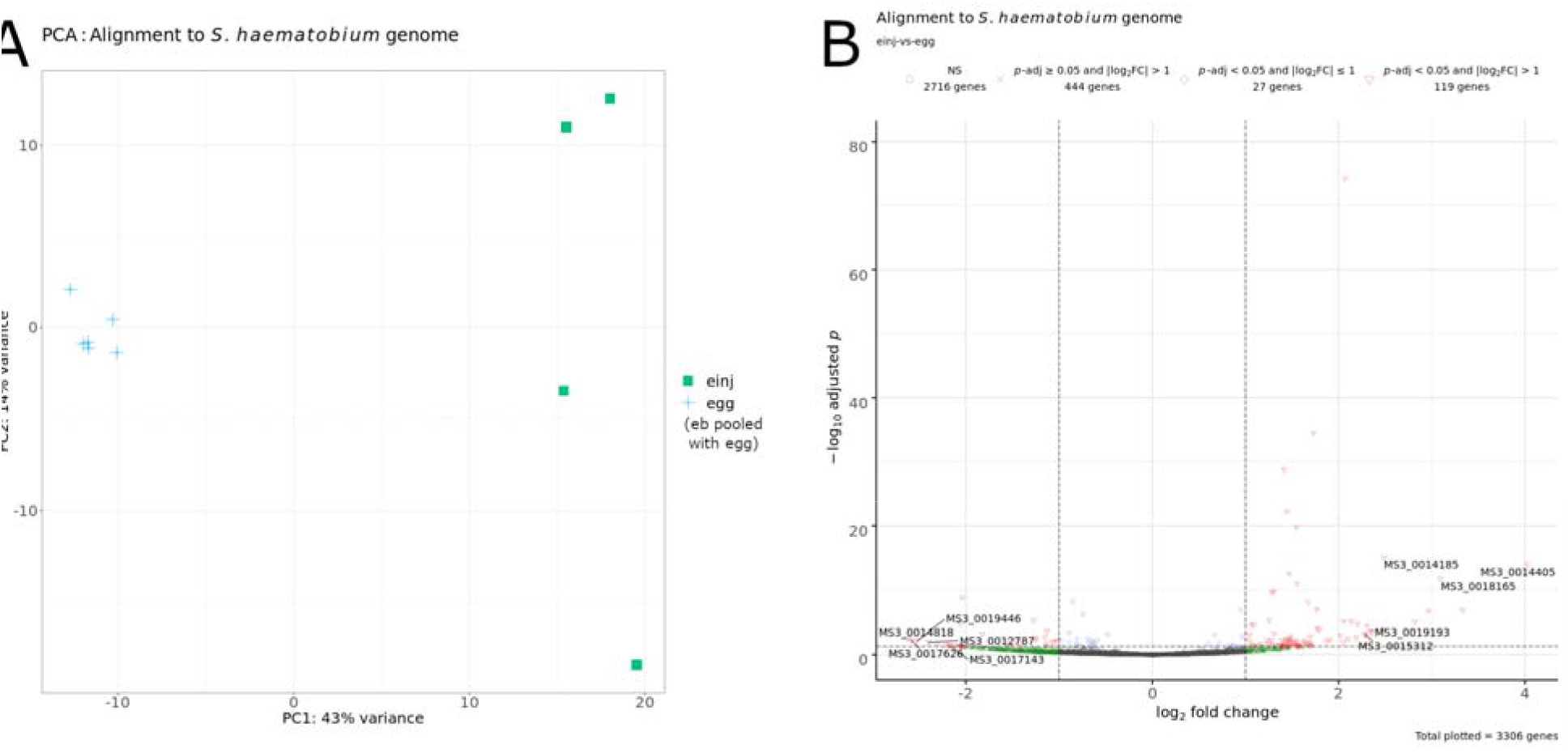
Comparison of samples aligned to the *S. haematobium* genome. A) Principal component analysis. Each point represents a sample, color-coded by treatment group. Key: einj, *S. haematobium* egg-injected (cyan-green filled squares); egg, *S. haematobium* egg alone, pooled with “eb” samples (*S. haematobium* egg mixed with bladder tissue from mice that did not undergo bladder wall injection) (blue crosses). B) Volcano plot labeled with a subset of differentially expressed genes in the comparison of egg-injected (“einj”) and pooled egg (“eb” pooled with “egg”) samples. Key: NS, not significant.

**Table 3.**
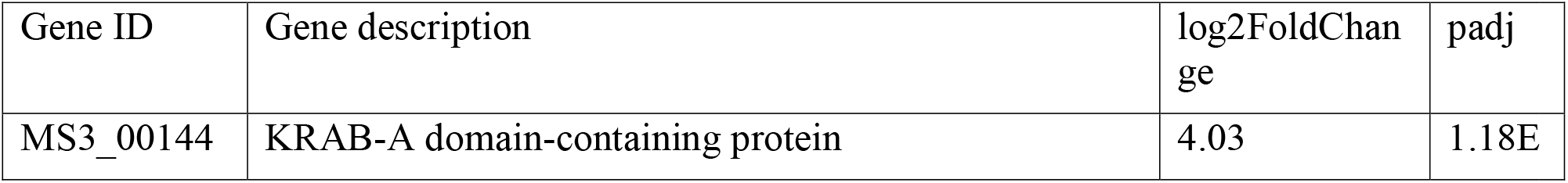
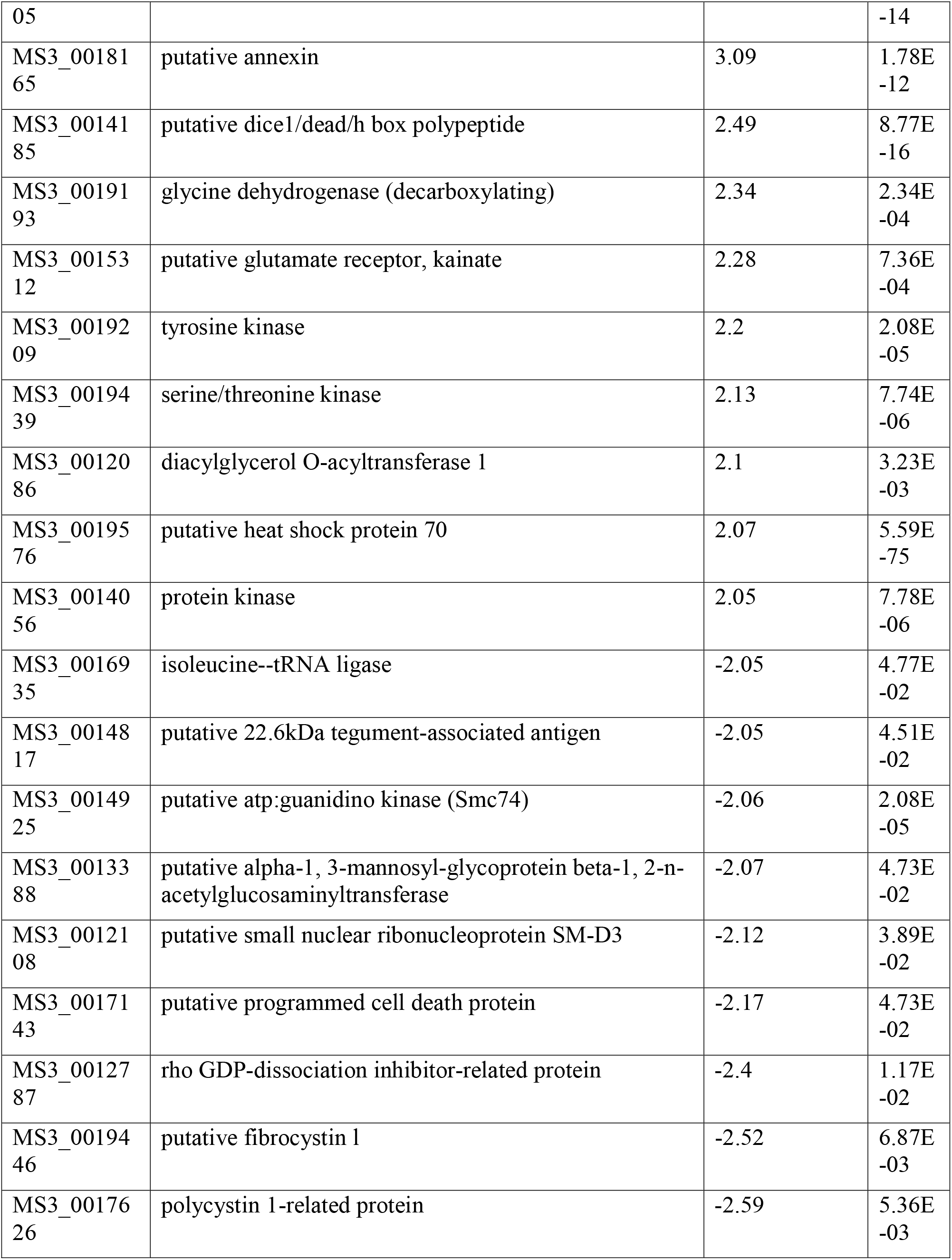
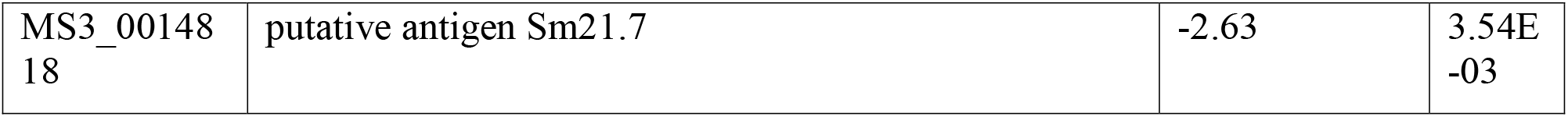
Top strongly up- and down-regulated *S. haematobium* genes (*p*-adj < 0.05; |log_2_FC| > 1) of known or putative function in the comparison of egg-injected (“einj”) and pooled egg (“eggs mixed with bladder tissue from mice that did not undergo bladder wall injection” (“eb”) pooled with “egg-alone” (“egg”)) samples.

| Gene ID | Gene description | log2FoldChange | padj |
| --- | --- | --- | --- |
| MS3_00144 | KRAB-A domain-containing protein | 4.03 | 1.18E |
| 05 |  |  | -14 |
| MS3_00181<br>65 | putative annexin | 3.09 | 1.78E<br>-12 |
| MS3_00141<br>85 | putative dice1/dead/h box polypeptide | 2.49 | 8.77E<br>-16 |
| MS3_00191<br>93 | glycine dehydrogenase (decarboxylating) | 2.34 | 2.34E<br>-04 |
| MS3_00153<br>12 | putative glutamate receptor, kainate | 2.28 | 7.36E<br>-04 |
| MS3_00192<br>09 | tyrosine kinase | 2.2 | 2.08E<br>-05 |
| MS3_00194<br>39 | serine/threonine kinase | 2.13 | 7.74E<br>-06 |
| MS3_00120<br>86 | diacylglycerol O-acyltransferase 1 | 2.1 | 3.23E<br>-03 |
| MS3_00195<br>76 | putative heat shock protein 70 | 2.07 | 5.59E<br>-75 |
| MS3_00140<br>56 | protein kinase | 2.05 | 7.78E<br>-06 |
| MS3_00169<br>35 | isoleucine--tRNA ligase | -2.05 | 4.77E<br>-02 |
| MS3_00148<br>17 | putative 22.6kDa tegument-associated antigen | -2.05 | 4.51E<br>-02 |
| MS3_00149<br>25 | putative atp:guanidino kinase (Smc74) | -2.06 | 2.08E<br>-05 |
| MS3_00133<br>88 | putative alpha-1, 3-mannosyl-glycoprotein beta-1, 2-n-acetylglucosaminyltransferase | -2.07 | 4.73E<br>-02 |
| MS3_00121<br>08 | putative small nuclear ribonucleoprotein SM-D3 | -2.12 | 3.89E<br>-02 |
| MS3_00171<br>43 | putative programmed cell death protein | -2.17 | 4.73E<br>-02 |
| MS3_00127<br>87 | rho GDP-dissociation inhibitor-related protein | -2.4 | 1.17E<br>-02 |
| MS3_00194<br>46 | putative fibrocystin 1 | -2.52 | 6.87E<br>-03 |
| MS3_00176<br>26 | polycystin 1-related protein | -2.59 | 5.36E<br>-03 |
| MS3_00148<br>18 | putative antigen Sm21.7 | -2.63 | 3.54E<br>-03 |

### Parasite gene expression in S. haematobium eggs declines rapidly following bladder wall injection

Following the observation that bladder wall-injected egg samples produced fewer sequencing reads when aligned to the *S. haematobium* genome in comparison to other egg-alone (“egg”) and egg/bladder mix (“eb”) samples, we sought to determine the stability of parasite gene transcript expression levels after bladder wall egg injection. We conducted a small experiment (1 mouse per time point) over the course of 4 days following injection, which revealed a considerable loss of signal in all bladder wall-injected samples when compared to the pre-injection (day 0) control sample in the gene expression level of *Ipse/alpha-1* (interleukin 4 inducing principle from *Schistosoma mansoni* eggs, a highly expressed gene in schistosome eggs) and *Cox1* (cytochrome oxidase subunit 1, a housekeeping gene) *S. haematobium* gene transcripts (Supplementary 15).

### Egg bladder wall injection in hamsters compared to bladder wall injection in mice results in a greater number of reads aligning to the S. haematobium genome

To determine whether we could obtain greater alignment rates to the *S. haematobium* genome by increasing the number of injected eggs, we conducted another small experiment (n=1 per group) in which we performed bladder wall injection in hamsters with eggs or vehicle control, followed by RNA-seq. In the mouse bladder wall injection experiment, the mice received two injections of 3000 eggs each (6000 eggs total), while the hamster received five injections (15000 eggs total). We also increased the RNA-seq read depth from 4e9 bp (∼20 million paired-end 100 bp reads) in the mouse experiment to 2.4e10 bp (∼120 million paired-end 100 bp reads) in the hamster experiment. For the egg-injected hamster sample, out of a total of around 130 million reads, the alignment to the hamster genome yielded around 120 million (92%) uniquely mapped reads, while the alignment to the *S. haematobium* genome yielded around 30000 (0.03%) uniquely mapped reads (Supplementary 12). In comparison, the egg-injected mouse samples (sample numbers 322-325), out of a total of around 25 million reads, the alignment to the mouse genome yielded around 23 million (90%) uniquely mapped reads, while the alignment to the *S. haematobium* genome yielded around 12000-20000 (0.03% to 0.07%) uniquely mapped reads (Supplementary 12).

## Discussion

Schistosomiasis is a major source of morbidity and mortality and a public health challenge. This highlights the need to better understand the parasite biology in the context of human-host interactions to help develop improved diagnostics, vaccines, and new therapeutics. This need especially pertains to urogenital schistosomiasis, whose study lags behind those of other schistosome species because of the difficulty in replicating the human urogenital disease in tractable animal models. In this study, we characterized the gene expression profile of both the mouse bladder and parasite eggs in the *S. haematobium* mouse bladder wall injection model by RNA-seq analysis. We chose to analyze egg-injected bladder samples 4 days after injection, in contrast to the longer periods (1, 3, and 5 weeks) of a previous microarray-based gene expression analysis [28], to measure parasite gene expression to minimize the effect of degradation of parasite RNA. We also chose to profile the gene expression by RNA-seq analysis, in contrast to microarray analysis, to take advantage of its greater sensitivity in detecting gene transcripts, and to measure gene expression differences in not only the host but also the parasite.

The host gene expression profile of the egg-injected samples compared to vehicle control-injected samples showed the upregulation of genes associated with ontologies and pathways related to inflammatory response, immune cell chemotaxis, and interleukin and cytokine signaling and the enrichment of genes associated with interferon-gamma and -alpha signaling, tumor necrosis factor-alpha signaling, inflammatory response, and the IL6-JAK/STAT3 signaling gene sets. Previous analyses of the mouse bladder wall injection model of *S. haematobium* bladder infection [26,28] similarly showed an upregulation of genes involved in inflammatory host response. For example, the RNA-seq and proteomics analyses both showed the upregulation of *Steap4* (six-transmembrane epithelial antigen of the prostate, member 4), a metalloreductase that is upregulated in response to inflammatory cytokines and thought to promote metabolic homeostasis [47,48]. Studies have shown that *Steap4*, also known as *Tiarp* (tumor necrosis factor-induced adipose related protein) and *Stamp2* (six-transmembrane protein of prostate, member 2), can increase the generation of reactive oxygen species, contributing to the development of colon and prostate cancer [49,50].

Accompanying the upregulated inflammatory host response, the RNA-seq analysis showed the enrichment of genes that belong to the apoptosis gene set. This finding contrasts with those of a study that exposed epithelial cells to *S. haematobium* worm antigen protein extract and observed decreased protein expression of the tumor suppressor p27 (CDKN1B) and an increased number of cells expressing the anti-apoptosis factor BCL2, suggesting that the infection can suppress apoptosis [51]. The RNA-seq analysis showed the upregulation of *Cdkn1b* (log_2_FC = 0.24) and downregulation of *Bcl2* (log_2_FC = -0.19; Supplementary 5, Supplementary 10). This discrepancy could result from the difference in the response of epithelial cells in isolation, compared to the bladder wall injection model, which includes the response of infiltrating cells and other resident cells in addition to bladder epithelial (urothelial) cells. Alternatively, the enrichment of genes in the apoptosis gene set may instead indicate the increased activity of another cell death program associated with inflammation, such as necroptosis or pyroptosis, since these programs share several signaling components with apoptosis [52].

Schistosome infection modulates the host immune system in a temporally distinct manner, beginning with the induction of responses characterized by type 1 helper T (Th1) cells during the early infection phase, type 2 helper T (Th2) cells coinciding with egg release, and regulatory (Treg) cells during the chronic infection phase [53,54]. While the RNA-seq analysis did not show the upregulation of the immunoregulatory cytokines *Il10* and *Tgfb1*, it showed the upregulation of *Foxp3* and *Il13* (and to a lesser degree, *Il4*, Supplementary 5, Supplementary 10), consistent with the idea that the parasite eggs dampen host inflammatory responses to prevent excessive damage to the host [55]. FOXP3 serves as a key factor in the development and maintenance of regulatory Treg cells, which play a role in immune tolerance and suppression [56]. IL-4 and IL-13 both promote type 2 immune responses that can induce immunoglobulin class switch to IgE and tissue repair through the activity of Th2 cells [57] and alternatively activated (M2) macrophages [58]. In concordance with the upregulation of these genes observed in this analysis, studies of mouse *S. mansoni* infection found: an increase in Treg and Th2 cell responses following exposure to egg antigens [59]; an increase in the frequency of Treg cells in liver granulomas during the chronic (16 weeks post-infection) [60], but not early (first 3 weeks of infection) [61], phases of infection; that retroviral overexpression of *Foxp3* during infection results in decreased granuloma size [60], likely explained by Treg cell-dependent suppression of Th2 cell responses [62]; and that IL-10 plays a limited role in the Treg response to the infection [59,60,63]. *S. mansoni* infection in mice also induces the expansion of the M2 macrophage population through a T cell-derived IL-4 and IL-13 mechanism, resulting in the protection against fatal pathology, likely by suppressing type 1 immune responses [64,65]. Consistent with an increase in the number of M2 macrophages, the RNA-seq analysis showed the upregulation of M2 macrophage-associated markers [58], such as *Il1r2* and *Clec7a* (encodes DECTIN1). The previous microarray analysis and proteomic analysis (but not the RNA-seq analysis) showed the upregulation of *Stab1* (stabilin-1, also known as MS-1 antigen). Stabilin-1 is expressed by endothelial cells and M2 macrophages [66], participates in receptor-mediated endocytosis and intracellular transport [67], has been proposed to serve as a prognostic marker for bladder cancer [68], and may confer resistance to immune checkpoint cancer therapies [69].

Concordant with modulation of the host immune phenotype, the RNA-seq analysis also showed the upregulation of genes that encode both stimulatory and inhibitory immune checkpoint molecules, suggesting that these immune regulators affect infiltration of immune cells to the bladder and alter their activity. Immune checkpoint molecules expressed on the surface of immune cells (for example, CTLA-4 on T cells) regulate cell activity by transducing stimulatory or inhibitory signals upon ligation with cognate molecules (for example, CD80/B7-1 expressed on the surface of antigen presenting cells), and many cancer therapies target inhibitory immune checkpoint molecules to achieve T cell (re)activation and tumor cell killing (for example, by blocking CTLA-4 on T cells) [70]. Studies of immune checkpoint molecules during schistosome infection have mainly examined the inhibitory molecules and the effect of blocking their function. In the mouse *S. japonicum* infection model, the antibody-mediated blockade of CTLA- 4 resulted in increased granuloma size [71], suggesting that CTLA-4 signaling suppresses granuloma formation, likely by blocking T cell activation. The blockade of TIM-3 signaling in spleen lymphocytes from *S. japonicum* infected mice by treatment with soluble TIM-3-Fc fusion protein increased the population of CD4+IFNG+ Th1 cells [72], suggesting that TIM-3 signaling suppresses Th1 cell activity during infection. Similarly, another study found that liver CD4+ T cells, CD8+ T cells, NK1.1+ cells, and CD11b+ cells have increased TIM-3 expression during *S. japonicum* infection, and antibody-mediated blockade of TIM-3 on CD11b+ cells (from infected mice) co-cultured with soluble egg antigen (SEA) resulted in the upregulation of the classically activated (M1) macrophage-associated markers *Nos2* (iNOS synthase) and *Il12* and the downregulation of M2 macrophage-associated markers *Arg1* (liver arginase) and *Il10* [73]. A study examining PD-1 in the context of schistosome infection (reviewed in [74]) found an increase in PD-1 expression in the liver of *S. japonicum*-infected mice and that infected PD-1- deficient (versus infected wild-type) mice have a decreased population of cells that express IL-4 and IL-13 and an increased population of cells that express IFNG and FOXP3 [75], suggesting that PD-1 signaling supports a type 2 immune phenotype. Studies on PD-1 binding partners PD- L1 and PD-L2 found that splenic F4/80+ macrophages from mice infected with *S. mansoni* express elevated levels of PD-L1 and can induce anergy in T cells in a PD-L1-dependent manner [76], and that *S. japonicum* SEA can induce PD-L2 expression on bone marrow-derived dendritic cells (BMDCs) in a TLR2-dependent manner [77], suggesting that schistosome infection can alter PD-L1 and PD-L2 signaling to suppress T cell responses. Complementing these observations, the RNA-seq analysis showed the upregulation of several inhibitory immune checkpoint genes, including *Ctla4*, *Havcr2* (encodes TIM-3), *Cd274* (encodes PD-L1), and *Pdcd1lg2* (encodes PD-L2).

The upregulation of *Foxp3* and inhibitory immune checkpoint genes found in the RNA-seq analysis suggests that *S. haematobium* infection can induce an immunosuppressive host phenotype, indirectly benefiting the parasite by preventing excessive inflammation-related damage to the host [55]. The immunosuppressive phenotype induced by the infection may also favor the development of bladder cancer, as such a phenotype can allow aberrant, neoplastic cells to escape immune surveillance and destruction [78,79]. Concordantly, a study of the role of Treg cells (reviewed in [80,81]) in canine bladder cancer showed that CCR4 blockade-induced Treg depletion can inhibit tumor growth [82], and studies of immune checkpoint molecules (reviewed in [83,84]) found that high-grade bladder cancer expresses higher levels of PD-1 and PD-L1 compared to low-grade bladder cancer [85] and the inhibition of CTLA-4 alone or in combination with the inhibition of PD-1 prevents the growth of mouse bladder cancer [86].

While the mechanisms that drive *S. haematobium*-induced bladder cancer remain poorly understood, studies have suggested a possible carcinogenic role for schistosome-derived estrogen-like metabolites through adduct formation with DNA [87,88] (and reviewed in [89,90]). Other studies showed that total antigen protein extract of *S. haematobium* worms can antagonize estradiol-dependent activation of estrogen receptor expression and signaling [91,92], proposing that an estradiol-like molecule found in the extract likely exerts these effects [93] and that altered estrogen receptor signaling may also contribute to carcinogenesis. In contrast, a gene array analysis of urothelial cells found that co-culturing with *S. mansoni* eggs resulted in the upregulation of the estrogen receptor gene *Esr1*, while co-culturing with *S. haematobium* eggs showed a gene expression profile that suggests, by upstream regulatory analysis, downregulated estrogen receptor signaling activity [94]. The RNA-seq analysis of the egg-injected samples also showed contrasting results, with the upregulation of *Esr1* (*p*-adj = 0.125) and downregulation of *Esr2* (*p*-adj = 0.012) (Supplementary 5), warranting further investigation addressing the effect of infection on estrogen receptor signaling and its role in carcinogenesis.

In addition to estrogen receptor signaling, cell-proliferative signaling pathways altered by infection with *S. haematobium* may also contribute to carcinogenesis. The RNA-seq analysis of the egg-injected bladders revealed the enrichment of host genes related to increased KRAS signaling in the egg-injected samples. Aberrant constitutive KRAS signaling drives cell proliferation, and several types of cancer, including pancreatic, colorectal, and lung cancer, exhibit KRAS-activating mutations with high incidence [95–98]. Studies of Egyptian [99] and Chinese [100] colorectal cancer patients with schistosome infection found an association of KRAS mutation with schistosome infection, and while a study of bladder cancer patients with schistosome infection did not find an increased frequency of mutation in NRAS or KRAS, it found an unusually high frequency of a mutation in codon 13 of HRAS, another member of the RAS family [101,102]. Similarly, a study of bladders from mice intravesically exposed to total antigen protein extract of *S. haematobium* worms reported that bladders with dysplasia carried a G12D mutation in KRAS (a mutation frequently found in pancreatic adenocarcinoma [97]) with greater frequency compared to the control group [103]. Upregulated KRAS signaling, whether from activating mutations or increased upstream activity, represents a possible mechanism by which *S. haematobium* induces cell proliferation. Such proliferation has been observed in studies using Chinese hamster ovary (CHO) epithelial cells [51] and mouse bladders [104] exposed to *S. haematobium* worm antigen, in a worm antigen-exposed CHO cell xenograft model [105], J82 human and Endo bovine urothelial cells exposed to *S. haematobium* SEA [106], and HCV-29 human urothelial cells co-cultured with *S. haematobium* eggs [94].

The ability for KRAS signaling to increase cell proliferation may also extend to angiogenesis, given that KRAS promotes CXCL1, CXCL5, IL-8, VEGF, which can enhance human umbilical vein endothelial cell tube formation [107,108]. Consistent with the upregulation of KRAS signaling, the RNA-seq analysis showed the upregulated expression of *Cxcl1* and *Cxcl5* (Supplementary 5), as well as the enrichment of genes that belong to the angiogenesis gene set. In support of these observations, studies in *S. mansoni* showed that eggs and SEA can induce the proliferation of endothelial cells *in vitro* [109], and SEA can also induce endothelial cell tube formation and upregulation of vascular endothelial growth factor [110,111], with the caveat that host genetic background can greatly affect the extent of angiogenesis and other pathology [112]. Another study showed that *S. haematobium* IPSE/alpha-1, a major component of egg secretions, can induce hyperplasia of urothelial cells and angiogenesis when injected into the mouse bladder wall, as well as cell cycle polarization of urothelial cells and tube formation of endothelial cells *in vitro* [36]. We speculate that the eggs not only induce angiogenesis through the action of their secreted products, but also by recruiting host immune cells and forming granulomas, as similarly described in *M. tuberculosis* infection [113]. The type 2 immune response induced by the eggs [9] leads to the formation of granulomas containing M2 macrophages [114], which can contribute to angiogenesis and remodeling of the extracellular matrix [58]. In addition, as a granuloma recruits more cells, the cells closest to the core may experience hypoxia, which can induce proangiogenic signaling. Consistent with this mechanism for granuloma-induced hypoxia, the RNA-seq analysis showed the enrichment of genes belonging to the hypoxia gene set and the negative enrichment of genes belonging to the oxidative phosphorylation and fatty acid metabolism gene sets.

Egg granuloma formation and excretion from the host requires an intact T, and specifically Th2, cell response, suggesting that the granuloma and the activity of its constituent cells enable the intermediate steps of traversing endothelial cell, extracellular matrix (ECM), and epithelial cell barriers [9,115,116]. Matrix metalloproteinases (MMPs), which remodel the ECM by degrading adhesion, structural, and other related molecules [117], may contribute to egg-induced fibrosis and tissue traversal by the eggs [9]. The RNA-seq and microarray (week 1) analyses found the upregulation of the matrix metalloproteinases *Mmp3*, *Mmp10*, and *Mmp13*, while the RNA-seq analysis identified the upregulation of a broader set of MMPs (and related genes), including *Mmp8*, *Mmp9*, and *Timp1* (tissue inhibitor of metalloproteinases-1), consistent with the observations from studies of fibrosis-related genes in *S. mansoni*-infected mice, which found the upregulation of *Mmp8* and *Timp1* in the liver [118] and the upregulation of *Mmp2*, *Mmp3*, *Mmp8*, and *Timp1* in the colon [119]. The upregulation of *Timp1*, which can antagonize MMP activity, as well as modulate immune cell activity [120], may represent a balancing host response against the upregulation of MMPs. MMPs may also play a role in *S. haematobium* infection-related bladder cancer, given 1) the association of some MMPs with metastasis, invasion, and angiogenesis [121]; 2) that the RNA-seq analysis showed the enrichment of the epithelial-mesenchymal transition gene set, which includes several MMPs and genes that support metastasis; and 3) that the elevated expression of various MMPs correlates with poor prognosis and malignant progression of some types of cancer [122].

With respect to endothelial and epithelial cell barriers, the RNA-seq analysis, in agreement with previous analyses, showed the downregulation of several genes associated with cell adhesion. While not found in the microarray week 1 post-injection analysis, the microarray week 3 post-injection analysis showed the downregulation of genes related to urothelial barrier function (reviewed in [123]), such as the uroplakins *Upk1a*, *Upk1b*, *Upk2*, *Upk3a*, *Upk3b*; and *Cldn8*, *Cldn10*, *Cldn23*, and *Jam4* (*Igsf5*) [28], many with which the RNA-seq analysis showed concordance. The downregulation of these adhesion molecules may enhance the traversal of *S. haematobium* eggs across the urothelium and, together with increased angiogenesis, likely promotes hematuria. In contrast, the microarray, proteomic, and RNA-seq analyses all showed the upregulation of *Serpina3n* (serine protease inhibitor, clade A, member 3N), which can hasten the resolution of inflammation in a mouse model of colitis by binding and inhibiting elastase, a protein that enhances inflammation by degrading cell adhesion molecules [124]. Another study showed that the overexpression of *Serpina3n* can also alleviate cyclophosphamide-induced interstitial cystitis and inhibit apoptosis in the bladder by activating Wnt/β-catenin signaling [125], leading us to speculate that *Serpina3n* expression may represent a host strategy to counteract the disruption of cell barriers and to repair tissue damage caused by the immune response to *S. haematobium* eggs in the bladder.

The host gene expression profile in the egg-injected samples in the mouse bladder wall injection model suggests that the interactions between the host bladder and *S. haematobium* eggs, at 4 days post-injection, induce host immunomodulation of a complex nature, with the simultaneous upregulation of genes related to inflammation (chemokine signaling), type 1 immune responses (interferon and tumor necrosis factor signaling; associated with early infection), type 2 immune responses (*Il4* and *Il13*; associated with patency/egg-laying), and immunoregulatory responses (*Foxp3* and inhibitory immune checkpoint genes; associated with chronic infection). We note that because of the differences between the bladder wall injection model and natural infection, these observations may not accurately describe host responses during natural infection. However, consistent with the hallmarks of cancer [79], the altered host immune phenotype, together with the enrichment of genes related to KRAS signaling and the perturbation of genes involved in angiogenesis, ECM remodeling, cell adhesion, and tissue repair, can contribute to a carcinogenic microenvironment and may provide clues about possible mechanisms of *S. haematobium* egg traversal of host tissues and *S. haematobium*-related bladder cancer.

In addition to examining changes in gene expression in the mouse host bladder in response to the *S. haematobium* eggs, we aimed to characterize those in the eggs in response to the bladder environment. We found that the number of unique reads aligning to the *S. haematobium* genome represented a small proportion of the overall number of reads in egg-injected samples. Injecting more eggs in the hamster bladder wall and increasing the read depth did result in an overall greater number of unique reads aligning to the *S. haematobium* genome, but the proportion of the total number of reads did not increase when compared to the egg-injected mouse samples. This suggests that analyzing a microdissected section of the egg bolus(es) instead of the entire bladder could increase both the proportion and overall number of detected gene transcripts from the parasite. The differentially expressed *S. haematobium* genes in the comparison of egg-injected samples and egg (pooled egg and egg/bladder mix) samples included those likely involved in transcription, signaling, metabolism, protein (re)folding, ribosomal processes, and cell death, as well as uncharacterized genes. Changes in the expression of these genes may suggest that the *S. haematobium* eggs activate a specific transcriptional and phenotypic response to the bladder environment. Further investigation examining the gene transcription profile of *S. haematobium* eggs injected into the liver, intestine, and bladder could provide insight regarding whether eggs adopt unique phenotypes in different organs.

As noted in previous studies, the *S. haematobium* mouse bladder wall injection model currently provides the best approximation of *S. haematobium*-related bladder pathology, as natural infection of mice with *S. haematobium* cercariae produces only hepatointestinal pathology [24]. Likewise, the observations in this study provide an approximation of the changes in bladder biology that occur during natural human infection. The key differences of the mouse model include: 1) the one-time bolus egg delivery to the bladder wall, versus chronic, ongoing deposition of eggs in natural infection; 2) the surgery to access and inject eggs into the bladder wall, which, although used in conjunction with a vehicle injection control procedure, may produce artifactual changes not found in a natural infection; and 3) the absence of adult worms and associated host-parasite interactions. With respect to the first difference, the spatially and temporally acute nature of the egg delivery remains a limitation in the mouse bladder wall egg injection model, and future studies will need to identify the mechanisms controlling the tropism of *S. haematobium* parasites in the mammalian host to allow for the development of an experimental model that features continuous worm-based bladder oviposition. Such a model may also exhibit bladder tumorigenesis, allowing for more in-depth studies of *S. haematobium*- induced bladder cancer. Second, to minimize the effect of surgery, we are exploring the use of an ultrasound-guided bladder wall injection technique for egg injection, originally demonstrated for the injection of cells [128]. Finally, we previously found minimal immunological differences in the response to egg bladder wall injection when comparing naive mice and mice exposed to *S. haematobium* cercariae (natural infection) prior to injection [34], suggesting that the absence of adult worms in this study may have minimal effect on the bladder’s gene expression profile following bladder wall egg injection.

We also note some limitations of the current study. In addition to the need to omit some samples owing to technical difficulty in injecting eggs, several samples, notably egg-alone samples, did not generate RNA of sufficient quality to proceed to library preparation and sequencing (Supplementary 12). Using a greater number of parasite eggs (>6000 eggs) would more likely produce RNA of sufficient quality and quantity. In the bioinformatic analysis, we combined the data from two independent experiments (sample numbers 2xx and 3xx; Supplementary 12), as well as those from a reference sequence of *S. haematobium* eggs [43,44], which showed distinct clustering of each treatment group in the PCA; however, we cannot discount the possibility of errors introduced by batch-specific differences in the egg preparation and sequencing platform. Indeed, we chose not to combine a third independent experiment (sample numbers 1xx; Supplementary 12) in the analysis after noting dissimilarity in the PCA compared to the respective sample groups from the other two experiments, possibly because of the difference in sequencing platform.

Gene expression profiling of the mouse bladder wall *S. haematobium* egg injection model by RNA-seq in this study identified differentially expressed genes in the host bladder environment, revealing the activation of signaling pathways related to inflammatory response, immune cell chemotaxis, and cytokine signaling, consistent with previous studies. Simultaneously, other differentially expressed genes suggested the induction of host immune phenotypes and responses to dampen inflammatory responses, enhance egg tissue traversal, and promote a carcinogenic microenvironment. Questions remain regarding how different cell types, such as the bladder urothelium, endothelium, resident immune cells, and infiltrating immune cells respond to the parasite eggs and how these changes contribute to egg traversal of host tissues and carcinogenesis. While the current study analyzed gene expression differences across the entire bladder, which includes the genes and signals from infiltrating immune cells, further investigation using a single-cell RNA-seq approach would help to characterize cell type-specific responses to the eggs for both resident and infiltrating cells, providing a deeper understanding of *S. haematobium*-related bladder pathology and carcinogenesis. Characterizing the parasite egg gene profile in the bladder environment in comparison to the intestinal or liver environments remains a challenge in the bladder wall injection model, which the development of an experimental model for natural urogenital *S. haematobium* infection will help to address. Taken together, the results from this RNA-seq study highlight genes and pathways that improve our understanding of the pathology and carcinogenicity of *S. haematobium* infection and help pave the way for deeper mechanistic studies and development of improved diagnostics, vaccines, and new therapeutics.

## Data availability

Sequencing data collected for this report have been deposited at NCBI under BioProject accession identifier PRJNA1129449.

## Author contributions

Conceptualization (KI, MHH). Data curation (KI, DNMO, MR, ECM). Formal analysis (KI, DNMO, MR, ECM, MHH). Funding acquisition (MHH). Investigation (KI, OKL, ECM, JJC, LL). Methodology (all authors). Writing: original draft (KI). Writing: review and editing (all authors).

## Acknowledgments

This work was supported by NIH R01DK113504 (MHH) and George Washington University Cancer Biology Training Program NIH T32CA247756 (KI). We thank Taiwo Ogunbayo, Sarah Li, and Margaret Mentink-Kane of the NIAID Schistosome Resource Center at Biomedical Research Institute for graciously providing *S. haematobium* eggs (Rockville, MD) through NIH- NIAID contract HHSN272201700014I for distribution through BEI Resources. We thank Hiroki Morizono and Payal Banerjee of the Children’s National Research Institute (CNRI) Bioinformatics Unit, Claudio Anselmi of the CNRI Chief Research Information Office (CRIO), and Susan Knoblach of the CNRI Research Center for Genetic Medicine for providing computing resources and assistance.

## Supplementary information

[z01_mm-igfiltered.txt]

Supplementary 1. Genes related to immunoglobulin variable regions removed from the differentially expressed gene list.

[z02_bioinf_scripts.zip] 01_fastqc_master.sh 02_STAR_master.sh 03_featureCounts_master.sh 04_deseq2_fc_v8_dualtx_r2r3_clean.R 05_gsea_master.sh 06_gsea_rpt_cleanup_master.sh 07_panther_plot_master.R 08_ipa_dot_plot_master.R 09_extract_exon_idproduct_map_sh_master.sh

**Supplementary 2. Scripts used in the bioinformatics analysis.** [z03_bioinformatics_methods.pdf]

Supplementary 3. Detailed methods for the bioinformatics analysis.

[z04_sh_primers.txt]

Supplementary 4. Sequence information of the primers used in the measurement of parasite gene transcript levels.

[z05_mm-einj-vs-hinj-alpha0.05-diffexp-padj0.05-l2fc1-igfilt.txt]

Supplementary 5. Differentially expressed mouse genes in the comparison of egg-injected and vehicle-injected samples, sorted by log_2_ fold change in descending order.

[z06_mm-pca.png]

Supplementary 6. Principal component analysis of samples from three independent sets of experiments aligned to the mouse genome. Several egg-injected samples (121, 221-225, 321) did not receive eggs because of technical error and cluster with the vehicle-injected samples. Subsequent analyses consider samples from two sets of experiments (2xx and 3xx). Key: naive, no injection (soft-red filled circles); hinj, vehicle-injected (olive- yellow filled triangles); einj, *S. haematobium* egg-injected (cyan-green filled squares); egg, *S. haematobium* egg alone (blue crosses); eb, *S. haematobium* egg mixed with bladder tissue from mice that did not undergo bladder wall injection (soft-magenta open squares).

[z07_gsea_report_mh_combined_q005.txt]

Supplementary 7. Subset of top positively and negatively enriched gene sets from the mouse hallmark gene set collection in the comparison of egg-injected and vehicle-injected samples aligned to the mouse genome, arranged by normalized enrichment score. Key: Name, gene set collection; Size, gene set size; ES, enrichment score; NES, normalized enrichment score; NOM p-val, nominal *p*-value; FDR q-val, false discovery rate likelihood; FWER p-val, family-wise Rank at max, rank of gene from the ranked gene list at peak of running enrichment score; Leading edge, tags, proportion of gene hits (matches to the genes in the gene set) to ranked gene list before (for positive enrichment scores) or after (for negative enrichment scores) the peak of the running enrichment score; Leading edge, list, proportion of ranked genes before (for positive enrichment scores) or after (for negative enrichment scores) the peak of the running enrichment score; Leading edge, signal, composite score of the “tags” and “list” leading edge attributes.

[z08_panther_output_upreg_bp.txt]

Supplementary 8. Gene ontology (biological process) analysis results from the PANTHER Overrepresentation Test for upregulated differentially expressed genes in the comparison of egg-injected and vehicle-injected samples.

[z09_ipa_signaling_pathways.xls]

Supplementary 9. Pathways, z-scores, log_10_ *p*-values, and associated genes from the “Signaling Pathways” analysis of Ingenuity Pathways Analysis using differentially expressed mouse genes in the comparison of egg- injected and vehicle-injected samples.

[z10_mm-einj-vs-hinj-alpha0.05-diffexp-rnaseq-microarraywk1-prot.txt]

Supplementary 10. Differentially expressed mouse genes or proteins (in the comparison of egg-injected against vehicle-injected samples) shared among the RNA-seq, microarray (week 1 data), and proteomics studies. Note that for the microarray data, for genes with multiple entries, one representative entry is shown. Key for each column: “gene_symb”, gene symbol associated with the gene accession identifier; “ensembl_acc_rnaseq”, Ensembl database accession identifier; “gene_desc”, gene description associated with the gene accession identifier; “l2fc_rnaseq”, log_2_ fold change from the RNA-seq data; “padj_rnaseq”, adjusted *p*-value from the RNA-seq data; “l2fc_microarraywk1”, log_2_ fold change derived from the week 1 microarray data; “uniprot_acc_proteomics”, UniProt database accession identifier from the proteomics data; “l2fc_proteomics”, log_2_ fold change from the proteomics data; “padj_proteomics”, adjusted *p*-value from the proteomics data.

[z11_panther_output_rm-a_bp.txt]

Supplementary 11. Gene ontology (biological process) analysis results from the PANTHER Overrepresentation Test for the shared differentially expressed genes (in the comparison of egg-injected and vehicle-injected samples) between the RNA-seq analysis in this study and microarray (week 1 data) analysis, sorted by false discovery rate (FDR) in ascending order.

[z12_sample_info_all.xlsx]

Supplementary 12. Overview of samples and associated information. Sample naming convention: hundreds’ place: experiment set or run; tens’ place: treatment group such that 0 = naive/no surgery (naive), 1 = vehicle control injection (hinj), 2 = egg injection (einj), 3 = egg-alone (egg), 4 = egg mixed with bladder tissue from mice that did not undergo bladder wall injection (eb); ones’ place: biological replicate. For example, sample 321 corresponds to set 3, egg injection treatment, replicate 1. Analyses presented in this report use pooled data from set 2 (samples 2xx) and set 3 (samples 3xx).

[z13_sh_pca_r31.tif]

Supplementary 13. Principal component analysis of samples from two independent sets of experiments aligned to the *S. haematobium* genome including egg reference data. Key: naive, no injection (soft-red filled circles); hinj, vehicle-injected (olive-yellow filled triangles); einj, *S. haematobium* egg-injected (cyan-green filled squares); egg, *S. haematobium* egg alone (blue crosses); eb, *S. haematobium* egg mixed with bladder tissue from mice that did not undergo bladder wall injection (soft-magenta open squares). A) all sample groups. B) egg-injected (einj), egg-alone (egg), and egg mixed with bladder tissue from mice that did not undergo injection (eb); eb samples are pooled together with egg samples.

[z14_sh-einj-vs-egg-alpha0.05-diffexp-padj0.05-l2fc1.txt]

Supplementary 14. Differentially expressed *S. haematobium* genes in the comparison of egg-injected (einj) against egg-alone (pooled egg and eb) samples, sorted by adjusted *p*-value in ascending order. Each row represents a gene. Columns represent: Gene identifier (NCBI SchHae_2.0), mean count, log_2_ fold change, log_2_ fold change error, Wald statistic, *p*-value, adjusted *p*-value, gene description.

[z15_qpcr.png]

Supplementary 15. Gene transcript expression level for *S. haematobium Ipse/alpha-1* and *Cox1* across 0 to 4 days following bladder wall injection determined by quantitative PCR of cDNA, expressed in cycle threshold (C_T_) values. Low cycle numbers indicate more expression, while high cycle numbers indicate less expression. Key: D, days post bladder wall injection; NTC, no-template control.

`GroupNameShort GroupNameLong BladderWallInjection RNA_source naive no surgery No Mouse bladder`

`hinj vehicle control bladder wall injection Yes Mouse bladder, 4 days after injection with hamster liver-intestine homogenate`

`einj egg bladder wall injection Yes Mouse bladder, 4 days after injection with S. haematobium eggs`

`egg egg alone N/A S. haematobium eggs`

`eb egg mixed with bladder tissue from mice that did not undergo bladder wall injection No Mix of naive mouse bladder and S. haematobium eggs`

`Gene name Gene description log2FoldChange padj`

`Ly6g lymphocyte antigen 6 complex, locus G 3.59 5.72e-16 Ccl4 chemokine (C-C motif) ligand 4 3.53 1.94e-37 Hcar2 hydroxycarboxylic acid receptor 2 3.45 2.58e-30`

`Ccl3 chemokine (C-C motif) ligand 3 3.33 2.00e-19 Mrgpra2a MAS-related GPR, member A2A 3.26 4.37e-14 Slfn4 schlafen 4 3.25 5.68e-30`

`Trim30b tripartite motif-containing 30B 3.16 4.35e-22 Nos2 nitric oxide synthase 2, inducible 3.15 7.86e-29 Gm43181 predicted gene 43181 3.15 6.02e-13`

`Rsad2 radical S-adenosyl methionine domain containing 2 3.11 2.10e-19 Pigr polymeric immunoglobulin receptor -1.72 4.08e-03`

`Hrh3 histamine receptor H3 -1.72 1.19e-07`

`Cxcl13 chemokine (C-X-C motif) ligand 13 -1.73 6.54e-10 2310002F09Rik RIKEN cDNA 2310002F09 gene -1.74 1.07e-07`

`Fam180a family with sequence similarity 180, member A -1.76 7.54e-11 Cryaa crystallin, alpha A -1.89 2.70e-08`

`Myoc myocilin -2.13 1.14e-11`

`Ttr transthyretin -2.15 6.29e-15`

`Rsph4a radial spoke head 4 homolog A (Chlamydomonas) -2.25 6.87e-19 Lyz1 lysozyme 1 -2.91 3.88e-42`

`Gene ID Gene description log2FoldChange padj MS3_0014405 KRAB-A domain-containing protein 4.03 1.18e-14 MS3_0018165 putative annexin 3.09 1.78e-12`

MS3_0014185 putative dice1/dead/h box polypeptide 2.49 8.77e-16 MS3_0019193 glycine dehydrogenase (decarboxylating) 2.34 2.34e-04 MS3_0015312 putative glutamate receptor, kainate 2.28 7.36e-04 MS3_0019209 tyrosine kinase 2.20 2.08e-05

MS3_0019439 serine/threonine kinase 2.13 7.74e-06 MS3_0012086 diacylglycerol O-acyltransferase 1 2.10 3.23e-03 MS3_0019576 putative heat shock protein 70 2.07 5.59e-75 MS3_0014056 protein kinase 2.05 7.78e-06

`MS3_0016935 isoleucine--tRNA ligase -2.05 4.77e-02`

MS3_0014817 putative 22.6kDa tegument-associated antigen -2.05 4.51e-02 MS3_0014925 putative atp:guanidino kinase (Smc74) -2.06 2.08e-05 MS3_0013388 putative alpha-1, 3-mannosyl-glycoprotein beta-1, 2-n- acetylglucosaminyltransferase -2.07 4.73e-02

`MS3_0012108 putative small nuclear ribonucleoprotein SM-D3 -2.12 3.89e-02`

`MS3_0017143 putative programmed cell death protein -2.17 4.73e-02 MS3_0012787 rho GDP-dissociation inhibitor-related protein -2.40`

`1.17e-02`

`MS3_0019446 putative fibrocystin l -2.52 6.87e-03 MS3_0017626 polycystin 1-related protein -2.59 5.36e-03 MS3_0014818 putative antigen Sm21.7 -2.63 3.54e-03`

