## Supplementary 2 for "RNA-seq gene expression profiling of the bladder in a mouse model of urogenital schistosomiasis": 01_fastqc_master.sh

```
# What this script does
  # Executes FastQC (version 0.11.9) to generate summary reports of sequence read quality

# How to use this script
  # Save a copy
  # Customize path/file variables (replace empty strings ""), other settings
  # Save
  # Feed script to bash (e.g., bash customizedscript.sh)

# Software
  # Official repository (requires a suitable Java Runtime Environment; see installation notes)
    # https://www.bioinformatics.babraham.ac.uk/projects/fastqc/
  # The conda package for FastQC 0.11.9 (2020-01-24) is also available on bioconda (all platforms)
    # conda install -c bioconda fastqc=0.11.9=0

# Installation, setup, usage
  # https://raw.githubusercontent.com/s-andrews/FastQC/master/INSTALL.txt

# Make sure the FastQC executable is on $PATH (Java must also be on $PATH)
  # Download and extract the FastQC package, then give the path of the directory that contains the FastQC executable and add it to $PATH
    # path_fastqc="" # path of the directory that contains the FastQC executable
    # export PATH="${PATH}:${path_fastqc}" # append the path of the directory that contains the FastQC executable to the $PATH variable
  # Alternative: conda (activating the environment will add the locations of the installed package executables to $PATH)
    conda activate "" # activate the conda env that contains the FastQC installation

# Set up path and file locations
  pathproj="" # path for project
  pathdata="" # path for input reads data, e.g. "${pathproj}/data"
  pathout="" # path for writing data, e.g., "${pathproj}/outfastqc"
  mkdir -p "${pathout}" # create the output path if it doesn't exist

# Create a bash array and populate it with path/file of reads
  filelist=()
  while read -r -d '' line; do
    filelist+=("${line}")
    done \
    < <(find "${pathdata}" -type f -name "*.fq.gz" -print0)

# Execute FastQC
fastqc \
  --threads 4 \
  --outdir="${pathout}" \
  "${filelist[@]}" \
  2>&1 \
  | \
  tee "${pathout}/fastqc.log"
```
