## Supplementary 2 for "RNA-seq gene expression profiling of the bladder in a mouse model of urogenital schistosomiasis": 02_STAR_master.sh

```
# What this script does
  # Executes STAR (version 2.7.3a) functions for genome index generation and alignment/mapping of reads to a genome

# How to use this script
  # Save a copy
  # Customize path/file variables (replace empty strings ""), other settings
  # Save
  # Feed script to bash (e.g., bash customizedscript.sh)

# Software
  # Official repository (requires compilation; see installation notes)
    # https://github.com/alexdobin/STAR/tree/2.7.3a
  # The conda package for STAR 2.7.3a (2019-10-08) is also available on bioconda for Linux (and OSX)
    # conda install -c bioconda star=2.7.3a=0

# Manual
  # https://github.com/alexdobin/STAR/blob/2.7.3a/doc/STARmanual.pdf

# Make sure the STAR executable is on $PATH
  # If STAR provided by module, load the module (module will add the location of the STAR executable location to $PATH automatically)
    module load STAR
  # Alternative 1: download/compile, then give the path of the directory that contains the STAR executable and add it to $PATH
    # path_star="" # path of the directory that contains the STAR executable
    # export PATH="${PATH}:${path_star}" # append the path of the directory that contains the STAR executable to the $PATH variable
  # Alternative 2: conda (activating the environment will add the locations of the installed package executables to $PATH)
    # conda activate "" # activate the conda env that contains the STAR installation

# Set up path and file locations
  pathref="" # full path/filename for reference genome indexed by STAR # No index? See below
  pathproj="" # path for project
  pathdata="" # path for input reads data
  pathout="" # path for writing data, e.g., "${pathproj}/output"
  mkdir -p "${pathout}" # create the output path if it doesn't exist
  outSTAR="" # path for writing data (STAR will write output files here), e.g., "${pathout}/star"
  tempSTAR="" # genomeLoad functions put file here, e.g., "${pathout}/star-temp"
  mkdir -p "${outSTAR}"
  mkdir -p "${tempSTAR}"
  cd "${tempSTAR}"

# If no index, generate it (see manual)
  # STAR should be on $PATH
  if [ ! -d "${pathref}/" ]; then # checks for the absence (non-existence) of the directory that contains index files
    mkdir -p "${pathref}" # create the directory where STAR should write index files
    # genomeFastaFiles: string; /path/to/fastaforward and (if paired-end) /path/to/fastareverse
    # sjdbGTFfile: string; /path/to/gtffile
    # sjdbOverhang: integer; ideally max(readlength - 1); default value 100
    STAR \
      --runThreadN 4 \
      --runMode genomeGenerate \
      --genomeDir "${pathref}" \
      --genomeFastaFiles "" "" \
      --sjdbGTFfile "" \
      --sjdbOverhang 100
    fi

# Execute STAR
  # STAR should be on $PATH
  # Use STAR's LoadAndExit function to load genome and keep in memory
    STAR --genomeLoad LoadAndExit --genomeDir "${pathref}"
  # Use find to identify (read 1) target files, then use while-read loop/block to provide path/file info to STAR
    # Alternative to using find: provide a txt file containing a list of full path/file of reads files to the while block by redirecting (<) the txt file into the done statement (e.g. done < listofreadsfiles.txt)
    # Assumes paired-end reads with naming _1.fq.gz and _2.fq.gz
      # For single-end reads, delete the second argument passed to --readFilesIn
    # Adjust accordingly: filenames may end with .fastq.gz, .fgz, .fq.gz, etc
    # runThreadN: number of CPUs to use
    # genomeLoad LoadAndKeep: loads genome index and keeps it in memory
    find "${pathdata}" -type f -name "*_1.fq.gz" -print0 \
      | \
      while read -r -d '' pf; do
        fileread="${pf##*/}"
        fileread2="${fileread/_1.fq.gz/}"
        STAR \
          --runThreadN 4 \
          --genomeDir "${pathref}" \
          --genomeLoad LoadAndKeep \
          --readFilesCommand gunzip -c \
          --readFilesIn "${pf}" "${pf/_1.fq.gz/_2.fq.gz}" \
          --outFileNamePrefix "${outSTAR}/${fileread2}." \
          --outSAMtype BAM Unsorted \
          --outSAMunmapped Within \
          --outSAMattributes Standard
        done

# Unload the genome from memory
  STAR --genomeLoad Remove --genomeDir "${pathref}"
```
