## Supplementary 2 for "RNA-seq gene expression profiling of the bladder in a mouse model of urogenital schistosomiasis": 03_featureCounts_master.sh

```
# What this script does
  # Executes featureCounts from the subread package (version 2.0.0) for reads counting

# How to use this script
  # Save a copy
  # Customize path/file variables (replace empty strings ""), other settings
  # Save
  # Feed script to bash (e.g., bash customizedscript.sh)

# Software
  # Official package (Linux)
    # https://sourceforge.net/projects/subread/files/subread-2.0.0/subread-2.0.0-Linux-x86_64.tar.gz/download
  # The conda package for subread 2.0.0 (2020-02-04) is also available on bioconda for Linux (and OSX)
    # conda install -c bioconda subread=2.0.0=hed695b0_0

# Manual
  # https://subread.sourceforge.net/featureCounts.html

# Make sure the featureCounts executable is on $PATH
  # Download and extract the subread package, then give the path of the directory that contains the featureCounts executable and add it to $PATH
    path_fc="" # path of the directory that contains the featureCounts executable
    export PATH="${PATH}:${path_fc}" # append the path of the directory that contains the featureCounts executable to the $PATH variable
  # Alternative: conda (activating the environment will add the locations of the installed package executables to $PATH)
    # conda activate "" # activate the conda env that contains the subread installation

# Set up path and file locations
  gtf="" # gtf file
  pathproj="" # path for project
  pathbamfiles="" # path of the directory that contains .bam files of mapped reads, e.g., "${pathproj}/star"
  outfc="" # path for writing data (featureCounts will write output files here), e.g., "${pathproj}/fc"
  mkdir -p "${outfc}"

# Execute featureCounts
  # For paired-end reads (-p switch)
    featureCounts \
      -T 4 \
      -p \
      -t exon \
      -g gene_id \
      -a "${gtf}" \
      -o "${outfc}/counts.txt" \
      "${pathbamfiles}"/*.bam \
      2>"${outfc}/fc.err"
  # The above ("${pathbamfiles}"/*.bam) assumes that all .bam files reside in the given path
    # To select specific .bam files, replace this with path/file strings, e.g. "/path/to/bam1" "/path/to/bam2", each separated by a space

  # For single-end reads
    # featureCounts \
      # -T 4 \
      # -t exon \
      # -g gene_id \
      # -a "${gtf}" \
      # -o "${outfc}/counts.txt" \
      # "${pathbamfiles}"/*.bam \
      # 2>"${outfc}/fc.err"
```
