## Supplementary 2 for "RNA-seq gene expression profiling of the bladder in a mouse model of urogenital schistosomiasis": 04_deseq2_fc_v8_dualtx_r2r3_clean.R

```
# R script that takes read counts files from featureCounts and runs DESeq2
	# Make a copy of this script
	# Customize path/file variables (replace applicable empty strings "") and other settings
		# pathproj, idmap_mm_file, idmap_sh_file, mhset_file, microarray_file, proteomics_file, pathout, pathcounts1, pathcounts2, pathcounts3, pathcountsref
	# Execute the customized script using Rscript under conda env deseq2-126
	# Feed script to Rscript (e.g., Rscript customizedscript.R)

# Parts of script taken from https://gist.github.com/stephenturner/f60c1934405c127f09a6

# Environment
	# conda 4.8.3
	# r 3.6.1 h3a67422_6 (2019-07-05)(bundled with bioconda deseq2)
	# deseq2 1.26.0 r36he1b5a44_0 (2019-11-03)(bioconda)
	# enhancedvolcano 1.4.0 r36_0 (2019-11-01)(bioconda) (modified code)
	# ggplot2 3.2.1 r36h6115d3f_0 (2019-08-12)(bundled with bioconda deseq2)
	# ggrepel 0.8.1 r36h0357c0b_1 (2019-07-18)(conda-forge)
	# r-venndiagram 1.6.20 r36h6115d3f_1002 (2020-05-04)(conda-forge)
	# r-readxl 1.3.1 r36h0357c0b_1 (2019-07-18)(conda-forge)

# Input files
	# Non-normalized counts (counts.txt) from e.g., featureCounts
	# Gene accession, description, symbol mapping files for mouse/mm (ENSEMBL) and S. haematobium/sh (NCBI)
	# GSEA MSigDB mouse hallmark gene set collection
	# Proteomics and microarray (week 1) differential expression data

# Output files
	# analysis-igfiltered.txt (mm only)
	# analysis-omit-pool-normcounts.txt
	# analysis-omit-pool-gsea_sample_class.cls (mm only)
	# analysis-omit-pool-normcounts_gsea.txt (mm only)
	# analysis-omit-pool-pca.png
	# analysis-omit-pool-alpha*-summary.txt
	# analysis-omit-pool-alpha*-diffexp.txt
	# analysis-omit-pool-alpha*-diffexp-padj*-l2fc*.txt
	# analysis-omit-pool-alpha*-diffexp-padj*-l2fc*-igfilt.txt (mm only)
	# analysis-omit-pool-alpha*-diffexp-padj*-l2fc*-igfilt-id-l2fc-padj.txt (mm only)
	# analysis-omit-pool-alpha*-diffexp-padj*-l2fc*(-igfilt)-topgenes_l2fc.txt
	# analysis-omit-pool-alpha*-diffexp-padj*-l2fc*(-igfilt)-topgenes_padj.txt
	# analysis-omit-pool-alpha*-diffexp-padj*-l2fc*(-igfilt)-enhvolcano.png
	# analysis-omit-pool-alpha*-venn.tiff (mm only)
	# analysis-omit-pool-alpha*-venn_numbers.txt (mm only)
# Key for output files
	# analysis = "mm", "sh"
	# omit = "omitn", "omity", "exclusive"
	# pool = "pooln", "pooly"
	# *=user-defined cutoff value; value for alpha matches value for padj
	# alpha*, padj*: * = myalpha
	# l2fc*: * = l2fccut

# Sample numbering code
	# hundreds place: run
		# possible values: 1, 2, 3
	# tens place: treatment group
		# possible values: 0, 1, 2, 3, 4
		# 0: naive
		# 1: hinj
		# 2: einj
		# 3: egg
		# 4: eb
	# ones place: sample number within treatment group
		# possible values: 1, 2, 3, 4, 5
	# example: 312 means run 3, hinj, sample 2

# DESeq2 convention for DESeq::results(dds, contrast=c("condition", "einj", "egg") ... )
	#  einj vs egg = exp vs ctrl

# Project root directory
	pathproj <- ""
	setwd(pathproj)

# Load libraries
	library("DESeq2")
	library("ggplot2")
	library("ggrepel")
	library("data.table")
	library("readxl")
	library("VennDiagram")

# See after mainfunction() block for analysis

# EnhancedVolcanoModified() function (modified from EnhancedVolcano 1.4.0)
EnhancedVolcanoModified <- function (
	toptable,
	lab,
	x,
	y,
	selectLab = NULL,
	xlim = c(min(toptable[[x]], na.rm = TRUE),
	max(toptable[[x]], na.rm = TRUE)),
	ylim = c(0, max(-log10(toptable[[y]]), na.rm = TRUE) + 5),
	xlab = bquote(~Log[2] ~ "fold change"),
	ylab = bquote(~-Log[10] ~ italic(P)),
	axisLabSize = 18,
	title = "Volcano plot",
	subtitle = "EnhancedVolcano",
	caption = paste0("Total = ", nrow(toptable), " variables"),
	titleLabSize = 18,
	subtitleLabSize = 14,
	captionLabSize = 14,
	pCutoff = 1e-05,
	pLabellingCutoff = pCutoff,
	FCcutoff = 1,
	cutoffLineType = "longdash",
	cutoffLineCol = "black",
	cutoffLineWidth = 0.4,
	transcriptPointSize = 0.8,
	transcriptLabSize = 3,
	transcriptLabCol = "black",
	transcriptLabFace = "plain",
	transcriptLabhjust = 0,
	transcriptLabvjust = 1.5,
	pointSize = 2,
	labSize = 3,
	labCol = "black",
	labFace = "plain",
	labhjust = 0,
	labvjust = 1.5,
	boxedlabels = FALSE,
	boxedLabels = FALSE,
	shape = 19,
	shapeCustom = NULL,
	col = c("grey30", "forestgreen", "royalblue", "red2"),
	colCustom = NULL,
	colAlpha = 1/2,
	legend = c("NS", "Log2 FC", "P", "P & Log2 FC"),
	legendLabels = c("NS", expression(Log[2] ~ FC), "p-value", expression(p - value ~ and ~ log[2] ~ FC)),
	legendPosition = "top",
	legendLabSize = 14,
	legendIconSize = 4,
	legendVisible = TRUE,
	shade = NULL,
	shadeLabel = NULL,
	shadeAlpha = 1/2,
	shadeFill = "grey",
	shadeSize = 0.01,
	shadeBins = 2,
	drawconnectors = FALSE,
	drawConnectors = FALSE,
	widthConnectors = 0.5,
	typeConnectors = "closed",
	endsConnectors = "first",
	lengthConnectors = unit(0.01, "npc"),
	colConnectors = "grey10",
	hline = NULL,
	hlineType = "longdash",
	hlineCol = "black",
	hlineWidth = 0.4,
	vline = NULL,
	vlineType = "longdash",
	vlineCol = "black",
	vlineWidth = 0.4,
	gridlines.major = TRUE,
	gridlines.minor = TRUE,
	border = "partial",
	borderWidth = 0.8,
	borderColour = "black") 
	{
		if (!is.numeric(toptable[[x]])) {
			stop(paste(x, " is not numeric!", sep = ""))
		}
		if (!is.numeric(toptable[[y]])) {
			stop(paste(y, " is not numeric!", sep = ""))
		}
		i <- xvals <- yvals <- Sig <- NULL
		if (!missing("transcriptPointSize")) {
			warning(paste0("transcriptPointSize argument deprecated in v1.4", 
				" - please use pointSize"))
			pointSize <- transcriptPointSize
		}
		if (!missing("transcriptLabSize")) {
			warning(paste0("transcriptLabSize argument deprecated in v1.4", 
				" - please use labSize"))
			labSize <- transcriptLabSize
		}
		if (!missing("transcriptLabCol")) {
			warning(paste0("transcriptLabCol argument deprecated in v1.4", 
				" - please use labCol"))
			labCol <- transcriptLabCol
		}
		if (!missing("transcriptLabFace")) {
			warning(paste0("transcriptLabFace argument deprecated in v1.4", 
				" - please use labFace"))
			labFace <- transcriptLabFace
		}
		if (!missing("transcriptLabhjust")) {
			warning(paste0("transcriptLabhjust argument deprecated in v1.4", 
				" - please use labhjust"))
			labhjust <- transcriptLabhjust
		}
		if (!missing("transcriptLabvjust")) {
			warning(paste0("transcriptLabvjust argument deprecated in v1.4", 
				" - please use labvjust"))
			labvjust <- transcriptLabvjust
		}
		if (!missing("boxedlabels")) {
			warning(paste0("boxedlabels argument deprecated in v1.4", 
				" - please use boxedLabels"))
			boxedLabels <- boxedlabels
		}
		if (!missing("drawconnectors")) {
			warning(paste0("drawconnectors argument deprecated since v1.2", 
				" - please use drawConnectors"))
			drawConnectors <- drawconnectors
		}
		toptable <- as.data.frame(toptable)
		toptable$Sig <- "NS"
		toptable$Sig[(abs(toptable[[x]]) > FCcutoff)] <- "FC"
		toptable$Sig[(toptable[[y]] < pCutoff)] <- "P"
		toptable$Sig[(toptable[[y]] < pCutoff) & (abs(toptable[[x]]) > 
			FCcutoff)] <- "FC_P"
		toptable$Sig <- factor(toptable$Sig, levels = c("NS", "FC", 
			"P", "FC_P"))
		if (min(toptable[[y]], na.rm = TRUE) == 0) {
			warning(paste("One or more p-values is 0.", "Converting to 10^-1 * current", 
				"lowest non-zero p-value..."), call. = FALSE)
			toptable[which(toptable[[y]] == 0), y] <- min(toptable[which(toptable[[y]] != 
				0), y], na.rm = TRUE) * 10^-1
		}
		toptable$lab <- lab
		toptable$xvals <- toptable[[x]]
		toptable$yvals <- toptable[[y]]
		if (!is.null(selectLab)) {
			names.new <- rep(NA, length(toptable$lab))
			indices <- which(toptable$lab %in% selectLab)
			names.new[indices] <- toptable$lab[indices]
			toptable$lab <- names.new
		}
		th <- theme_bw(base_size = 24) + theme(legend.background = element_rect(), 
			plot.title = element_text(angle = 0, size = titleLabSize, 
				face = "bold", vjust = 1), plot.subtitle = element_text(angle = 0, 
				size = subtitleLabSize, face = "plain", vjust = 1), 
			plot.caption = element_text(angle = 0, size = captionLabSize, 
				face = "plain", vjust = 1), axis.text.x = element_text(angle = 0, 
				size = axisLabSize, vjust = 1), axis.text.y = element_text(angle = 0, 
				size = axisLabSize, vjust = 1), axis.title = element_text(size = axisLabSize), 
			legend.position = legendPosition, legend.key = element_blank(), 
			legend.key.size = unit(0.5, "cm"), legend.text = element_text(size = legendLabSize), 
			title = element_text(size = legendLabSize), legend.title = element_blank())
		if (!is.null(colCustom) & !is.null(shapeCustom)) {
			plot <- ggplot(toptable, aes(x = xvals, y = -log10(yvals))) + 
				th + guides(colour = guide_legend(order = 1, override.aes = list(size = legendIconSize)), 
				shape = guide_legend(order = 2, override.aes = list(size = legendIconSize))) + 
				geom_point(aes(color = factor(names(colCustom)), 
					shape = factor(names(shapeCustom))), alpha = colAlpha, 
					size = pointSize, na.rm = TRUE) + scale_color_manual(values = colCustom) + 
				scale_shape_manual(values = shapeCustom)
		}
		else if (!is.null(colCustom) & is.null(shapeCustom) & length(shape) == 
			1) {
			plot <- ggplot(toptable, aes(x = xvals, y = -log10(yvals))) + 
				th + guides(colour = guide_legend(order = 1, override.aes = list(size = legendIconSize)), 
				shape = guide_legend(order = 2, override.aes = list(size = legendIconSize))) + 
				geom_point(aes(color = factor(names(colCustom))), 
					alpha = colAlpha, shape = shape, size = pointSize, 
					na.rm = TRUE) + scale_color_manual(values = colCustom) + 
				scale_shape_manual(guide = TRUE)
		}
		else if (!is.null(colCustom) & is.null(shapeCustom) & length(shape) == 
			4) {
			plot <- ggplot(toptable, aes(x = xvals, y = -log10(yvals))) + 
				th + guides(colour = guide_legend(order = 1, override.aes = list(size = legendIconSize)), 
				shape = guide_legend(order = 2, override.aes = list(size = legendIconSize))) + 
				geom_point(aes(color = factor(names(colCustom)), 
					shape = factor(Sig)), alpha = colAlpha, size = pointSize, 
					na.rm = TRUE) + scale_color_manual(values = colCustom) + 
				scale_shape_manual(values = c(NS = shape[1], FC = shape[2], 
					P = shape[3], FC_P = shape[4]), labels = c(NS = legendLabels[1], 
					FC = legendLabels[2], P = legendLabels[3], FC_P = legendLabels[4]), 
					guide = TRUE)
		}
		else if (is.null(colCustom) & !is.null(shapeCustom)) {
			plot <- ggplot(toptable, aes(x = xvals, y = -log10(yvals))) + 
				th + guides(colour = guide_legend(order = 1, override.aes = list(size = legendIconSize)), 
				shape = guide_legend(order = 2, override.aes = list(size = legendIconSize))) + 
				geom_point(aes(color = factor(Sig), shape = factor(names(shapeCustom))), 
					alpha = colAlpha, size = pointSize, na.rm = TRUE) + 
				scale_color_manual(values = c(NS = col[1], FC = col[2], 
					P = col[3], FC_P = col[4]), labels = c(NS = legendLabels[1], 
					FC = legendLabels[2], P = legendLabels[3], FC_P = legendLabels[4])) + 
				scale_shape_manual(values = shapeCustom)
		}
		else if (is.null(colCustom) & is.null(shapeCustom) & length(shape) == 
			1) {
			plot <- ggplot(toptable, aes(x = xvals, y = -log10(yvals))) + 
				th + guides(colour = guide_legend(order = 1, override.aes = list(shape = shape, 
				size = legendIconSize))) + geom_point(aes(color = factor(Sig)), 
				alpha = colAlpha, shape = shape, size = pointSize, 
				na.rm = TRUE, show.legend = legendVisible) + scale_color_manual(values = c(NS = col[1], 
				FC = col[2], P = col[3], FC_P = col[4]), labels = c(NS = legendLabels[1], 
				FC = legendLabels[2], P = legendLabels[3], FC_P = legendLabels[4]))
		}
		else if (is.null(colCustom) & is.null(shapeCustom) & length(shape) == 
			4) {
			plot <- ggplot(toptable, aes(x = xvals, y = -log10(yvals))) + 
				th + guides(colour = guide_legend(order = 1, override.aes = list(shape = c(NS = shape[1], 
				FC = shape[2], P = shape[3], FC_P = shape[4]), size = legendIconSize))) + 
				geom_point(aes(color = factor(Sig), shape = factor(Sig)), 
					alpha = colAlpha, size = pointSize, na.rm = TRUE, 
					show.legend = legendVisible) + scale_color_manual(values = c(NS = col[1], 
				FC = col[2], P = col[3], FC_P = col[4]), labels = c(NS = legendLabels[1], 
				FC = legendLabels[2], P = legendLabels[3], FC_P = legendLabels[4])) + 
				scale_shape_manual(values = c(NS = shape[1], FC = shape[2], 
					P = shape[3], FC_P = shape[4]), guide = FALSE)
		}
		plot <- plot + xlab(xlab) + ylab(ylab) + xlim(xlim[1], xlim[2]) + 
			ylim(ylim[1], ylim[2]) + geom_vline(xintercept = c(-FCcutoff, 
			FCcutoff), linetype = cutoffLineType, colour = cutoffLineCol, 
			size = cutoffLineWidth) + geom_hline(yintercept = -log10(pCutoff), 
			linetype = cutoffLineType, colour = cutoffLineCol, size = cutoffLineWidth)
		plot <- plot + labs(title = title, subtitle = subtitle, caption = caption)
		if (!is.null(vline)) {
			plot <- plot + geom_vline(xintercept = vline, linetype = vlineType, 
				colour = vlineCol, size = vlineWidth)
		}
		if (!is.null(hline)) {
			plot <- plot + geom_hline(yintercept = -log10(hline), 
				linetype = hlineType, colour = hlineCol, size = hlineWidth)
		}
		if (border == "full") {
			plot <- plot + theme(panel.border = element_rect(colour = borderColour, 
				fill = NA, size = borderWidth))
		}
		else if (border == "partial") {
			plot <- plot + theme(axis.line = element_line(size = borderWidth, 
				colour = borderColour), panel.border = element_blank(), 
				panel.background = element_blank())
		}
		else {
			stop("Unrecognised value passed to 'border'. Must be 'full' or 'partial'")
		}
		if (gridlines.major == TRUE) {
			plot <- plot + theme(panel.grid.major = element_line())
		}
		else {
			plot <- plot + theme(panel.grid.major = element_blank())
		}
		if (gridlines.minor == TRUE) {
			plot <- plot + theme(panel.grid.minor = element_line())
		}
		else {
			plot <- plot + theme(panel.grid.minor = element_blank())
		}
		if (boxedLabels == FALSE) {
			if (drawConnectors == TRUE && is.null(selectLab)) {
				plot <- plot + geom_text_repel(data = subset(toptable, 
					toptable[[y]] < pLabellingCutoff & abs(toptable[[x]]) > 
					  FCcutoff), aes(label = subset(toptable, toptable[[y]] < 
					pLabellingCutoff & abs(toptable[[x]]) > FCcutoff)[["lab"]]), 
					size = labSize, segment.color = colConnectors, 
					segment.size = widthConnectors,
					# arrow = arrow(
						# length = lengthConnectors,
						# type = typeConnectors,
						# ends = endsConnectors),
					hjust = labhjust, vjust = labvjust, colour = labCol, 
					fontface = labFace, na.rm = TRUE)
			}
			else if (drawConnectors == TRUE && !is.null(selectLab)) {
				plot <- plot + geom_text_repel(data = subset(toptable, 
					!is.na(toptable[["lab"]])), aes(label = subset(toptable, 
					!is.na(toptable[["lab"]]))[["lab"]]), size = labSize, 
					segment.color = colConnectors, segment.size = widthConnectors, 
					# arrow = arrow(
						# length = lengthConnectors,
						# type = typeConnectors,
						# ends = endsConnectors),
					hjust = labhjust, vjust = labvjust, 
					colour = labCol, fontface = labFace, na.rm = TRUE)
			}
			else if (drawConnectors == FALSE && !is.null(selectLab)) {
				plot <- plot + geom_text(data = subset(toptable, 
					!is.na(toptable[["lab"]])), aes(label = subset(toptable, 
					!is.na(toptable[["lab"]]))[["lab"]]), size = labSize, 
					check_overlap = TRUE, hjust = labhjust, vjust = labvjust, 
					colour = labCol, fontface = labFace, na.rm = TRUE)
			}
			else if (drawConnectors == FALSE && is.null(selectLab)) {
				plot <- plot + geom_text(data = subset(toptable, 
					toptable[[y]] < pLabellingCutoff & abs(toptable[[x]]) > 
					  FCcutoff), aes(label = subset(toptable, toptable[[y]] < 
					pLabellingCutoff & abs(toptable[[x]]) > FCcutoff)[["lab"]]), 
					size = labSize, check_overlap = TRUE, hjust = labhjust, 
					vjust = labvjust, colour = labCol, fontface = labFace, 
					na.rm = TRUE)
			}
		}
		else {
			if (drawConnectors == TRUE && is.null(selectLab)) {
				plot <- plot + geom_label_repel(data = subset(toptable, 
					toptable[[y]] < pLabellingCutoff & abs(toptable[[x]]) > 
					  FCcutoff), aes(label = subset(toptable, toptable[[y]] < 
					pLabellingCutoff & abs(toptable[[x]]) > FCcutoff)[["lab"]]), 
					size = labSize, segment.color = colConnectors, 
					segment.size = widthConnectors, arrow = arrow(length = lengthConnectors, 
					  type = typeConnectors, ends = endsConnectors), 
					hjust = labhjust, vjust = labvjust, colour = labCol, 
					fontface = labFace, na.rm = TRUE)
			}
			else if (drawConnectors == TRUE && !is.null(selectLab)) {
				plot <- plot + geom_label_repel(data = subset(toptable, 
					!is.na(toptable[["lab"]])), aes(label = subset(toptable, 
					!is.na(toptable[["lab"]]))[["lab"]]), size = labSize, 
					segment.color = colConnectors, segment.size = widthConnectors, 
					arrow = arrow(length = lengthConnectors, type = typeConnectors, 
					  ends = endsConnectors), hjust = labhjust, vjust = labvjust, 
					colour = labCol, fontface = labFace, na.rm = TRUE)
			}
			else if (drawConnectors == FALSE && !is.null(selectLab)) {
				plot <- plot + geom_label(data = subset(toptable, 
					!is.na(toptable[["lab"]])), aes(label = subset(toptable, 
					!is.na(toptable[["lab"]]))[["lab"]]), size = labSize, 
					hjust = labhjust, vjust = labvjust, colour = labCol, 
					fontface = labFace, na.rm = TRUE)
			}
			else if (drawConnectors == FALSE && is.null(selectLab)) {
				plot <- plot + geom_label(data = subset(toptable, 
					toptable[[y]] < pLabellingCutoff & abs(toptable[[x]]) > 
					  FCcutoff), aes(label = subset(toptable, toptable[[y]] < 
					pLabellingCutoff & abs(toptable[[x]]) > FCcutoff)[["lab"]]), 
					size = labSize, hjust = labhjust, vjust = labvjust, 
					colour = labCol, fontface = labFace, na.rm = TRUE)
			}
		}
		if (!is.null(shade)) {
			plot <- plot + stat_density2d(data = subset(toptable, 
				rownames(toptable) %in% shade), fill = shadeFill, 
				alpha = shadeAlpha, geom = "polygon", contour = TRUE, 
				size = shadeSize, bins = shadeBins, show.legend = FALSE, 
				na.rm = TRUE) + scale_fill_identity(name = shadeLabel, 
				labels = shadeLabel, guide = "legend")
		}
		return(plot)
	}


# Define mainfunction
mainfunction <- function (pathproj, analysis, omit_samples, pool_eggeb, anum, includefirst, shref){

	# Project root directory
		pathproj <- pathproj # set when this function is called
		setwd(pathproj)
	# Other file locations (specify full path/file)
		idmap_mm_file <- "" # Ensembl BioMart file containing 3 columns: gene accessions, names/symbols, and descriptions (has header)
		idmap_sh_file <- "" # Mapping file for shv2 genome extracted from gtf file containing 2 columns: gene ids and descriptions (does not have header)
		mhset_file <- "" # MSigDB mouse hallmark gene set collection
		microarray_file <- "" # From egg bladder wall injection microarray study; week 1 data extracted manually from file "https://doi.org/10.1371/journal.pntd.0001912.s001"
		proteomics_file <- "" # From egg bladder wall injection proteomics study; directly from file "https://doi.org/10.1371/journal.pntd.0010176.s011"
	# Directory for output of analysis
		if (analysis == "mm"){
			pathout <- paste(pathproj, sep="", "/mm/output/", anum, "-deseq2-", omit_samples, "-", pool_eggeb, "-includefirst", includefirst, "-shref", shref)
			}
		if (analysis == "sh"){
			pathout <- paste(pathproj, sep="", "/shv2/output/", anum, "-deseq2-", omit_samples, "-", pool_eggeb, "-includefirst", includefirst, "-shref", shref)
			}
		if (dir.exists(pathout)==TRUE){
			message(paste0("Analysis destination directory \"", pathout, "\" already exists. Skipping."))
			return(NULL)
			}
		if (dir.exists(pathout)==FALSE){
			dir.create(pathout, recursive=TRUE)
			message(paste0("Writing to analysis destination directory \"", pathout, "\"."))
			}

	# Location of counts files
		if (analysis == "mm"){
			pathcounts1 <- paste(pathproj, sep="", "") # data from run 1 (mm)
			pathcounts2 <- paste(pathproj, sep="", "") # data from run 2 (mm)
			pathcounts3 <- paste(pathproj, sep="", "") # data from run 3 (mm)
			}
		if (analysis == "sh"){
			pathcounts2 <- paste(pathproj, sep="", "") # data from run 2 (sh)
			pathcounts3 <- paste(pathproj, sep="", "") # data from run 3 (sh)
			pathcountsref <- "" # data from eggref (sh)
			}


	message("reading in counts table...", appendLF=FALSE)
	# Read data from file
		if (includefirst){
			countdata1 <- read.table(file.path(pathcounts1, "counts.txt"), header=TRUE, row.names=1, check.names=FALSE)
			countdata1 <- countdata1[ ,6:ncol(countdata1)]
			}
		countdata2 <- read.table(file.path(pathcounts2, "counts.txt"), header=TRUE, row.names=1, check.names=FALSE)
		countdata2 <- countdata2[ ,6:ncol(countdata2)]
		countdata3 <- read.table(file.path(pathcounts3, "counts.txt"), header=TRUE, row.names=1, check.names=FALSE)
		countdata3 <- countdata3[ ,6:ncol(countdata3)]
		if (shref){
			countdataref <- read.table(file.path(pathcountsref, "counts.txt"), header=TRUE, row.names=1, check.names=FALSE)
			countdataref <- countdataref[ ,6, drop = FALSE]
			}

	# Combine/merge countdata
		# using match() and cbind()
			countdata_all <- cbind( countdata2, countdata3[match(rownames(countdata2), rownames(countdata3)), ] ) # 2 and 3
			if (shref){
				countdata_all <- cbind( countdata_all, countdataref[match(rownames(countdata_all), rownames(countdataref)), , drop=FALSE] ) # (2+3)+ref
				}
			if (includefirst){
				countdata_all <- cbind( countdata_all, countdata1[match(rownames(countdata_all), rownames(countdata1)), ] ) # add set 1 to the data frame
				}

	# Rename samples to standardized format see "sample numbering code"
		# Refer to example sample names below for substitution:
			# /data/kishida/rnaseq/dualtx/100/outSTAR/101.fq.gz.Aligned.out.bam
			# /data/kishida/rnaseq/dualtx/200/outSTAR/201_1.fq.gz.Aligned.out.bam
			# /data/kishida/rnaseq/dualtx/Clean/outSTAR/301.Aligned.out.bam
			# /data/kishida/rnaseq/dualtx/200/output/2-shv2/21-star/201.Aligned.out.bam
			# /data/kishida/rnaseq/dualtx/Clean/shv2/outSTAR/301.Aligned.out.bam
			# /data/kishida/rnaseq/dualtx/stroehlein2019/egg/outSTAR/SRR6655493.Aligned.out.bam
		colnames(countdata_all) <- sub("^.*/", "", colnames(countdata_all)) # remove path from names
		colnames(countdata_all) <- sub("^([0-9A-Z]+).*", "\\1", colnames(countdata_all)) # isolate 3-number sample name
		colnames(countdata_all) <- sub("SRR6655493", "R31", colnames(countdata_all))

	# Remove samples
		# 121, 221-225, 321 einj (failed to receive eggs)
		if (omit_samples == "omity" | omit_samples == "exclusive"){
			countdata_all <- countdata_all[,-grep("121|22[1-9]|321",colnames(countdata_all))] #removes 121, 221-225, 321
			}

	# Remove more samples for exclusive analysis
		if (analysis == "mm" & omit_samples == "exclusive"){
			countdata_all <- countdata_all[, -grep(".[034].", colnames(countdata_all))]
			} # removes naive, egg, and eb samples
		if (analysis == "sh" & omit_samples == "exclusive"){
			countdata_all <- countdata_all[, -grep(".[01].", colnames(countdata_all))]
			} # removes naive and hinj samples

	# Remove/filter-out genes (rows) for immunoglobulins except for Igha, Ighd, Ighe, Ighg, Ighm (only for analysis "mm")
		if (analysis == "mm"){
			# Read-in Ensembl accession mapping file
				idmap_mm_df <- read.table(idmap_mm_file, header=TRUE, check.names=FALSE, sep="\t", quote="")
				# sep="\t" required because headers and gene descriptions contain spaces
				# quote="" required because of names like 3'-terminal phosphate cyclase
			# Select Ensembl accessions for immunoglobulin genes except for Igha, Ighd, Ighe, Ighg, Ighm
				igfilter_idx <- grep("Ig(k|h(v|d[0-9]-[0-9]|j)|l[a-z][0-9])", idmap_mm_df$"Gene name") # obtain index numbers
				igfilter_df <- idmap_mm_df[igfilter_idx, ] # select records using index
				write.table(
					igfilter_df,
					file=paste0(pathout, "/", analysis, "-igfiltered.txt"),
					sep="\t",
					quote=F,
					row.names=F) # write selected records to file
				igfilter_vec <- as.character(igfilter_df$"Gene stable ID") # select the gene accessions
				# igfilter_vec <- as.character(idmap_mm_df$"Gene stable ID"[grep("Ig(k|h(v|d[0-9]-[0-9]|j)|l[a-z][0-9])", idmap_mm_df$"Gene name")]) # does the same thing as above
			}
	# Read-in gene ids for sh here, similar to mm case
		if (analysis == "sh"){
				idmap_sh_df <- read.table(idmap_sh_file, header=FALSE, check.names=FALSE, sep="\t", quote="")
			}

	# Assign treatments to each sample following the "sample numbering code"
		countdata_all <- as.matrix(countdata_all) # convert data.frame to matrix
		condition <- factor(substr(colnames(countdata_all),2,2)) # extract treatment group number from sample number
		if (pool_eggeb == "pooln"){
			# to keep eb and egg as separate groups
			code_assign <- list(
				c("0","naive"),
				c("1","hinj"),
				c("2","einj"),
				c("3","egg"),
				c("4","eb")
				)
			}
		if (pool_eggeb == "pooly"){
			# to group eb together with egg
			code_assign <- list(
				c("0","naive"),
				c("1","hinj"),
				c("2","einj"),
				c("3","egg"),
				c("4","egg")
				)
			}
		cond_lev <- levels(condition) # levels() returns a character vector
		for(i in 1:length(code_assign)){
			cond_lev <- sub(code_assign[[i]][1], code_assign[[i]][2], cond_lev)
			} # loop through code_assign to substitute values in cond_lev
		levels(condition) <- cond_lev # rename the values of condition from numeric to treatment name
		coldata <- data.frame(row.names=colnames(countdata_all), condition) # assign samples to conditions in this data.frame
		message("done", appendLF=TRUE)

	# DESeq2 function for combining countdata and sample-treatment association
		message("DESeq2 dds...", appendLF=FALSE)
		dds <- DESeqDataSetFromMatrix(countData=countdata_all, colData=coldata, design=~condition)
		dds <- DESeq(dds)
		message("done", appendLF=TRUE)

	# Normalized counts
		message("writing normcounts...", appendLF=FALSE)
		normcounts <- counts(dds, normalized=TRUE)
		write.table(normcounts, file=paste0(pathout, "/", analysis, "-normcounts.txt"), sep="\t", quote=F, col.names=NA)
		message("done", appendLF=TRUE)

	# Prepare files for input into GSEA (mm only)
		if (analysis == "mm"){
			# GSEA file prep (part 1 of 3): samples reordering: GSEA requires contiguous groups for sample groups
				# naive naive naive hinj hinj einj # Correct
					# 101 102 201 111 211 121
				# naive naive hinj einj naive hinj # Incorrect
					# 101 102 111 121 201 211
				# The order of samples in the normalized counts file must match
				# Extract positions
					position1 <- substr(colnames(countdata_all), 1, 1) # first position (run)
					position2 <- substr(colnames(countdata_all), 2, 2) # second position (treatment)
					position3 <- substr(colnames(countdata_all), 3, 3) # third position (replicate)
				# Sorting on multiple conditions: sort on treatment, run, replicate
					samplesortindex <- order(position2, position1, position3) # returns the indexes
				# Use the sort index to sort the values
					# Using the same sort index for condition and normcounts guarantees matching order
					condition_sorted <- condition[samplesortindex]
					normcounts_sorted <- normcounts[, samplesortindex]

				# # Sorting sample order example
					# # 212 102 104 113 114 203 204 211
					# myvec1 <- c("212", "102", "104", "113", "114", "203", "204", "211")
					# order(myvec1) # Outputs index numbers of target corresponding to sort: 2 3 4 5 6 7 8 1
					# myvec1[order(myvec1)] # Use index numbers to obtain values of each index
					# # Desired order
						# # 102 104 203 204 113 114 211 212
						# # Index numbers 2 3 6 7 4 5 8 1
						# # Sort order: middleposition, firstposition, lastposition
						# # Extract each position
							# myvec1_first <- substr(myvec1,1,1)
							# myvec1_mid <- substr(myvec1,2,2)
							# myvec1_last <- substr(myvec1,3,3)
						# # Apply sorting sequentially (sort lastposition, firstposition, middleposition)?
							# # Sorting applied from least important key to most important key
								# # myvec1 <- myvec1[order(myvec1_last)] # 211 212 102 113 203 104 114 204
								# # myvec1 <- myvec1[order(myvec1_first)] # 102 113 104 114 211 212 203 204
								# # myvec1 <- myvec1[order(myvec1_mid)] # 102 104 203 204 113 114 211 212
							# # Problem: doesn't provide index numbers that we can apply to the original
						# # Apply multiple sorting with order()
							# mysortindex <- order(myvec1_mid, myvec1_first, myvec1_last)
							# myvec1[mysortindex]

			# GSEA file prep (part 2 of 3): class (.cls) file
				fx <- file(paste0(pathout, "/", analysis, "-gsea_sample_class.cls"))
					writeLines(
						text=c(
							paste(length(condition), length(levels(condition)), "1", sep=" "),
							paste("#", paste(levels(condition), collapse=" "), sep=" "),
							paste(as.character(condition[samplesortindex]), collapse=" ")
							),
						con=fx
						)
				close(fx)

			# GSEA file prep (part 3 of 3): GSEA-formatted normcounts file
				# Create new data.frame containing the required format
					normcountsnew <- data.frame(
						NAME=row.names(normcounts),
						DESCRIPTION="na",
						row.names=row.names(normcounts)
						)
				# Paste together the sorted normcounts and write to file
					normcountsnew <- cbind(normcountsnew, normcounts[, samplesortindex])
					write.table(
						normcountsnew,
						row.names=FALSE,
						file=paste0(pathout, "/", analysis, "-normcounts_gsea.txt"),
						sep="\t",
						quote=F)
		}


	# Log transform the deseq data set (use in PCA, heatmap)
		message("rlogTransformation...", appendLF=FALSE)
		logdds <- rlogTransformation(dds)
		message("done", appendLF=TRUE)


	# PCA
		message("pca plot...", appendLF=FALSE)

		if (analysis == "mm"){
			chart_title <- bquote(Alignment~to~mouse~genome)
			}
		if (analysis == "sh"){
			chart_title <- bquote(Alignment~to~italic(S.~haematobium)~genome)
			}
		if (omit_samples == "omitn"){
			pca_label <- factor(colnames(logdds))
			}
		if (omit_samples == "omity" | omit_samples == "exclusive"){
			pca_label <- factor(gsub("[0-9]{3}", "", colnames(logdds)))
			}

		# Statically assign colors and shapes to each treatment group
			group_colors <- c(
				naive="#F8766D",
				hinj="#A3A500",
				einj="#00BF7D",
				egg="#00B0F6",
				eb="#E76BF3"
				)
			group_shapes <- c(
				naive=16,
				hinj=17,
				einj=15,
				egg=3,
				eb=7
				)

		pcaplot <- DESeq2::plotPCA(logdds,intgroup="condition") +
			geom_text_repel(
				aes_string(
					label=pca_label
					#x="PC1", y="PC2", label=factor(paste0(logdds$condition,":",colnames(logdds)))
					#x="PC1", y="PC2", label=factor(colnames(logdds))
					#x="PC1", y="PC2", label=factor(gsub("[0-9]{3}", "", colnames(logdds)))
					) ,
				point.padding = 0.1,
				size=10,
				show.legend=F
				) +
			geom_point(size=6) +
			aes_string(color="group", shape="group") +
			labs(title=bquote(paste(PCA:.(chart_title))),
				subtitle=" ") +
			scale_color_manual(values=group_colors) +
			scale_shape_manual(values=group_shapes) +
			guides(color=guide_legend(title="")) +
			guides(shape=guide_legend(title="")) +
			# scale_color_discrete(name="newlegendtitle") +
			# scale_shape_discrete(name="newlegendtitle") +
			theme_bw() +
			theme(
				title=element_text(size=20),
				axis.text=element_text(size=20),
				axis.title=element_text(size=20),
				legend.text=element_text(size=20),
				#legend.title=element_text(size=20),
				aspect.ratio=1
				)
		png(paste0(pathout, "/", analysis, "-pca.png"),1000,1000,pointsize=20)
		print(pcaplot)
		dev.off()
		message("done", appendLF=TRUE)


	# Results by DESeq2
		# Set alpha; should use same value as intended p-adj cutoff
			# https://hbctraining.github.io/DGE_workshop_salmon_online/lessons/05b_wald_test_results.html
			myalpha <- 0.05
		# Set a cutoff for log2FoldChange (l2fc)
			l2fccut <- 1
		# Select treatment groups to compare (pairwise)
			if (analysis == "mm"){
				cond_exp <- "einj"
				cond_ctrl <- "hinj"
				}
			if (analysis == "sh"){
				cond_exp <- "einj"
				cond_ctrl <- "egg"
					# note that "egg" here refers to pooled egg and eb, depending on the value of the code_assign variable
				}
		# Execute results function
			res <- results(dds, contrast=c("condition", cond_exp, cond_ctrl), independentFiltering=TRUE, alpha=myalpha, pAdjustMethod="BH", parallel=TRUE)
		# Execute lfcShrink function
			res <- lfcShrink(dds, contrast=c("condition", cond_exp, cond_ctrl), res=res)
		# Arrange/sort results dataframe by padj values
			res <- res[order(res$padj), ]
		# Store analysis info in this string
			filenamestring <- paste0(analysis, "-", cond_exp, "-vs-", cond_ctrl, "-alpha", myalpha)
		# Write summary and diffexp files
			sink(paste0(pathout, "/", filenamestring, "-summary.txt")); summary(res); sink()
			write.table(res, file=paste0(pathout, "/", filenamestring, "-diffexp.txt"), sep="\t", quote=F, col.names=F)

	# Process results (l2fc and padj) and write tables
		# Select l2fc and padj; map ids; apply cutoff; extract top genes; apply formatting; write file
			res_map <- data.frame(res[c("log2FoldChange","padj")])
		# Map ids
			if (analysis == "mm"){
				res_map <- cbind(res_map, idmap_mm_df[match(rownames(res_map), idmap_mm_df$"Gene stable ID"), ])
				}
			if (analysis == "sh"){
				res_map <- cbind(res_map, idmap_sh_df[match(rownames(res_map), idmap_sh_df$"V1"), ])
				}
		# Sort by l2fc
			res_map_sort <- res_map[order(-res_map$"log2FoldChange"), ]
		# Apply l2fc cutoff (use l2fccut value)
			res_map_sort_cut <- subset(res_map_sort, res_map_sort$"padj" < myalpha & abs(res_map_sort$"log2FoldChange") > l2fccut)
		# For mm only: filter out Ig variable genes and write both unfiltered and filtered files
			if (analysis == "mm"){
				# Select columns for writing (unfiltered)
					res_map_sort_cut_out <- res_map_sort_cut[c("Gene stable ID", "Gene name", "Gene description", "log2FoldChange", "padj")]
				# Write the file (unfiltered)
					write.table(res_map_sort_cut_out, file=paste0(pathout, "/", filenamestring, "-diffexp-padj", myalpha, "-l2fc", l2fccut, ".txt"), sep="\t", quote=F, row.names=F)

				# Apply Ig filtering
					res_map_sort_filt <- res_map_sort[!(res_map_sort$"Gene stable ID" %in% igfilter_vec),]
				# Apply l2fc and padj cutoff to Ig-filtered list
					res_map_sort_filt_cut <- subset(res_map_sort_filt, res_map_sort_filt$"padj" < myalpha & abs(res_map_sort_filt$"log2FoldChange") > l2fccut)
				# Select columns for writing (Ig-filtered)
					res_map_sort_filt_cut_out <- res_map_sort_filt_cut[c("Gene stable ID", "Gene name", "Gene description", "log2FoldChange", "padj")]
				# Write the file (Ig-filtered)
					write.table(res_map_sort_filt_cut_out, file=paste0(pathout, "/", filenamestring, "-diffexp-padj", myalpha, "-l2fc", l2fccut, "-igfilt.txt"), sep="\t", quote=F, row.names=F)
				# Write a file with IDs, l2fc, padj (Ig-filtered)
					write.table(res_map_sort_filt_cut_out[c("Gene stable ID", "log2FoldChange", "padj")], file=paste0(pathout, "/", filenamestring, "-diffexp-padj", myalpha, "-l2fc", l2fccut, "-igfilt-id-l2fc-padj.txt"), sep="\t", quote=F, row.names=F)
				}
			if (analysis == "sh"){
				# Select columns for writing and name them appropriately
					res_map_sort_cut_out <- res_map_sort_cut[c("V1", "V2", "log2FoldChange", "padj")]
					colnames(res_map_sort_cut_out) <- c("Gene ID", "Gene description", "log2FoldChange", "padj")
				# Write the file
					write.table(res_map_sort_cut_out, file=paste0(pathout, "/", filenamestring, "-diffexp-padj", myalpha, "-l2fc", l2fccut, ".txt"), sep="\t", quote=F, row.names=F)
				}
		# Top genes by: l2fc, padj (for mm, post-igfilt)
			if (analysis == "mm"){
				# Create separate data.frames for differentially expressed genes with positive and negative log2FoldChange
					respos <- subset(res_map_sort_filt_cut, res_map_sort_filt_cut$"log2FoldChange" > l2fccut)
					resneg <- subset(res_map_sort_filt_cut, res_map_sort_filt_cut$"log2FoldChange" < -l2fccut)
					# Write a file with IDs, l2fc, padj (Ig-filtered, only l2fc > l2fccut)
						write.table(respos[c("Gene stable ID", "log2FoldChange", "padj")], file=paste0(pathout, "/", filenamestring, "-diffexp-padj", myalpha, "-l2fc", l2fccut, "-igfilt-id-l2fc-padj-pos.txt"), sep="\t", quote=F, row.names=F)
					# Write a file with IDs, l2fc, padj (Ig-filtered, only l2fc < -l2fccut)
						write.table(resneg[c("Gene stable ID", "log2FoldChange", "padj")], file=paste0(pathout, "/", filenamestring, "-diffexp-padj", myalpha, "-l2fc", l2fccut, "-igfilt-id-l2fc-padj-neg.txt"), sep="\t", quote=F, row.names=F)
				# Top/bottom by l2fc
					restop_l2fc <- rbind(head(respos, n=10), tail(resneg, n=10))
					restop_l2fc <- restop_l2fc[c("Gene name", "Gene description", "log2FoldChange", "padj")]
					restop_l2fc$"Gene description" <- sub(" \\[.*\\]$", "", restop_l2fc$"Gene description")
				# Top by padj
					restop_padj <- head( res_map_sort_filt_cut[order(res_map_sort_filt_cut$"padj"), ], n=20 )
					restop_padj <- restop_padj[c("Gene name", "Gene description", "log2FoldChange", "padj")]
					restop_padj$"Gene description" <- sub(" \\[.*\\]$", "", restop_padj$"Gene description")
				# Set this string for file naming purposes
					igfilter_str <- "-igfilt"
				}
			if (analysis == "sh"){
				# Remove records for which "Gene description" has value "uncharacterized protein"
					res_map_sort_cut <- subset(res_map_sort_cut, res_map_sort_cut$"V2" != "uncharacterized protein")
				# Create separate data.frames for differentially expressed genes with positive and negative log2FoldChange
					respos <- subset(res_map_sort_cut, res_map_sort_cut$"log2FoldChange" > l2fccut)
					resneg <- subset(res_map_sort_cut, res_map_sort_cut$"log2FoldChange" < -l2fccut)
				# Top/bottom by l2fc
					restop_l2fc <- rbind(head(respos, n=10), tail(resneg, n=10))
					restop_l2fc <- restop_l2fc[c("V1", "V2", "log2FoldChange", "padj")]
					colnames(restop_l2fc) <- c("Gene ID", "Gene description", "log2FoldChange", "padj")
				# Top by padj
					restop_padj <- head( res_map_sort_cut[order(res_map_sort_cut$"padj"), ], n=20 )
					restop_padj <- restop_padj[c("V1", "V2", "log2FoldChange", "padj")]
					colnames(restop_padj) <- c("Gene ID", "Gene description", "log2FoldChange", "padj")
				# Set this string for file naming purposes
					igfilter_str <- NULL
				}
			# Tidy the l2fc and padj (2 digits after decimal)
				restop_l2fc_out <- restop_l2fc
				restop_l2fc_out$"log2FoldChange" <- sapply(restop_l2fc_out$"log2FoldChange", formatC, format="f", digits=2)
				restop_l2fc_out$"padj" <- sapply(restop_l2fc_out$"padj", formatC, format="e", digits=2)
				restop_padj_out <- restop_padj
				restop_padj_out$"log2FoldChange" <- sapply(restop_padj_out$"log2FoldChange", formatC, format="f", digits=2)
				restop_padj_out$"padj" <- sapply(restop_padj_out$"padj", formatC, format="e", digits=2)
			# Write files for top/bottom l2fc and top padj
				write.table(restop_l2fc_out, file=paste0(pathout, "/", filenamestring, "-diffexp-padj", myalpha, "-l2fc", l2fccut, igfilter_str, "-topgenes_l2fc.txt"), sep="\t", quote=F, row.names=F)
				write.table(restop_padj_out, file=paste0(pathout, "/", filenamestring, "-diffexp-padj", myalpha, "-l2fc", l2fccut, igfilter_str, "-topgenes_padj.txt"), sep="\t", quote=F, row.names=F)
			# Message
				message("DESeq2 results done")


	# Volcano plot
		# Symbols have printing issues if underlying shell env has set LC_ALL=C
			# LC_ALL should have null value or "en_US.UTF-8"
			# locale command (in bash) will list language locale env variables
		# Check locale values in R using below
			# sessionInfo()
		# Check shell variables from R env using below
			# Sys.getenv()
		# Set locale from R env using below, assuming R uses the LANG variable
			# Sys.setenv(LANG = "en_US.UTF-8")

		# EnhancedVolcano package
			# Vignette:
				# https://www.bioconductor.org/packages/release/bioc/vignettes/EnhancedVolcano/inst/doc/EnhancedVolcano.html
			# Bioconductor:
				# https://bioconductor.org/packages/release/bioc/html/EnhancedVolcano.html
			# Github:
				# https://github.com/kevinblighe/EnhancedVolcano
			# Repo @ Anaconda/bioconda/packages
				# https://anaconda.org/bioconda/bioconductor-enhancedvolcano
		# Paths
			pathout <- pathout # set previously
		# Load the library (loaded previously)
			# library("EnhancedVolcano")
			# library("ggrepel")
		# Set cutoffs and create subsets of result
			# For reference, fold change and log2 fold change
				# 2-fold = 1
				# 1.5-fold = 0.585
				# 1.25-fold = 0.322
		# Previously defined values
			# res # DESeq2 result object
			# filenamestring # string for file naming
			# myalpha # DESeq2 independent filtering value
			# chart_title # set in PCA section
			# myalpha # set in section for DESeq2
			# l2fccut # set in section for DESeq2

		if (analysis == "mm"){
			# To label points on volcano, need to map gene names
				# res_df <- as.data.frame(res) # res is a deseq2 results object; convert to data.frame
				# resmap_df <- merge(x=res_df, y=idmap_mm_df, by.x="row.names", by.y="Gene stable ID") # mapping with this line
					# Check new mapped column names
						# colnames(resmap_df)
						# Note that we can now access gene symbols from resmap_df$"Gene name"
				# resd <- res # reassign original res object to another name
				# res <- resmap_df # overwrite res object with mapped
				res_orig <- res # store original results here
				res <- res_map_sort_filt # overwrite res with mapped, sorted, igfiltered dataframe
			# We want to select a subset of genes that belong to a gene set of choice
				# Start with reading in gene set collection (.gmt)
				mhset_file <- mhset_file # set previously
					# The .gmt file has no header and is tab-delimited
					# Each row contains: name of gene set, url link, followed by a variable number of gene set members
					# Importantly, not all rows have the same number of columns, so need to count before reading-in as df
						mhset_maxnumofcols <- max(count.fields(mhset_file, sep = "\t"))
				mhset_df <- read.table(mhset_file, header=FALSE, check.names=FALSE, sep="\t", quote="", fill=TRUE, col.names = paste0("V",seq_len(mhset_maxnumofcols)))
				mhset_inflam_idx <- grep("HALLMARK_INFLAMMATORY_RESPONSE", mhset_df[,1]) # get the index
				mhset_inflamm_vec <- as.character(unlist(mhset_df[mhset_inflam_idx, ])) # get the row using [idx, ] notation (gives a data.frame object), use unlist() to convert data.frame to factor, then as.character() to convert factor to character
				mhset_inflamm_vec_nonblank <- mhset_inflamm_vec[ nzchar(mhset_inflamm_vec) ] # remove blanks
				mhset_inflamm_genes <- mhset_inflamm_vec_nonblank[3:length(mhset_inflamm_vec_nonblank)] # vector of gene names
				res_select <- res[res$"Gene name" %in% mhset_inflamm_genes, ]
				res_subselect <- subset(res_select, res_select$"padj" < myalpha & abs(res_select$"log2FoldChange") > l2fccut)
				select_genes <- as.character(res_subselect$"Gene name")
				volcano_gene_labs <- as.character(res$"Gene name")
			}
		if (analysis == "sh"){
			# Subset of the top l2fc for labeling on volcanoplot
				restop_l2fc_vec <- c(
					as.character(head(restop_l2fc, n=5)$"Gene ID"),
					as.character(tail(restop_l2fc, n=5)$"Gene ID")
					)
				select_genes <- restop_l2fc_vec # for volcano plot
			# Gene labels for volcanoplot
				volcano_gene_labs <- rownames(res)
			}


		# cutoff for p value
			# pcut <- 0.1
			pcut <- myalpha # use the alpha value chosen for DESeq2::results
		# cutoff for log2FoldChange
			l2fccut <- l2fccut # defined previously
		# total number of genes
			totn <- nrow(res)
		# total number of genes not "NA" = plotted
			plotn <- nrow(subset(res, !is.na(res$padj) & !is.na(res$log2FoldChange)))
		# number of genes below pcut
			pcutn <- nrow(na.omit(subset(res, res$padj < pcut)))
		# number of genes greater than absolute l2fc cutoff
			l2fccutn <- nrow(na.omit(subset(res, res$log2FoldChange > l2fccut | res$log2FoldChange < -l2fccut)))
		# number of genes less than pcut
			pl2fccutn <- nrow(na.omit(subset(res, res$padj < pcut & (res$log2FoldChange > l2fccut | res$log2FoldChange < -l2fccut))))
		# number of genes less than pcut and not greater than absolute l2fc cutoff
			# pcutn - pl2fccutn == nrow(na.omit(subset(res, res$padj < pcut & !(res$log2FoldChange > l2fccut | res$log2FoldChange < -l2fccut))))
		# number of genes greater or equal to pcut and greater than absolute l2fc cutoff
			# l2fccutn - pl2fccutn == nrow(na.omit(subset(res, res$padj >= pcut & (res$log2FoldChange > l2fccut | res$log2FoldChange < -l2fccut))))
		# number of genes not meeting padj or l2fc cutoffs
			# plotn - pcutn - (l2fccutn - pl2fccutn) == nrow(na.omit(subset(res, !(res$padj < pcut) & !(res$log2FoldChange > l2fccut) & !(res$log2FoldChange < -l2fccut))))
			# totn - pcutn - (l2fccutn - pl2fccutn) == nrow(subset(res, !(res$padj < pcut) & !(res$log2FoldChange > l2fccut) & !(res$log2FoldChange < -l2fccut))) + nrow(subset(res, is.na(res$padj) | is.na(res$log2FoldChange)))


		# Plotting
			volcanoplot <- EnhancedVolcanoModified(
				toptable=res,
				# lab=as.character(rownames(res)),
				lab=volcano_gene_labs,
				x="log2FoldChange",
				y="padj",
				selectLab=select_genes,
				# selectLab=c("Lyz1"),
				#	selectLab=as.character(mappedtopten$"Gene name"),
				#selectLab=c("ENSMUSG00000001156", "ENSMUSG00000045502"),
				xlab=bquote(log[2]~fold~change),
				ylab=bquote(~-log[10]~adjusted~italic(p)),
				pCutoff=pcut,
				FCcutoff=l2fccut,
				#ylim=c(0,5),
				title=chart_title,
				# title="Alignment to mouse genome",
				# title=bquote(italic(S.)~italic(haematobium)~"(SchHae_2.0) genome alignment"),
				# title=bquote(italic(S)*italic(haematobium)*" genome: einj vs egg+eb"*),
				subtitle=paste0(cond_exp, "-vs-", cond_ctrl),
				# subtitle="einj vs egg+eb",
				caption = paste0('Total plotted = ', plotn, ' genes'),
				axisLabSize = 20,
				titleLabSize = 20,
				shape = c(1, 4, 23, 25),
				# legendLabels=c(
					# bquote(atop(
						# "NS",
						# .(plotn - pcutn - (l2fccutn - pl2fccutn))~"genes"
						# )),
					# bquote(atop(
						# italic(p)*"-adj >= "*.(pcut)*" and |"*log[2]*"FC| > "*.(l2fccut),
						# .(l2fccutn - pl2fccutn)~"genes"
						# )),
					# bquote(atop(
						# italic(p)*"-adj < "*.(pcut)*" and |"*log[2]*"FC| <= "*.(l2fccut),
						# .(pcutn - pl2fccutn)~"genes"
						# )),
					# bquote(atop(
						# italic(p)*"-adj < "*.(pcut)*" and |"*log[2]*"FC| > "*.(l2fccut),
						# .(pl2fccutn)~"genes"
						# ))
				legendLabels=c(
					bquote(atop(
						"NS",
						.(plotn - pcutn - (l2fccutn - pl2fccutn))~"genes"
						)),
					bquote(atop(
						italic(p)*"-adj \u2265 "*.(pcut)*" and |"*log[2]*"FC| > "*.(l2fccut),
						.(l2fccutn - pl2fccutn)~"genes"
						)),
					bquote(atop(
						italic(p)*"-adj < "*.(pcut)*" and |"*log[2]*"FC| \u2264 "*.(l2fccut),
						.(pcutn - pl2fccutn)~"genes"
						)),
					bquote(atop(
						italic(p)*"-adj < "*.(pcut)*" and |"*log[2]*"FC| > "*.(l2fccut),
						.(pl2fccutn)~"genes"
						))
					# sprintf("Adjusted p-value < %s and |Log2(FC)| > %s", pcut, l2fccut)
					),
				# labhjust = 1,
				labSize = 5,
				drawConnectors = TRUE
				) +
				theme(plot.title=element_text(face="plain"))
			png(paste0(pathout, "/", filenamestring, "-padj", myalpha, "-l2fc", l2fccut, igfilter_str, "-enhvolcano.png"), 1000, 1000)
			print(volcanoplot)
			dev.off()
		message("volcano plot done")

	# VennDiagram; only for mm
	if (analysis == "mm"){
		# Processing for microarray_file
			# Read file using read.table, then change to data.table (need data.table's unique(by=); base data.frame has unique() but not by=)
				library("data.table") # load the library
				microarray_file_dt <- as.data.table(read.table(file=microarray_file, header=TRUE, check.names=FALSE, sep="\t", quote="")) # read file into data.frame, then coerce to data.table
			# Need to convert line endings from \r\n to \n
				# Actually not needed?
			# When there exist multiple entries of same gene, select the first occurrence
				microarray_file_u_dt <- unique(microarray_file_dt, by="Gene Symbol")
			# Need to convert fold-change to log2foldchange
				microarray_fc <- microarray_file_u_dt$"Fold Change" # select the fold change values
				microarray_fcc <- ifelse(microarray_fc < 0, -1/microarray_fc, microarray_fc) # convert negative fold-change values into positive values between 0 and 1, e.g., -2 becomes 0.5
				microarray_l2fc <- log2(microarray_fcc) # convert fold-change values to log2foldchange values
				microarray_file_u_l2fc_df <- data.frame(
					"gene_symb" = microarray_file_u_dt$"Gene Symbol",
					"l2fc_microarraywk1" = microarray_l2fc
					) # select symbol and l2fc and create new data.frame
		# Processing for proteomics_file
			# Read file using read.excel
				library("readxl")
				proteomics_file_df <- read_xlsx(proteomics_file)
				proteomics_symbol <- sub( ".*GN=([^ ]+).*", "\\1", as.character(proteomics_file_df$"Description") ) # extract gene symbol from GN tag in "Description" (regex match entire line and replacement with captured group)
				proteomics_new_df <- data.frame(
					"uniprot_acc_proteomics" = proteomics_file_df$"Accession",
					"gene_symb" = proteomics_symbol,
					"l2fc_proteomics" = proteomics_file_df$"log2(FC)",
					"padj_proteomics" = proteomics_file_df$"p.ajusted"
					)
		# Processing for rnaseq deseq2 results; not igfiltered
			res_map_new <- data.frame(
				"ensembl_acc_rnaseq" = res_map$"Gene stable ID",
				"gene_symb" = res_map$"Gene name",
				"gene_desc" = res_map$"Gene description",
				"l2fc_rnaseq" = res_map$"log2FoldChange",
				"padj_rnaseq" = res_map$"padj"
				)
		# Use Reduce() and merge() to join rnaseq, microarray, and proteomics studies by gene symbol
			allstudies_df <- Reduce(
				function(x, y) merge(x, y, by="gene_symb", all=TRUE),
				list(
					res_map_new,
					microarray_file_u_l2fc_df,
					proteomics_new_df
					) )
		# Write combined data.frame to file
			write.table(allstudies_df, file=paste0(pathout, "/", filenamestring, "-diffexp-rnaseq-microarraywk1-prot.txt"), sep="\t", quote=F, row.names=F)
		# VennDiagram
			library("VennDiagram")
			# nrow(allstudies_df) # check total number of records
				# length(allstudies_df$"gene_symb")
			symb_rnaseq <- as.character(subset(allstudies_df, allstudies_df$"padj_rnaseq" < myalpha & abs(allstudies_df$"l2fc_rnaseq") > l2fccut)$"gene_symb")
				# length(symb_rnaseq)
			symb_rnaseq <- symb_rnaseq[ nzchar(symb_rnaseq) ] #in this case, vector does not contain empty string
				# length(symb_rnaseq)
			symb_m <- as.character(subset(allstudies_df, !is.na(allstudies_df$"l2fc_microarraywk1") )$"gene_symb")
				# length(symb_m)
			symb_m <- symb_m[ nzchar(symb_m) ]
				# length(symb_m)
			symb_p <- as.character(subset(allstudies_df, !is.na(allstudies_df$"l2fc_proteomics") )$"gene_symb")
				# length(symb_p)
			symb_p <- symb_p[ nzchar(symb_p) ]
				# length(symb_p)
			venn_list <- list("RNA-seq"=symb_rnaseq, "Microarray"=symb_m, "Proteomics"=symb_p)
			pf_venn <- paste0(pathout, "/", filenamestring, "-venn.tiff")
			venn.plot <- venn.diagram(
				x=venn_list,
				filename=pf_venn,
				resolution=600,
				scaled=FALSE,
				euler.d=FALSE,
				fontfamily="sans",
				cex=2.5,
				cat.fontfamily="sans",
				cat.cex=1)
		# Tests
			# check that microarraywk1 data: all records (260) satisfy |l2fc| > 1
				nrow(subset(allstudies_df, !is.na(allstudies_df$"l2fc_microarraywk1") )) ==
				nrow(subset(allstudies_df, abs(allstudies_df$"l2fc_microarraywk1") > 1))
			# check that proteomics data: all records (45) satisfy padj < 0.05
				nrow(subset(allstudies_df, !is.na(allstudies_df$"l2fc_proteomics") )) ==
				nrow(subset(allstudies_df, allstudies_df$"padj_proteomics" < 0.05 ))
				# note that all |l2fc_proteomics| < 1
					abs(subset(allstudies_df, !is.na(allstudies_df$"l2fc_proteomics") )$"l2fc_proteomics") < 1
			# confirm that there exist no records for which "padj_rnaseq" has a value but "l2fc_rnaseq" has NA
				# if true, this allows for us to check is.na for padj_rnaseq alone, without having to also check is.na for l2fc_rnaseq; in other words, if padj_rnaseq is NA, we can treat the record as NA for rnaseq without needing to check cases in which l2fc_rnaseq is NA but padj_rnaseq has a value
				nrow(subset(allstudies_df,
					!is.na(allstudies_df$"padj_rnaseq") &
					is.na(allstudies_df$"l2fc_rnaseq")
					))

		# Values in venn diagram, including total/agreeement/disagreement
			# Apply l2fc cutoff (l2fccut) to microarray and rnaseq but not to proteomics
			# Filter by condition; 15 total
				# 3 (differentially expressed genes/proteins unique to each study)
					# n_uniq_r, n_uniq_m, n_uniq_p
				# 3 overlap between rnaseq and microarray (total, agreements, disagreements)
					# n_rm, n_rm_a, n_rm_d
				# 3 overlap between rnaseq and protemoics
					# n_rp, n_rp_a, n_rp_d
				# 3 overlap between microarray and proteomics
					# n_mp, n_mp_a, n_mp_d
				# 3 overlap (all studies)
					# n_rmp, n_rmp_a, n_rmp_d
			# unique to rnaseq (|l2fc| > l2fccut and padj < myalpha)
				n_uniq_r <- nrow(subset(allstudies_df,
					abs(allstudies_df$"l2fc_rnaseq") > l2fccut &
					allstudies_df$"padj_rnaseq" < myalpha &
					is.na(allstudies_df$"l2fc_proteomics") &
					is.na(allstudies_df$"l2fc_microarraywk1")
					))
			# unique to microarray
				n_uniq_m <- nrow(subset(allstudies_df,
					is.na(allstudies_df$"l2fc_proteomics") &
					!is.na(allstudies_df$"l2fc_microarraywk1") &
					(
						is.na(allstudies_df$"l2fc_rnaseq") |
						is.na(allstudies_df$"padj_rnaseq") |
						abs(allstudies_df$"l2fc_rnaseq") <= l2fccut |
						allstudies_df$"padj_rnaseq" >= myalpha
						)
					))
			# unique to proteomics
				n_uniq_p <- nrow(subset(allstudies_df,
					!is.na(allstudies_df$"l2fc_proteomics") &
					is.na(allstudies_df$"l2fc_microarraywk1") &
					(
						is.na(allstudies_df$"l2fc_rnaseq") |
						is.na(allstudies_df$"padj_rnaseq") |
						abs(allstudies_df$"l2fc_rnaseq") <= l2fccut |
						allstudies_df$"padj_rnaseq" >= myalpha
						)
					))
			# overlap rnaseq and microarray (agreements+disagreements)
				n_rm <- nrow(subset(allstudies_df,
					is.na(allstudies_df$"l2fc_proteomics") &
					!is.na(allstudies_df$"l2fc_microarraywk1") &
					!is.na(allstudies_df$"l2fc_rnaseq") &
					!is.na(allstudies_df$"padj_rnaseq") &
					abs(allstudies_df$"l2fc_rnaseq") > l2fccut &
					allstudies_df$"padj_rnaseq" < myalpha
					))
			# overlap rnaseq and microarray (agreements)
				rm_a <- subset(allstudies_df,
					is.na(allstudies_df$"l2fc_proteomics") &
					!is.na(allstudies_df$"l2fc_microarraywk1") &
					!is.na(allstudies_df$"l2fc_rnaseq") &
					allstudies_df$"padj_rnaseq" < myalpha &
					(
						(allstudies_df$"l2fc_rnaseq" > l2fccut &
						allstudies_df$"l2fc_microarraywk1" > l2fccut) |
						(allstudies_df$"l2fc_rnaseq" < -l2fccut &
						allstudies_df$"l2fc_microarraywk1" < -l2fccut)
						)
					)
				n_rm_a <- nrow(rm_a)
			# overlap rnaseq and microarray (disagreements)
				n_rm_d <- nrow(subset(allstudies_df,
					is.na(allstudies_df$"l2fc_proteomics") &
					!is.na(allstudies_df$"l2fc_microarraywk1") &
					!is.na(allstudies_df$"l2fc_rnaseq") &
					allstudies_df$"padj_rnaseq" < myalpha &
					(
						(allstudies_df$"l2fc_rnaseq" > l2fccut &
						allstudies_df$"l2fc_microarraywk1" < -l2fccut) |
						(allstudies_df$"l2fc_rnaseq" < -l2fccut &
						allstudies_df$"l2fc_microarraywk1" > l2fccut)
						)
					))
			# overlap rnaseq and proteomics (agreements+disagreements)
				n_rp <- nrow(subset(allstudies_df,
					!is.na(allstudies_df$"l2fc_proteomics") &
					is.na(allstudies_df$"l2fc_microarraywk1") &
					!is.na(allstudies_df$"l2fc_rnaseq") &
					abs(allstudies_df$"l2fc_rnaseq") > l2fccut &
					allstudies_df$"padj_rnaseq" < myalpha
					))
			# overlap rnaseq and proteomics (agreements)
				rp_a <- subset(allstudies_df,
					!is.na(allstudies_df$"l2fc_proteomics") &
					is.na(allstudies_df$"l2fc_microarraywk1") &
					!is.na(allstudies_df$"l2fc_rnaseq") &
					abs(allstudies_df$"l2fc_rnaseq") > l2fccut &
					allstudies_df$"padj_rnaseq" < myalpha &
					(
						(allstudies_df$"l2fc_rnaseq" > l2fccut &
						allstudies_df$"l2fc_proteomics" > 0) |
						(allstudies_df$"l2fc_rnaseq" < -l2fccut &
						allstudies_df$"l2fc_proteomics" < 0)
						)
					)
				n_rp_a <- nrow(rp_a)
			# overlap rnaseq and proteomics (disagreements)
				n_rp_d <- nrow(subset(allstudies_df,
					!is.na(allstudies_df$"l2fc_proteomics") &
					is.na(allstudies_df$"l2fc_microarraywk1") &
					!is.na(allstudies_df$"l2fc_rnaseq") &
					abs(allstudies_df$"l2fc_rnaseq") > l2fccut &
					allstudies_df$"padj_rnaseq" < myalpha &
					(
						(allstudies_df$"l2fc_rnaseq" > l2fccut &
						allstudies_df$"l2fc_proteomics" < 0) |
						(allstudies_df$"l2fc_rnaseq" < -l2fccut &
						allstudies_df$"l2fc_proteomics" > 0)
						)
					))
			# overlap microarray and proteomics (agreements+disagreements)
				n_mp <- nrow(subset(allstudies_df,
					!is.na(allstudies_df$"l2fc_proteomics") &
					!is.na(allstudies_df$"l2fc_microarraywk1") &
					(
						is.na(allstudies_df$"l2fc_rnaseq") |
						is.na(allstudies_df$"padj_rnaseq") |
						abs(allstudies_df$"l2fc_rnaseq") <= l2fccut |
						allstudies_df$"padj_rnaseq" >= myalpha
						)
					))
			# overlap microarray and proteomics (agreements)
				mp_a <- subset(allstudies_df,
					!is.na(allstudies_df$"l2fc_proteomics") &
					!is.na(allstudies_df$"l2fc_microarraywk1") &
					(
						is.na(allstudies_df$"l2fc_rnaseq") |
						is.na(allstudies_df$"padj_rnaseq") |
						abs(allstudies_df$"l2fc_rnaseq") <= l2fccut |
						allstudies_df$"padj_rnaseq" >= myalpha
						) &
					(
						(allstudies_df$"l2fc_proteomics" > 0 &
							allstudies_df$"l2fc_microarraywk1" > 0
							) |
						(allstudies_df$"l2fc_proteomics" < 0 &
							allstudies_df$"l2fc_microarraywk1" < 0
							)
						)
					)
				n_mp_a <- nrow(mp_a)
			# overlap microarray and proteomics (disagreements)
				n_mp_d <- nrow(subset(allstudies_df,
					!is.na(allstudies_df$"l2fc_proteomics") &
					!is.na(allstudies_df$"l2fc_microarraywk1") &
					(
						is.na(allstudies_df$"l2fc_rnaseq") |
						is.na(allstudies_df$"padj_rnaseq") |
						abs(allstudies_df$"l2fc_rnaseq") <= l2fccut |
						allstudies_df$"padj_rnaseq" >= myalpha
						) &
					(
						(allstudies_df$"l2fc_proteomics" > 0 &
							allstudies_df$"l2fc_microarraywk1" < 0
							) |
						(allstudies_df$"l2fc_proteomics" < 0 &
							allstudies_df$"l2fc_microarraywk1" > 0
							)
						)
					))
			# overlap all studies (agreements+disagreements)
				n_rmp <- nrow(subset(allstudies_df,
					!is.na(allstudies_df$"l2fc_proteomics") &
					!is.na(allstudies_df$"l2fc_microarraywk1") &
					!is.na(allstudies_df$"l2fc_rnaseq") &
					abs(allstudies_df$"l2fc_rnaseq") > l2fccut &
					allstudies_df$"padj_rnaseq" < myalpha
					))
			# overlap all studies (agreements)
				rmp_a <- subset(allstudies_df,
					!is.na(allstudies_df$"l2fc_proteomics") &
					!is.na(allstudies_df$"l2fc_microarraywk1") &
					!is.na(allstudies_df$"l2fc_rnaseq") &
					allstudies_df$"padj_rnaseq" < myalpha &
					(
						(allstudies_df$"l2fc_proteomics" > 0 &
							allstudies_df$"l2fc_microarraywk1" > l2fccut &
							allstudies_df$"l2fc_rnaseq" > l2fccut
							) |
						(allstudies_df$"l2fc_proteomics" < 0 &
							allstudies_df$"l2fc_microarraywk1" < -l2fccut &
							allstudies_df$"l2fc_rnaseq" < -l2fccut
							)
						)
					)
				n_rmp_a <- nrow(rmp_a)
			# overlap all studies (disagreements)
				n_rmp_d <- nrow(subset(allstudies_df,
					!is.na(allstudies_df$"l2fc_proteomics") &
					!is.na(allstudies_df$"l2fc_microarraywk1") &
					!is.na(allstudies_df$"l2fc_rnaseq") &
					allstudies_df$"padj_rnaseq" < myalpha &
					abs(allstudies_df$"l2fc_microarray") > l2fccut &
					abs(allstudies_df$"l2fc_rnaseq") > l2fccut &
					!(
						(allstudies_df$"l2fc_proteomics" > 0 &
							allstudies_df$"l2fc_microarraywk1" > l2fccut &
							allstudies_df$"l2fc_rnaseq" > l2fccut
							) |
						(allstudies_df$"l2fc_proteomics" < 0 &
							allstudies_df$"l2fc_microarraywk1" < -l2fccut &
							allstudies_df$"l2fc_rnaseq" < -l2fccut
							)
						)
					))
		# Write agreements to file
			write.table(rm_a, file=paste0(pathout, "/", filenamestring, "-diffexp-venn-rm-a.txt"), sep="\t", quote=F, row.names=F)
			write.table(rp_a, file=paste0(pathout, "/", filenamestring, "-diffexp-venn-rp-a.txt"), sep="\t", quote=F, row.names=F)
			write.table(mp_a, file=paste0(pathout, "/", filenamestring, "-diffexp-venn-mp-a.txt"), sep="\t", quote=F, row.names=F)
			write.table(rmp_a, file=paste0(pathout, "/", filenamestring, "-diffexp-venn-rmp-a.txt"), sep="\t", quote=F, row.names=F)
		# Write venn diagram values to file
			venn_num_fx <- file(paste0(pathout, "/", filenamestring, "-venn_numbers.txt"))
			writeLines(
				text=c(
					"Venn diagram numbers",
					paste0("RNA-seq conditions: p-adj<", myalpha, ", |l2fc|>", l2fccut),
					paste0("Microarray (week 1) conditions: |l2fc|>1 (already satisfied) "),
					paste0("Proteomics conditions: p-adj<0.05 (already satisfied), l2fc cutoff not applied"),
					"",
					paste0("Unique to RNA-seq, microarray, proteomics"),
					paste(n_uniq_r, n_uniq_m, n_uniq_p, sep=" "),
					"Overlap RNA-seq and microarray (total, agreements, disagreements)",
					paste(n_rm, n_rm_a, n_rm_d, sep=" "),
					"Overlap RNA-seq and proteomics (total, agreements, disagreements)",
					paste(n_rp, n_rp_a, n_rp_d, sep=" "),
					"Overlap microarray and proteomics (total, agreements, disagreements)",
					paste(n_mp, n_mp_a, n_mp_d, sep=" "),
					"Overlap all studies (total, agreements, disagreements)",
					paste(n_rmp, n_rmp_a, n_rmp_d, sep=" ")
					),
				con=venn_num_fx
				)
			close(venn_num_fx)
		}

	message("venn diagram done")
	message(paste0("Analysis done: ", analysis, " with ", anum, "-deseq2-", omit_samples, "-", pool_eggeb, "-includefirst", includefirst, "-shref", shref))

	}

# Feed mainfunction() with analysis parameters
	script_ver <- 8
	iter <- 1
	# Explanation of inputs
		# pathproj: character string containing path to project directory
		# analysis: character string "mm" or "sh" indicating mouse or S. haematobium analysis
			# "mm": also applies filtering for immunoglobulin genes other than Igha, Ighd, Ighe, Ighg, Ighm
		# omit_samples: character string "omitn", "omity", or "exclusive"
			# "omitn": don't omit any samples
			# "omity": omit einj samples that failed to receive eggs (22x and 321)
			# "exclusive": analysis-dependent sample removal (also removes 22x and 321 as in "omity")
				# if analysis is "mm", remove naive (x0x), egg (x3x), eb (x4x) such that hinj (x1x), einj (x2x) remain
				# if analysis is "sh", remove naive (x0x), hinj (x1x) such that einj (x2x), egg (x3x), eb (x4x) remain
		# pool_eggeb: character string "pooln" or "pooly"
		# anum: character string that incorporates script version and iteration number
		# includefirst: logical value that, if true, causes the analysis to include run 1 data
		# shref: logical value that, if true, causes the analysis to include sh reference data

	# mm omitn pooln
		# discover samples to omit (22x and 321)
		# enable point labeling on PCA
		# only interested in PCA for this analysis
		anum <- sprintf("%s%02d-%02d", "v", script_ver, iter)
		mainfunction(pathproj=pathproj, analysis="mm", omit_samples="omitn", pool_eggeb="pooln", anum=anum, includefirst=TRUE, shref=FALSE)
		# iter <- iter + 1

	# mm omity pooln
		# we know egg and eb look different for mm, so don't pool them
		# only interested in PCA for this analysis
		iter <- 2
		anum <- sprintf("%s%02d-%02d", "v", script_ver, iter)
		mainfunction(pathproj=pathproj, analysis="mm", omit_samples="omity", pool_eggeb="pooln", anum=anum, includefirst=FALSE, shref=FALSE)
		# iter <- iter + 1

	# mm exclusive pooln
		# we know egg and eb look different for mm, so don't pool them
		# einj vs hinj exclusive analysis (omit other samples)
		iter <- 3
		anum <- sprintf("%s%02d-%02d", "v", script_ver, iter)
		mainfunction(pathproj=pathproj, analysis="mm", omit_samples="exclusive", pool_eggeb="pooln", anum=anum, includefirst=FALSE, shref=FALSE)
		# iter <- iter + 1

	# sh omity pooln
		# discover that for sh: egg, eb, R31 look the same
		iter <- 4
		anum <- sprintf("%s%02d-%02d", "v", script_ver, iter)
		mainfunction(pathproj=pathproj, analysis="sh", omit_samples="omity", pool_eggeb="pooln", anum=anum, includefirst=FALSE, shref=TRUE)
		# iter <- iter + 1

	# sh omity pooly
		# pool egg, eb, R31 into egg group
		iter <- 5
		anum <- sprintf("%s%02d-%02d", "v", script_ver, iter)
		mainfunction(pathproj=pathproj, analysis="sh", omit_samples="omity", pool_eggeb="pooly", anum=anum, includefirst=FALSE, shref=TRUE)
		# iter <- iter + 1

	# sh exclusive pooly
		# pool egg, eb, R31 into egg group
		# einj vs (egg+eb+R31) exclusive analysis (omit other samples)
		iter <- 6
		anum <- sprintf("%s%02d-%02d", "v", script_ver, iter)
		mainfunction(pathproj=pathproj, analysis="sh", omit_samples="exclusive", pool_eggeb="pooly", anum=anum, includefirst=FALSE, shref=TRUE)
		# iter <- iter + 1

	# sh exclusive pooly
		# don't include ref data R31
		# pool egg, eb, R31 into egg group
		# einj vs (egg+eb+R31) exclusive analysis (omit other samples)
		iter <- 7
		anum <- sprintf("%s%02d-%02d", "v", script_ver, iter)
		mainfunction(pathproj=pathproj, analysis="sh", omit_samples="exclusive", pool_eggeb="pooly", anum=anum, includefirst=FALSE, shref=FALSE)
		# iter <- iter + 1
```
