## Supplementary 2 for "RNA-seq gene expression profiling of the bladder in a mouse model of urogenital schistosomiasis": 05_gsea_master.sh

```
# What this script does
  # Executes GSEA (version 4.2.3)

# How to use this script
  # Save a copy
  # Customize path/file variables (replace empty strings ""), other settings
  # Save
  # Feed script to bash (e.g., bash customizedscript.sh)

# Software and resources
  # Official repository for GSEA executable
    # https://www.gsea-msigdb.org/gsea/index.jsp
  # Gene set collections (.gmx/.gmt)
    # ftp://ftp.broadinstitute.org/distribution/gsea/msigdb/mouse/gene_sets/
  # Gene mapping/annotation files (.chip)
    # ftp://ftp.broadinstitute.org/distribution/gsea/msigdb/mouse/annotations/

# Manual
  # https://software.broadinstitute.org/cancer/software/gsea/wiki/index.php/Main_Page
  # https://www.gsea-msigdb.org/gsea/doc/GSEAUserGuideFrame.html

# Paths
  # GSEA executable
    path_gsea=""
  # Output
    path_out=""

# Input files (full path and file)
  # Normalized gene counts from e.g. DESeq2; see manual for proper format
    pf_normcounts=""
  # Class (.cls) file that associates each sample to a treatment group
    pf_cls=""
  # Gene set collection (.gmx) file that associates genes to gene sets
    pf_gmx=""
  # Gene mapping (.chip) file that associates ENSEMBL gene accessions to gene symbols and descriptions
    pf_chip=""

# Other settings
  # Reference treatment group
    treat_ref="hinj"
  # Experiment treatment group
    treat_exp="einj"
  # Report name
    name_report=""
  # Analysis timestamp
    mytime=$(date +%s)

# Execute GSEA
  "${path_gsea}/gsea-cli.sh" \
    GSEA \
    -res "${pf_normcounts}" \
    -cls "${pf_cls}#${treat_exp}_versus_${treat_ref}" \
    -gmx "${pf_gmx}" \
    -collapse Collapse \
    -mode Max_probe \
    -norm meandiv \
    -nperm 1000 \
    -permute gene_set \
    -rnd_type no_balance \
    -scoring_scheme weighted \
    -rpt_label "${name_report}" \
    -metric Signal2Noise \
    -sort real \
    -order descending \
    -chip "${pf_chip}" \
    -create_gcts true \
    -create_svgs false \
    -include_only_symbols true \
    -make_sets true \
    -median false \
    -num 100 \
    -plot_top_x 20 \
    -rnd_seed timestamp \
    -save_rnd_lists false \
    -set_max 500 \
    -set_min 15 \
    -zip_report false \
    -out "${path_out}" \
    2>&1 | tee "${path_out}/stdouterr_${mytime}.log"
```
