## Supplementary 2 for "RNA-seq gene expression profiling of the bladder in a mouse model of urogenital schistosomiasis": 06_gsea_rpt_cleanup_master.sh

```
# What this script does
  # GSEA report table cleanup

pf_up="" # path-file GSEA pos enriched (tsv report file)
pf_dn="" # path-file GSEA neg enriched (tsv report file)
pf_out="" # path-file for writing output

awkcmd_head='BEGIN{FS=OFS="\t"}; NR==1 {print $1, $4, $5, $6, $7, $8, $9, $10, $11}'
awkcmd_body='BEGIN{FS=OFS="\t"}; NR>1 && $8<0.05 {printf "%s\t%s\t%3.2f\t%3.2f\t%3.2e\t%3.2e\t%3.2e\t%s\t%s\n", $1, $4, $5, $6, $7, $8, $9, $10, $11}'

awk "$awkcmd_head" "$pf_up" > "$pf_out"
awk "$awkcmd_body" "$pf_up" \
  | sort -t $'\t' -gk 3,3 -r \
  >> "$pf_out"
awk "$awkcmd_body" "$pf_dn" \
  | sort -t $'\t' -gk 3,3 -r \
  >> "$pf_out"

# Subset of columns
# echo $'Name\tSize\tNES\tFDR q-val\tRank at max\tLeading edge' > "$pf_out"
# awk \
  # 'BEGIN{FS=OFS="\t"}
  # NR>1 && $8<0.05 {printf "%s\t%s\t%3.2f\t%3.2e\t%s\t%s\n", $1, $4, $6, $8, $10, $11}' "$pf_up" \
  # | sort -t $'\t' -gk 3,3 -r \
  # >> "$pf_out"
# awk \
  # 'BEGIN{FS=OFS="\t"}
  # NR>1 && $8<0.05 {printf "%s\t%s\t%3.2f\t%3.2e\t%s\t%s\n", $1, $4, $6, $8, $10, $11}' "$pf_dn" \
  # | sort -t $'\t' -gk 3,3 -r \
  # >> "$pf_out"
```
