## Supplementary 2 for "RNA-seq gene expression profiling of the bladder in a mouse model of urogenital schistosomiasis": 07_panther_plot_master.R

```
# Load libraries
	library("ggplot2")
	library("ggrepel")

prep_df_func <- function(pf_panther, plot_label_vec, fdr_cut, fc_cut)
	{
	# Read table into R and clean up
	if (TRUE)
		{
		# Output file from PANTHER has 5 blank lines, followed by 6 lines of analysis parameters, a line of column names, then data
		# The number of fields (columns) at beginning of the file does not match throughout
		# For correct reading, need to specify maximum number of columns that exists in the file (expect 8)
			panther_df_colmax <- max(count.fields(pf_panther, sep = "\t"))
		# Provide this number of columns to read.table()
			panther_df <-
				read.table(
					file = pf_panther,
					header = FALSE,
					sep = "\t",
					quote = "",
					row.names = NULL,
					col.names = paste0("V", seq_len(panther_df_colmax)),
					check.names = FALSE,
					fill = TRUE,
					blank.lines.skip = TRUE,
					stringsAsFactors = FALSE
					)
		# Remove columns other than name, fold enrichment, and false discovery rate
			panther_df <- panther_df[ ,-c(2, 3, 4, 5, 7)]
		# Remove first 7 rows (analysis parameters and column names row)
			panther_df <- panther_df[-(1:7), ]
		# Set column names
			colnames(panther_df) <- c("name", "fe", "fdr")
		# Remove gene ontology number (GO:xxxxxxx) from names
			panther_df$"name" <- sub(" \\(.*\\)$", "", panther_df$"name")
		# Change fold enrichment and false discovery rate from string to number
			panther_df$"fe" <- as.numeric(panther_df$"fe") # some have value "< 0.01", which R will coerce to NA
			panther_df$"fdr" <- as.numeric(panther_df$"fdr")
		}
	# Get index values (row numbers) for rows whose name matches those specified for labeling
		idx <- which(panther_df$"name" %in% plot_label_vec)
	# Create a new column for labels and populate only the records to label
		panther_df$"label" <- NA
		panther_df[idx, ]$"label" <- panther_df[idx, ]$"name"
	# Subset the data by the cutoffs
		panther_df_cut <- subset(panther_df, panther_df$"fdr" < fdr_cut & abs(panther_df$"fe") > fc_cut)
	# Return cleaned data.frame
		return(panther_df_cut)
	}

if (TRUE) # User input
	{
	# Specify full path/file of output from the PANTHER Overrepresentation Test
		pf_panther_bp <- ""
		# pf_panther_pathway <- ""
	# Specify analysis
		analysis <- "upreg" # choose: "upreg", "dnreg"
	# Specify analysis category (string used in filename)
		category <- "bp" # examples: "bp", "cc", "mf"
	# Specify path for writing files
		path_out <- ""
	# Specify plot labels for biological process (bp)
		plot_label_bp_vec <- c(
			"defense response",
			"response to other organism",
			"response to cytokine",
			"immune system process",
			"immune response",
			"cytokine-mediated signaling pathway",
			"inflammatory response",
			"response to interferon-beta",
			"leukocyte chemotaxis"
			)
	# Specify plot labels for pathway
		plot_label_pathway_vec <- c(
			"Interleukin signaling pathway",
			"Inflammation mediated by chemokine and cytokine signaling pathway"
			)
	# Specify fdr cutoff
		fdr_cut <- 1e-5
	# Specify fold change cutoff
		fc_cut <- 2
	}

# Get data.frames and combine (or not)
	bp_df <- prep_df_func(pf_panther_bp, plot_label_bp_vec, fdr_cut, fc_cut)
	# pathway_df <- prep_df_func(pf_panther_pathway, plot_label_pathway_vec, fdr_cut, fc_cut)
	# panther_df_cut <- rbind(bp_df, pathway_df)
	panther_df_cut <- bp_df
# Set the point color of selected/labeled
	if (analysis == "upreg"){
		point_color <- "red"
		plot_title_str <- "upregulates"
		}
	if (analysis == "dnreg"){
		point_color <- "blue"
		plot_title_str <- "downregulates"
		}
# Set seed for recreating output
	set.seed(42)
# Plot
	panther_plot <-
		ggplot(
			data = panther_df_cut,
			aes(
				x = log2(fe),
				y = -log10(fdr)
				)
			) +
		labs(
			x = bquote(log[2]~fold~enrichment),
			y = bquote(-log[10]~adjusted~italic(p))
			# title = paste(strwrap(paste0("The infection ", plot_title_str, " the expression of genes associated with the following biological processes"), width = 70), collapse = "\n")
			) +
		scale_x_continuous(expand = c(0.01,0)) +
		scale_y_continuous(expand = c(0.01,0)) +
		geom_point(
			data = subset(panther_df_cut, is.na(panther_df_cut$"label")),
			color = "grey50",
			na.rm = TRUE
			) +
		geom_point(
			data = subset(panther_df_cut, !is.na(panther_df_cut$"label")),
			color = point_color,
			na.rm = TRUE
			) +
		geom_label_repel(
			data = panther_df_cut,
			box.padding = 1,
			aes(
				label = panther_df_cut$"label",
				size = 10
				),
			na.rm = TRUE
			) +
		theme_bw() +
		theme(
			panel.border = element_blank(),
			title = element_text(size=20),
			axis.line = element_line(
				size = 0.8,
				color = "black"
				),
			axis.text = element_text(size=20),
			axis.title = element_text(size=20),
			# legend.text = element_text(size=20),
			# legend.title = element_text(size=20),
			legend.position = "none",
			aspect.ratio = 1
			)
	# tiff(file.path(path_out, "pantherplot.tiff"), width=1000, height=1000)
	png(paste0(path_out, "/pantherplot_", analysis, "_", category, ".png"), width=1000, height=1000)
	print(panther_plot)
	dev.off()
```
