## Supplementary 2 for "RNA-seq gene expression profiling of the bladder in a mouse model of urogenital schistosomiasis": 08_ipa_dot_plot_master.R

```
# Load libraries
	library("ggplot2")
	library("readxl")
# Full path and filename for exported IPA pathways file
	input_file <- ""
# Path for writing output
	path_out <- ""
# Expected file format
	# First row: IPA info line
	# Second row: column names
	# Third row and beyond: data
# Expected sort order
	# Largest-to-smallest "-log(p-value)"
# Read file, trim, set colnames
	ipa_tbl <- read_excel(path = input_file, col_names = FALSE)
	colnames(ipa_tbl) <- ipa_tbl[2, ]
	ipa_tbl <- ipa_tbl[-(1:2), ]
# colnames(ipa_tbl) expected
	# "Ingenuity Canonical Pathways"
	# "-log(p-value)"
	# "Ratio"
	# "z-score"
	# "Molecules"
	# "group" (added below)

# Create z-score groups
	ipa_tbl[which(ipa_tbl['z-score'] == 0.00),"group"] <- "No Pattern"
	ipa_tbl[which(ipa_tbl['z-score'] < 0.00),"group"] <- "Negative z-score"
	ipa_tbl[which(ipa_tbl['z-score'] > 0.00),"group"] <- "Positive z-score"

# Retain only records for which z-score value exists
	ipa_tbl <- subset(ipa_tbl, !is.na(ipa_tbl$"z-score"))
# Make these columns numeric (convert string to numeric)
	ipa_tbl$"-log(p-value)" <- as.numeric(ipa_tbl$"-log(p-value)")
	ipa_tbl$"z-score" <- as.numeric(ipa_tbl$"z-score")
# Select top 50 by largest "-log(p-value)" (assumes already in descending sort order)
	ipa_tbl_nocut <- ipa_tbl
	ipa_tbl <- ipa_tbl_nocut[1:50, ]
# Factorize pathways names and specify the levels in the order corresponding to largest-to-smallest "-log(p-value)"
	ipa_tbl$"Ingenuity Canonical Pathways" <-
		factor(ipa_tbl$"Ingenuity Canonical Pathways", levels = ipa_tbl$"Ingenuity Canonical Pathways"[order(ipa_tbl$"-log(p-value)", decreasing = FALSE)])
# Dot plot
	ipa_tbl_dot <-
		ggplot(
			data = ipa_tbl,
			aes(x = ipa_tbl$"-log(p-value)",
				y = ipa_tbl$"Ingenuity Canonical Pathways",
				size = abs(ipa_tbl$"z-score"),
				color = group)
			) + 
		geom_point(alpha = 3, na.rm = TRUE) + 
		theme_classic() +
		#scale_color_manual(values = c("No Pattern" = "gray", "Negative z-score" = "blue", "Positive z-score" = "red")) +
		scale_color_manual(values = c("blue", "grey", "red")) +
		guides(color = guide_legend(title = "z-score direction")) +
		guides(size = guide_legend(title = "z-score magnitude")) +
		# zscore range for color
		#scale_color_gradient(low = "blue", high = "red", limit = c(-4.2,4), name = "z-score")+
		# Ratio range for size
		xlab("-log(p-value)") +
		ylab("Signaling Pathways") +
		# ggtitle("Perturbed Pathways (IL27+ vs Neg (Signaling)") +
		theme(plot.title = element_text(size = 10),
			# axis.text.x = element_text(size = 8, angle = 90),
			legend.title = element_text(size = 10), 
			legend.text = element_text(size = 8),
			panel.border = element_rect(colour = "grey", fill = NA, size = 1),
			panel.grid.major = element_line(size = (0.2), colour = "grey"))
	png(file.path(path_out, "ipa_dot_plot_signalingpathways.png"), width = 1000, height = 1000)
	print(ipa_tbl_dot)
	dev.off()
```
