## Supplementary 2 for "RNA-seq gene expression profiling of the bladder in a mouse model of urogenital schistosomiasis": 09_extract_exon_idproduct_map_sh_master.sh

```
# Extract gene_id and product values for each exon record in shv2 gtf file

pf_gtf="" # path-file for shv2 gtf GCF_000699445.2_SchHae_2.0_genomic.gtf
pf_out="" # path-file for writing output
grep -v "^#" "$pf_gtf" \
  | awk -F "\t" '$3=="exon"{print $9}' \
  \
  \
  | head \
  \
  \
  | awk \
    'BEGIN{FS=OFS="; "}
    {
      j=1; for(i=1;i<=NF;i++){
        match($i, /(gene_id|product) \"([^\";]+)\"/, arr);
        if(arr[0]!=""){
          printf("%s%s", arr[2], (j==1 ? "\t" : "\n") ); j+=1
          }
        }
      }' \
  | uniq \
  > "$pf_out"

# Warnings
  # both .gene_id. key and .product. key must exist
  # .gene_id. key must occur before .product. key
  # values for these keys must exist
  # arr[0]!="" only catches non-matches; any unexpected cases such as product key occurs before gene_id tag will result in printing of the product value first
  # also will fail to (correctly) handle unexpected cases that multiple gene_id and/or product key occur on one line (will print the first two results with tab separation, then will proceed to line-separate subsequent results from that line
```
