## Supplementary 3 for "RNA-seq gene expression profiling of the bladder in a mouse model of urogenital schistosomiasis"

**Detailed methods for bioinformatics analysis**

Files containing RNA-sequencing reads were obtained from BGI and processed through several different software programs, running on a workstation equipped with dual Intel Xeon E5-2630 v4 CPUs and 512 GB RAM, under a CentOS Linux 7 x86_64-based operating system and the miniconda2 package and environment management system (version 4.8.3). The files were checked with FastQC (version 0.11.9 [1], from the conda channel bioconda). Alignment to the host mouse genome was performed with STAR (version 2.7.3a [2]) and mouse genome file (version GRCm38.p6 Ensembl release 98 obtained from [3]). Annotation and counting of the mapped reads was performed with featureCounts of the Subread software (version 2.0.0 [4]) using a mouse gene annotation file (version GRCm38.p6 Ensembl release 98 obtained from [5]), selecting "exon" as the feature type, selecting "gene_id" as the gene identifier field, and specifying paired-end mode. Read count normalization, principal component analysis, and differential gene expression analysis were performed with DESeq2 (bioconductor-deseq2, version 1.2.6 [6], build r36he1b5a44_0, from the conda channel bioconda) running under the R software environment (version 3.6.1 [7], build h3a67422_6, from the conda channel conda-forge) using an R script modified from a freely available template [8]. Gene identifier mappings were obtained from the BioMart interface on the Ensembl website [9], selecting the database "Ensembl Genes 98", dataset "Mouse genes (GRCm38.p6)", and the attributes "Gene stable ID", "Gene description", and "Gene name". Results of principal component analyses (PCA) were plotted using ggplot2 (version 3.2.1 [10]) and ggrepel (version 0.8.1 [11]) libraries for R. Volcano plots were generated using EnhancedVolcano (bioconductor-enhancedvolcano, version 1.4.0 [12], build r36_0, from the conda channel bioconda). Command-line utility software (such as awk, grep, head, sed, sort, tail, uniq, and wc) and Excel spreadsheet software (Microsoft) were used to parse gene annotation files, map gene accession identifiers to gene names and descriptions, and prepare and arrange gene (and gene set) lists and tables. Notepad++ software (version 7.8.2 [13]) was used to draft scripts that specify the parameters for analysis.

To associate differentially expressed genes with known signaling pathways and biological phenotypes, the normalized counts file generated by DESeq2 was used as an input file to gene set enrichment analysis (GSEA) software (version 4.2.3 [14]), along with a mapping file containing Ensembl gene accession identifiers and symbols [15] and mouse gene set collection files for hallmark gene sets ("MH" [16]) from the molecular signatures database (MSigDB [17]) and a sample class descriptor file (.cls), with the software parameters "permutation type" set to "gene_set" and "create_gcts" set to "true". Heat maps were generated using the Morpheus matrix visualization and analysis software [18].

Phenotype association analysis was also performed with Ingenuity Pathway Analysis (IPA, Qiagen). A file containing Ensembl accession identifiers, $\log_2$ fold change, and adjusted $p$ values for differentially expressed genes identified by DESeq2 (alpha = 0.05) satisfying the conditions $p$-adj < 0.05 and $|\log_2 FC| > 1$, filtered such that genes related to immunoglobulin variable regions were removed, was used as input for IPA analysis. Pathways and their associated z-scores and -$\log_{10}$ $p$-values from the "Signaling Pathways" analysis were visualized as a dot plot using ggplot2.

For gene ontology analysis, differentially expressed genes identified by DESeq2 (alpha = 0.05) satisfying the conditions $p$-adj < 0.05 and $|\log_2 FC| > 1$ were filtered such that genes related to immunoglobulin variable regions were removed. The resulting data were separated into lists of upregulated and downregulated genes, which were used as input for the PANTHER Overrepresentation Test (version 18.0, released 2023-10-17 [19,20]), with annotations from the Gene Ontology Data Archive (version 2024-01-17 [21]) for various annotation data sets, selecting *Mus musculus* as the target and reference organism. The fold enrichment and false discovery rate values were plotted using ggplot2 for the gene ontology for biological process. For the comparison of differentially expressed genes with those found in previous studies, genes were matched by gene symbol, which were used as input for the PANTHER Overrepresentation Test as described above. Venn diagram visualization was produced with VennDiagram (r-venndiagram, version 1.6.20 [22], build r36h6115d3f_1002, from the conda channel conda-forge) and GIMP (version 2.10.18 [23]).

Sequencing read alignment to the *S. haematobium* genome, read counting, normalization, principal component analysis, and differential gene expression analysis were done similarly as described above, using SchHae_2.0 genome [24] and gene annotation [25] files available from the National Center for Biotechnology Information (NCBI). For the read alignment of samples from the hamster bladder wall injection, the *Mesocricetus auratus* MesAur1.0 genome [26] and gene annotation [27] files from the Ensembl database (release 104) were used.

22. Chen H, Boutros P. VennDiagram: Generate High-Resolution Venn and Euler Plots. Available: https://cran.r-project.org/package=VennDiagram

23. GNU Image Manipulation Program (GIMP). Available: https://www.gimp.org/

24. NCBI Schistosoma haematobium assembly SchHae_2.0 genome. Available: ftp://ftp.ncbi.nlm.nih.gov/genomes/all/GCF/000/699/445/GCF_000699445.2_SchHae_2.0/GCF_000699445.2_SchHae_2.0_genomic.fna.gz

25. NCBI Schistosoma haematobium assembly SchHae_2.0 gene annotation. Available: ftp://ftp.ncbi.nlm.nih.gov/genomes/all/GCF/000/699/445/GCF_000699445.2_SchHae_2.0/GCF_000699445.2_SchHae_2.0_genomic.gtf.gz

26. Ensembl 104 MesAur1.0 hamster genome (toplevel). Available: ftp://ftp.ensembl.org/pub/release-104/fasta/mesocricetus_auratus/dna/Mesocricetus_auratus.MesAur1.0.dna.toplevel.fa.gz

27. Ensembl 104 MesAur1.0 hamster gene annotations. Available:

ftp://ftp.ensembl.org/pub/release-104/gtf/mesocricetus_auratus/Mesocricetus_auratus.MesAur1.0.104.gtf.gz
