## Supplementary figures and images for "RNA-seq gene expression profiling of the bladder in a mouse model of urogenital schistosomiasis"

### Supplementary 6

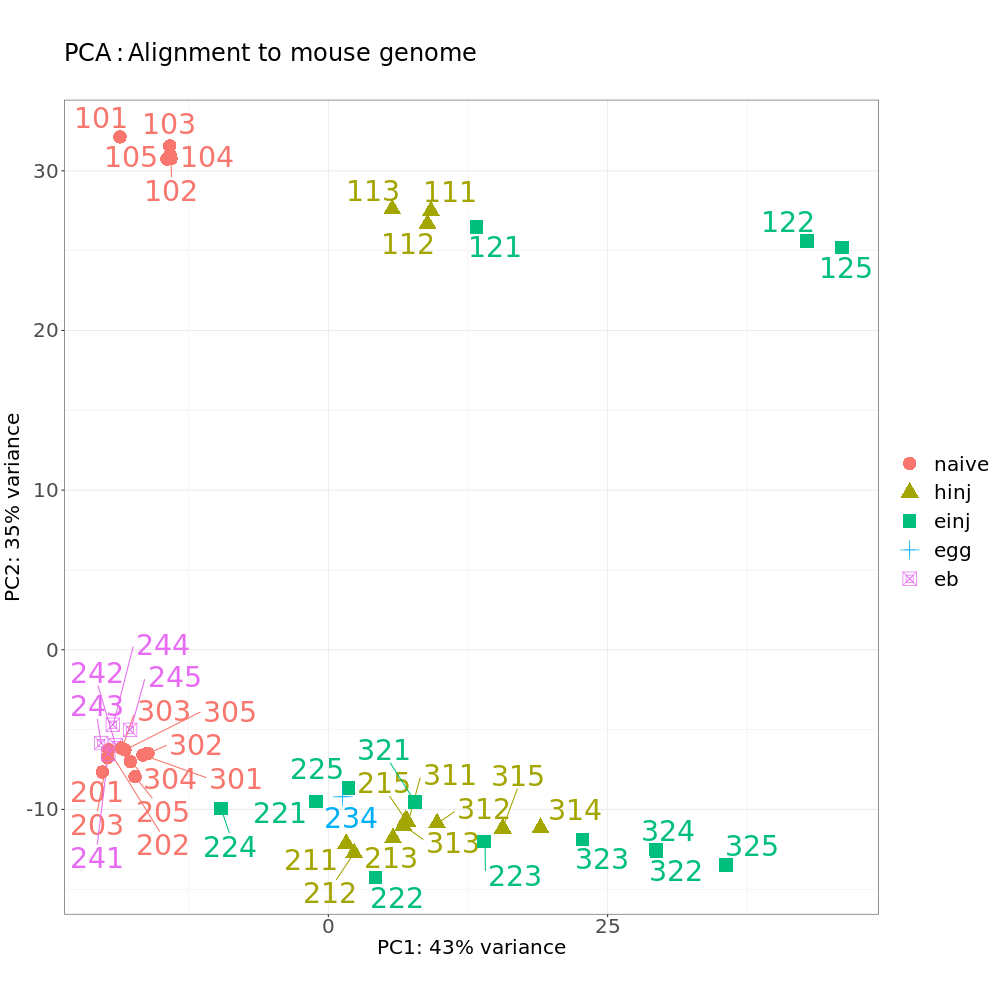

### Supplementary 13

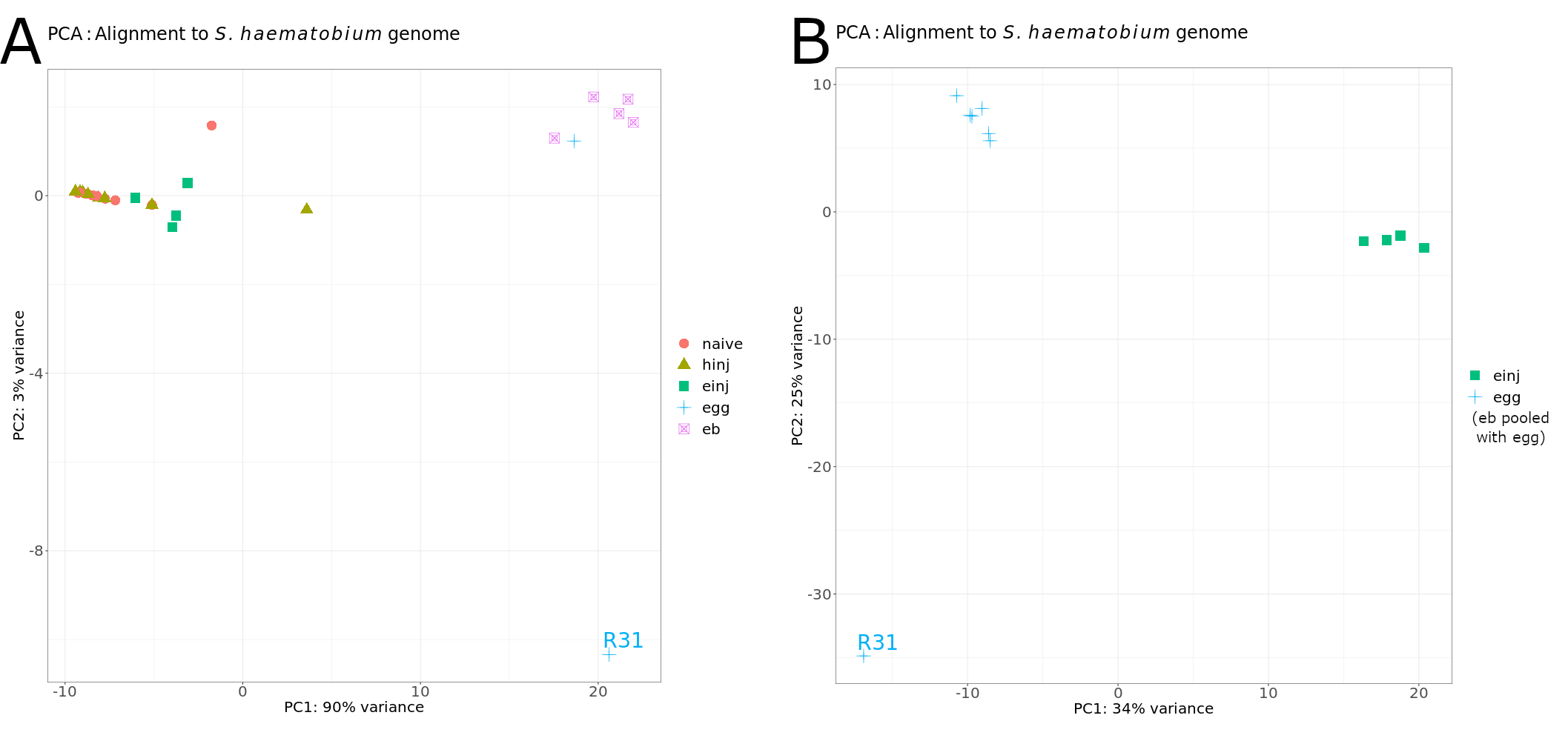
